# Disentangling Sources of Gene Tree Discordance in Phylogenomic Datasets: Testing Ancient Hybridizations in Amaranthaceae s.l

**DOI:** 10.1101/794370

**Authors:** Diego F. Morales-Briones, Gudrun Kadereit, Delphine T. Tefarikis, Michael J. Moore, Stephen A. Smith, Samuel F. Brockington, Alfonso Timoneda, Won C. Yim, John C. Cushman, Ya Yang

**Affiliations:** Department of Plant and Microbial Biology, University of Minnesota-Twin Cities, 1445 Gortner Avenue, St. Paul, MN 55108, USA; Institut für Molekulare Physiologie, Johannes Gutenberg-Universität Mainz, D-55099, Mainz, Germany; Department of Biology, Oberlin College, Science Center K111, 119 Woodland Street, Oberlin, OH 44074-1097, USA; Department of Ecology & Evolutionary Biology, University of Michigan, 830 North University Avenue, Ann Arbor, MI 48109-1048, USA; Department of Plant Sciences, University of Cambridge, Tennis Court Road, Cambridge, CB2 3EA, United Kingdom; Department of Biochemistry and Molecular Biology, University of Nevada, Reno, NV, 89577, USA

**Keywords:** Amaranthaceae, gene tree discordance, hybridization, incomplete lineage sorting, phylogenomics, transcriptomics, species tree, species network

## Abstract

Gene tree discordance in large genomic datasets can be caused by evolutionary processes such as incomplete lineage sorting and hybridization, as well as model violation, and errors in data processing, orthology inference, and gene tree estimation. Species tree methods that identify and accommodate all sources of conflict are not available, but a combination of multiple approaches can help tease apart alternative sources of conflict. Here, using a phylotranscriptomic analysis in combination with reference genomes, we test a hypothesis of ancient hybridization events within the plant family Amaranthaceae s.l. that was previously supported by morphological, ecological, and Sanger-based molecular data. The dataset included seven genomes and 88 transcriptomes, 17 generated for this study. We examined gene-tree discordance using coalescent-based species trees and network inference, gene tree discordance analyses, site pattern tests of introgression, topology tests, synteny analyses, and simulations. We found that a combination of processes might have generated the high levels of gene tree discordance in the backbone of Amaranthaceae s.l. Furthermore, we found evidence that three consecutive short internal branches produce anomalous trees contributing to the discordance. Overall, our results suggest that Amaranthaceae s.l. might be a product of an ancient and rapid lineage diversification, and remains, and probably will remain, unresolved. This work highlights the potential problems of identifiability associated with the sources of gene tree discordance including, in particular, phylogenetic network methods. Our results also demonstrate the importance of thoroughly testing for multiple sources of conflict in phylogenomic analyses, especially in the context of ancient, rapid radiations. We provide several recommendations for exploring conflicting signals in such situations.

---

The exploration of gene tree discordance has become common in the phylogenetic era (Salichos et al. 2014; Smith et al. 2015; Huang et al. 2016; Pease et al. 2018) and is essential for understanding the underlying processes that shape the Tree of Life. Discordance among gene trees can be the product of multiple sources. These include errors and noise in data assembly and filtering, hidden paralogy, incomplete lineage sorting (ILS), gene duplication/loss (Pamilo and Nei 1988; Doyle 1992; Maddison 1997; Galtier and Daubin 2008), random noise from uninformative genes, as well as misspecified model parameters of molecular evolution such as substitutional saturation, codon usage bias, or compositional heterogeneity (Foster 2004; Cooper 2014; Cox et al. 2014; Liu et al. 2014). Among these potential sources of gene tree discordance, ILS is the most studied in the systematics literature (Edwards 2009), and several phylogenetic inference methods have been developed to accommodate ILS as the source of discordance (reviewed in Edwards et al. 2016; Mirarab et al. 2016; Xu and Yang 2016). More recently, methods that account for additional processes such as hybridization or introgression have gained attention. These include methods that estimate phylogenetic networks while accounting for ILS and hybridization simultaneously (e.g., Solís-Lemus and Ané 2016; Wen et al. 2018), and methods that detect introgression based on site patterns or phylogenetic invariants (e.g., Green et al. 2010; Durand et al. 2011; Kubatko and Chifman 2019). Frequently, multiple processes can contribute to gene tree heterogeneity (Holder et al. 2001; Buckley et al. 2006; Meyer et al. 2017; Knowles et al. 2018). However, at present, no method can estimate species trees from phylogenomic data while modeling multiple sources of conflict and heterogeneity in molecular substitution simultaneously. To overcome these limitations, the use of multiple phylogenetic tools and data partitioning schemes in phylogenomic datasets is essential to disentangle sources of gene tree heterogeneity and resolve recalcitrant relationships at deep and shallow nodes of the Tree of Life (e.g., Alda et al. 2019; Widhelm et al. 2019; Prasanna et al. 2020; Roycroft et al. 2020).

In this study, we evaluate multiple sources of gene tree conflict to test controversial hypotheses of ancient hybridization among subfamilies in the plant family Amaranthaceae s.l. Amaranthaceae s.l. includes the previously segregated family Chenopodiaceae (Hernández-Ledesma et al. 2015; The Angiosperm Phylogeny Group 2016). With ca. 2050 to 2500 species in 181 genera and a worldwide distribution (Hernández-Ledesma et al. 2015), Amaranthaceae s.l. is iconic for the repeated evolution of complex traits representing adaptations to extreme environments such as C_4_ photosynthesis in hot and often dry environments (e.g., Kadereit et al. 2012; Bena et al. 2017), various modes of extreme salt tolerance (e.g., Flowers and Colmer 2015; Piirainen et al. 2017) that in several species are coupled with heavy metal tolerance (Moray et al. 2016), and very fast seed germination and production of multiple diaspore types on one individual (Kadereit et al. 2017). Several important crops are members of Amaranthaceae s.l., such as the pseudocereals quinoa and amaranth, sugar beet, spinach, glassworts, and saltworts. Many species of the family are also important fodder plants in arid regions and several are currently being investigated for their soil remediating and desalinating effects (e.g., Li et al. 2019). Due to their economic importance, reference genomes are available for *Beta vulgaris* (sugar beet, subfamily Betoideae; Dohm et al. 2014), *Chenopodium quinoa* (quinoa, Chenopodioideae; Jarvis et al. 2017), *Spinacia oleracea* (spinach; Chenopodioideae; Xu et al. 2017) and *Amaranthus hypochondriacus* (amaranth; Amaranthoideae; Lightfoot et al. 2017), representing three of the 13 currently recognized subfamilies of Amaranthaceae s.l. (sensu Kadereit et al. 2003; Kadereit et al. 2017).

Within the core Caryophyllales the previously segregated families Amaranthaceae s.s. and Chenopodiaceae have always been regarded as closely related, and their separate family status has long been the subject of phylogenetic and taxonomic debate (Kadereit et al. 2003; Masson and Kadereit 2013; Hernández-Ledesma et al. 2015; Walker et al. 2018; Fig. 1). Their close affinity is supported by a number of shared morphological, anatomical and phytochemical synapomorphies, and has been substantiated by molecular phylogenetic studies (discussed in Kadereit et al. 2003). Amaranthaceae s.s. has a predominantly tropical and subtropical distribution with the highest diversity found in the Neotropics, eastern and southern Africa and Australia (Müller and Borsch 2005), while Chenopodiaceae predominantly occurs in temperate regions and semi-arid or arid environments of subtropical regions (Kadereit et al. 2003). The key problem has always been the species-poor and heterogeneous subfamilies Polycnemoideae and Betoideae, neither of which fit easily within Chenopodiaceae or Amaranthaceae s.s. (cf. Table 5 in Kadereit et al. 2003). Polycnemoideae is similar in ecology and distribution to Chenopodiaceae but shares important floral traits such as petaloid tepals, filament tubes and 2-locular anthers with Amaranthaceae s.s. Morphologically, Betoideae fits into either Chenopodiaceae or Amaranthaceae s.s. but has a unique fruit type—a capsule that opens with a circumscissile lid (Kadereit et al. 2006). Both Betoideae and Polycnemoideae possess only a few species each and each has a strongly disjunct distribution pattern across three continents. Furthermore, the genera of both subfamilies display a number of morphologically dissociating features. Both intercontinental disjunctions of species-poor genera and unique or intermediate morphological traits led to the hypothesis that Betoideae and Polycnemoideae might have originated from hybridization events among early-branching lineages in Amaranthaceae s.l. (Hohmann et al. 2006; Masson and Kadereit 2013). To test this hypothesis, a phylotranscriptomic approach is particularly compelling as it not only provides thousands of low-copy nuclear genes for dissecting sources of phylogenetic discordance, but also enables future studies associating gene tree topology with gene function and habitat adaptation.

**FIGURE 1.**
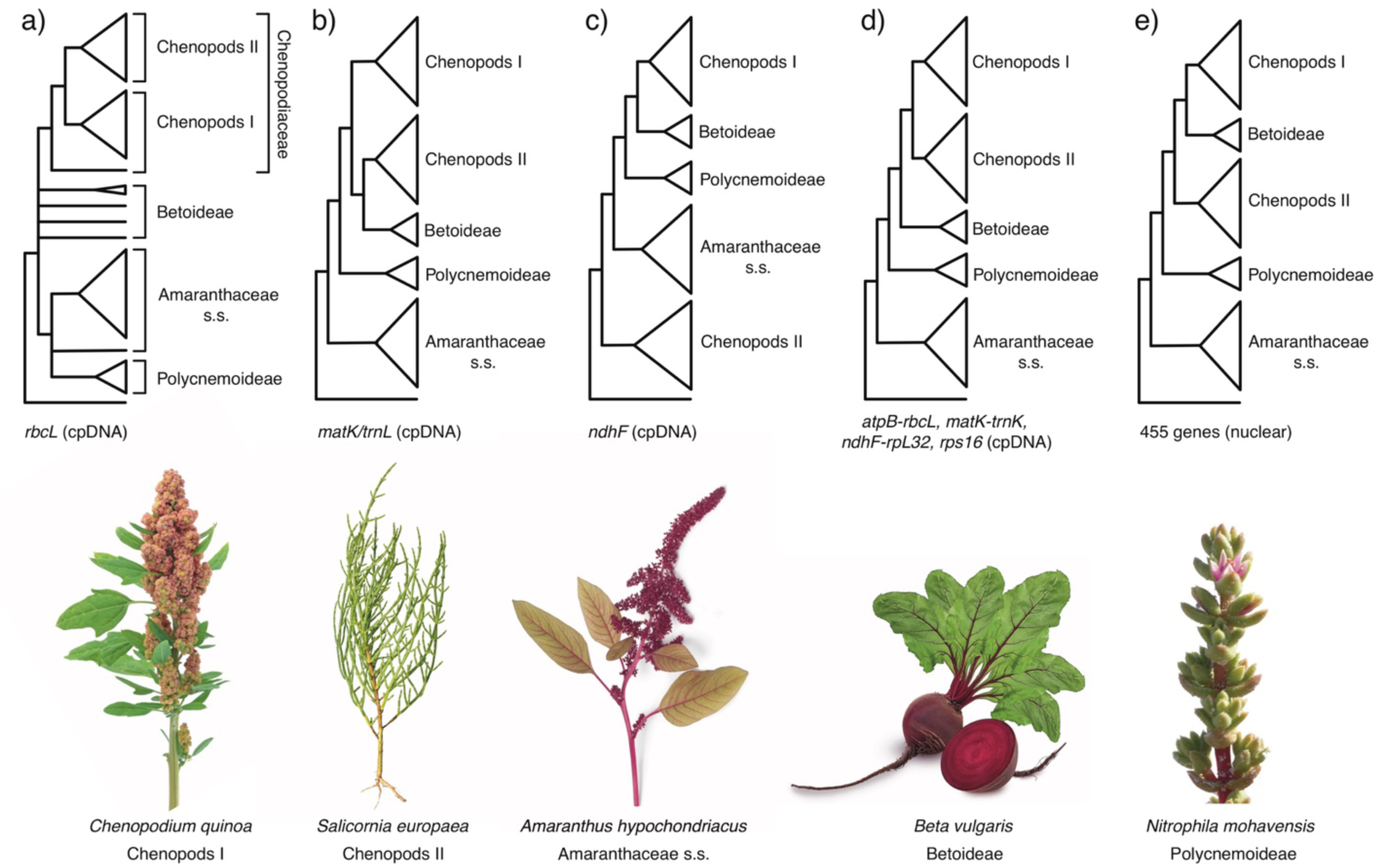
Phylogenetic hypothesis of Amaranthaceae s.l. from previous studies. a) Kadereit et al. (2003) using the plastid (cpDNA) *rbcL* coding region. b) Müller and Borsch (2005); using the cpDNA *matK* coding region and partial *trnL* intron. c) Hohmann et al. (2006) using the cpDNA *ndhF* coding region. d) Kadereit et al. (2017) using the cpDNA *atpB-rbcL* spacer, *matK* with *trnL* intron, *ndhF-rpL32* spacer, and *rps16* intron e) Walker et al. (2018) using 455 nuclear genes from transcriptome data. Major clades of Amaranthaceae s.l. named following the results of this study. Image credits: *Amaranthus hypochondriacus* by Picture Partners, *Beta vulgaris* by Olha Huchek, *Chenopodium quinoa* by Diana Mower, *Nitrophila mohavensis* by James M. André, and *Salsola soda* by Homeydesign.

Previous molecular phylogenetic analyses struggled to resolve the relationships among Betoideae, Polycnemoideae and the rest of the Amaranthaceae s.l. (Fig. 1). The first phylogenomic study of Amaranthaceae s.l. using nuclear loci (Walker et al. 2018; Fig. 1e) revealed that gene tree discordance mainly occurred at deep nodes of the phylogeny involving Betoideae. Polycnemoideae was sister to Chenopodiaceae, albeit supported by only 17% of gene trees, which contradicted previous analyses based on plastid data (Fig. 1, a–d). However, only a single species of Betoideae (the cultivated beet and its wild relative) was sampled in Walker et al. (2018). Furthermore, Walker et al. (2018) found conflicting topologies between concatenated and coalescent-based analyses, but sources of conflicting signals among gene trees remained unexplored.

In this study, we used a large genomic dataset to examine sources of gene tree discordance in Amaranthaceae s.l. Specifically, we tested whether Polycnemoideae and Betoideae result from independent hybridizations between Amaranthaceae s.s. and Chenopodioideae by distinguishing the signal of hybridization from gene tree discordance produced by ILS, uninformative gene trees, hidden paralogy, misspecifications of model of molecular evolution, and hard polytomy.

## Materials and Methods

An overview of all dataset and phylogenetic analyses can be found in Figure S1.

### Taxon sampling, transcriptome sequencing

We sampled 92 ingroup species (88 transcriptomes and four genomes) representing 53 genera (out of ca. 181) of all 13 currently recognized subfamilies and 16 out of 17 tribes of Amaranthaceae s.l. (sensu [Kadereit et al. 2003; Kadereit et al. 2017]). In addition, 13 outgroups across the Caryophyllales were included (ten transcriptomes and three genomes; Table S1). We generated 17 new transcriptomes for this study on an Illumina HiSeq2500 platform (Table S2). Library preparation was carried out using either poly-A enrichment or ribosomal RNA depletion. See Supplemental Methods for details on tissue collection, RNA isolation, library preparation, and quality control.

### Transcriptome data processing, assembly, homology and orthology inference

Read processing, assembly, translation, and homology and orthology inference followed the ‘phylogenomic dataset construction’ pipeline (Yang and Smith 2014) with multiple updates. We briefly describe our procedure below, with details in the Supplemental Methods and updated scripts in https://bitbucket.org/yanglab/phylogenomic_dataset_construction/

We processed raw reads for all 88 transcriptome datasets (except *Bienertia sinuspersici*) used in this study (Table S1). Reads were corrected for errors, trimmed for sequencing adapters and low-quality bases, and filtered for organellar reads. *De novo* assembly of processed nuclear reads was carried out with Trinity v 2.5.1 (Grabherr et al. 2011) with default settings, but without *in silico* normalization. Low-quality and chimeric transcripts were removed. Filtered transcripts were clustered into putative genes with Corset v 1.07 (Davidson and Oshlack 2014) and only the longest transcript of each putative gene was retained (Chen et al. 2019). Lastly, transcripts were translated, and identical coding sequences (CDS) were removed. Homology inference was carried out on CDS using reciprocal BLASTN, followed by orthology inference using the ‘monophyletic outgroup’ approach (Yang and Smith 2014), keeping only ortholog groups with at least 25 ingroup taxa.

### Assessment of recombination

Coalescent species tree methods assume that there is free recombination between loci and no recombination within loci. To determine the presence of recombination in our dataset, we used the pairwise homoplasy index test Φ for recombination, as implemented in PhiPack (Bruen et al. 2006). We tested recombination on the final set of ortholog alignments (with a minimum of 25 taxa) with the default sliding window size of 100 bp. Alignments that showed a strong signal of recombination with p ≤ 0.05 were removed from all subsequent phylogenetic analyses.

### Nuclear phylogenetic analysis

We used both concatenation and coalescent-based methods to reconstruct the phylogeny of Amaranthaceae s.l. Sequences from final orthologs were aligned using MAFFT v 7.307 (Katoh and Standley 2013) with settings ‘—genafpair --maxiterate 1000’. Columns with more than 70% missing data were trimmed with Phyx (Brown et al. 2017), and alignments with at least 1,000 characters and 99 out of 105 taxa were retained. We first estimated a maximum likelihood (ML) tree of the concatenated matrix with RAxML v 8.2.11 (Stamatakis 2014) using a partition-by-gene scheme with GTRCAT model for each partition and clade support assessed with 200 rapid bootstrap (BS) replicates. To estimate a coalescent-based species tree, first we inferred individual ML gene trees using RAxML with a GTRCAT model and 200 BS replicates to assess clade support. Gene trees were then used to infer a species tree with ASTRAL-III v5.6.3 (Zhang et al. 2018) using local posterior probabilities (LPP; Sayyari and Mirarab 2016) to assess clade support.

### Detecting and visualizing nuclear gene tree discordance

To explore discordance among gene trees, we first calculated the internode certainty all (ICA) value to quantify the degree of conflict on each node of a target tree (i.e., species tree) given individual gene trees (Salichos et al. 2014). In addition, we calculated the number of conflicting and concordant bipartitions on each node of the species trees. Both the ICA scores and conflicting/concordant bipartitions were calculated with Phyparts (Smith et al. 2015), mapping against the inferred ASTRAL species trees, using individual gene trees with BS support of at least 50% for the corresponding node. Additionally, in order to distinguish strong conflict from weakly supported branches, we carried out Quartet Sampling (QS; Pease et al. 2018) with 100 replicates. Quartet Sampling subsamples quartets from the input tree and alignment and assesses the confidence, consistency, and informativeness of each internal branch by the relative frequency of the three possible quartet topologies (Pease et al. 2018). Both ICA and Quartet Sampling scores provide an alternative branch support that reflects underlying gene tree conflict and that is not affected by anomalous high levels of bootstrap support common in phylogenomic data (Kumar et al. 2012).

To further visualize conflict, we built a cloudogram using DensiTree v2.2.6 (Bouckaert and Heled 2014). As DensiTree cannot accommodate missing taxa among gene trees, we reduced the final ortholog alignments to include 41 species (38 ingroup and 3 outgroups) in order to include as many orthologs as possible while representing all main clades of Amaranthaceae s.l. (see results). Individual gene trees were inferred as previously described. Trees were time-calibrated with TreePL v1.0 (Smith and O’Meara 2012) by fixing the crown age of Amaranthaceae s.l. to 66–72.1 based on a pollen record of *Polyporina cribraria* from the late Cretaceous (Maastrichtian; Srivastava 1969), and the root for the reduced 41-species dataset (most common recent ancestor of Achatocarpaceae and Aizoaceae) was set to 95 Ma based on the time-calibrated plastid phylogeny of Caryophyllales from Yao et al. (2019).

### Plastid assembly and phylogenetic analysis

Although DNase treatment was carried out to remove genomic DNA, due to their high copy number, plastid sequences are often carried over in RNA-seq libraries. In addition, as young leaf tissue was used for RNA-seq, the presence of RNA from plastid genes is expected to be represented. To investigate phylogenetic signal from plastid sequences, *de novo* assemblies were carried out with the Fast-Plast v.1.2.6 pipeline (https://github.com/mrmckain/Fast-Plast) using the filtered organelle reads. Contigs produced by Spades v 3.9.0 (Bankevich et al. 2012) were mapped to the closest available reference plastomes (Table S3), one copy of the Inverted Repeat was removed, and the remaining contigs manually edited in Geneious v.11.1.5 (Kearse et al. 2012) to produce the final oriented contigs.

Contigs were aligned with MAFFT with the setting ‘--auto’. Two samples (*Dysphania schraderiana* and *Spinacia turkestanica*) were removed due to low sequence occupancy. Using the annotations of the reference genomes (Table S3), the coding regions of 78 genes were extracted and each gene alignment was visually inspected in Geneious to check for potential misassemblies. From each gene alignment, taxa with short sequences (i.e., < 50% of the aligned length) were removed and the remaining sequences realigned with MAFFT. The genes *rpl32* and *ycf2* were excluded from downstream analyses due to low taxon occupancy (Table S4). For each individual gene we performed extended model selection (Kalyaanamoorthy et al. 2017) followed by ML gene tree inference and 1,000 ultrafast bootstrap replicates for branch support (Hoang and Chernomor 2018) in IQ-TREE v.1.6.1 (Nguyen et al. 2015). For the concatenated matrix we searched for the best partition scheme (Lanfear et al. 2012) followed by ML gene tree inference and 1,000 ultrafast bootstrap replicates for branch support in IQ-Tree. Additionally, we evaluated branch support with QS using 1,000 replicates and gene tree discordance with PhyParts. Lastly, to identify the origin of the plastid reads (i.e., genomic or RNA), we predicted RNA editing from CDS alignments using PREP (Mower 2009) with the alignment mode (PREP-aln), and a cutoff value of 0.8.

### Species network analysis using a reduced 11-taxon dataset

We inferred species networks that model ILS and gene flow using a maximum pseudo-likelihood approach (Yu and Nakhleh 2015). Species network searches were carried out with PhyloNet v.3.6.9 (Than et al. 2008) with the command ‘InferNetwork_MPL’ and using the individual gene trees as input. Due to computational restrictions, and given our main focus to identify potential reticulating events among major clades of Amaranthaceae s.l., we reduced our taxon sampling to one outgroup and ten ingroup taxa to include two representative species from each of the five well-supported major lineages in Amaranthaceae s.l. (see results). We filtered the final 105-taxon ortholog set to include genes that have all 11 taxa [referred herein as 11-taxon(net) dataset; Fig S1.]. After alignment and trimming we kept genes with a minimum of 1,000 aligned base pairs and individual ML gene trees were inferred using RAxML with a GTRGAMMA model and 200 bootstrap replicates. We carried out five network searches by allowing one to five reticulation events and ten runs for each search. To estimate the optimum number of reticulations, we optimized the branch lengths and inheritance probabilities and computed the likelihood of the best scored network from each of the five maximum reticulation events searches. Network likelihoods were estimated given the individual gene trees using the command ‘CalGTProb’ in PhyloNet (Yu et al. 2012). Then, we performed model selection using the bias-corrected Akaike information criterion (AICc; Sugiura 1978), and the Bayesian information criterion (BIC; Schwarz 1978). The number of parameters was set to the number of branch lengths being estimated plus the number of hybridization probabilities being estimated. The number of gene trees used to estimate the likelihood was used to correct for finite sample size. To compare network models to bifurcating trees, we also estimated bifurcating concatenated ML and coalescent-based species trees and a plastid tree as previously described with the reduced 11-species taxon sampling.

### Hypothesis testing and detecting introgression using four-taxon datasets

Given the signal of multiple clades potentially involved in hybridization events detected by PhyloNet (see results), we next conducted quartet analyses to explore a single event at a time. First, we further reduced the 11-taxon(net) dataset to six taxa that included one outgroup genome (*Mesembryanthemum crystallinum*) and one ingroup from each of the five major ingroup clades: *Amaranthus hypochondriacus* (genome), *Beta vulgaris* (genome), *Chenopodium quinoa* (genome), *Caroxylon vermiculatum* (transcriptome), and *Polycnemum majus* (transcriptome) to represent Amaranthaceae s.s., Betoideae, ‘Chenopods I’, ‘Chenopods II’ and Polycnemoideae, respectively. We carried out a total of ten quartet analyses using all ten four-taxon combinations that included three out of five ingroup species and one outgroup. We filtered the final set of 105-taxon orthologs for genes with all four taxa for each combination and inferred individual gene trees as described before. For each quartet we carried out the following analyses. We first estimated a species tree with ASTRAL and explored gene tree conflict with PhyParts. We then explored individual gene tree resolution by calculating the Tree Certainty (TC) score (Salichos et al. 2014) in RAxML using the majority rule consensus tree across the 200 bootstrap replicates. Next, we explored potential correlation between TC score and alignment length, GC content and alignment gap proportion using a linear regression model in R v.3.6.1 (R Core Team 2019). Lastly, we tested for the fit of gene trees to the three possible rooted quartet topologies for each gene using the approximately unbiased (AU) tests (Shimodaira 2002). We carried out ten constraint searches for each of three topologies in RAxML with the GTRGAMMA model, then calculated site-wise log-likelihood scores for the three constraint topologies in RAxML using GTRGAMMA and carried out the AU test using Consel v.1.20 (Shimodaira and Hasegawa 2001). In order to detect possible introgression among species of each quartet, first we estimated a species network with PhyloNet using a full maximum likelihood approach (Yu et al. 2014) with 100 runs per search while optimizing the likelihood of the branch lengths and inheritance probabilities for every proposed species network. Furthermore, we also carried out the ABBA/BABA test to detect introgression (Green et al. 2010; Durand et al. 2011) in each of four-taxon species trees. We calculated the *D-*statistic and associated *z* score for the null hypothesis of no introgression (*D* = 0) following each quartet ASTRAL species tree for taxon order assignment using 100 jackknife replicates and a block size of 10,000 bp with evobiR v1.2 (Blackmon and Adams) in R.

Additionally, to detect any non-random genomic block of particular quartet topology (Fontaine et al. 2015), we mapped the physical location of genes supporting each alternative quartet topology onto the *Beta vulgaris* reference genome using a synteny approach (See Supplemental Information for details).

### Assessment of substitutional saturation, codon usage bias, compositional heterogeneity, and model of sequence evolution misspecification

Analyses were carried out in a 11-taxon dataset [referred herein as 11-taxon(tree); Fig. S1] that included the same taxa used for species network analyses, but was processed differently to account for codon structure (see Supplemental Methods for details). Saturation was evaluated by plotting the uncorrected genetic distances of the concatenated alignment against the inferred distances (see Supplemental Methods for details). To determine the effect of saturation in the phylogenetic inferences we estimated individual ML gene trees using an unpartitioned alignment, a partition by first and second codon positions, and the third codon positions, and by removing all third codon positions. All tree searches were carried out in RAxML with a GTRGAMMA model and 200 bootstrap replicates. We then estimated a coalescent-based species tree and explored gene tree discordance with PhyParts.

Codon usage bias was evaluated using a correspondence analysis of the Relative Synonymous Codon Usage (RSCU; see Supplemental Methods for details). To determine the effect of codon usage bias in the phylogenetic inferences we estimated individual gene trees using codon-degenerated alignments (see Supplemental Methods for details). Gene tree inference and discordance analyses were carried out on the same three data schemes as previously described.

Among-lineage compositional heterogeneity was evaluated on individual genes using a compositional homogeneity test (Supplemental Methods for details). To assess if compositional heterogeneity had an effect in species tree inference and gene tree discordance, gene trees that showed the signal of compositional heterogeneity were removed from saturation and codon usage analyses and the species tree and discordance analyses were rerun.

To explore the effect of sequence evolution model misspecification, we reanalyzed the datasets from the saturation and codon usage analyses using inferred gene trees that accounted for model selection. Additionally, we also explored saturation and model misspecification in phylogenetic trees from amino acid alignments (see Supplemental Methods for details).

### Polytomy test

To test if the gene tree discordance among the main clades of Amaranthaceae s.l. could be explained by polytomies instead of bifurcating nodes, we carried out the quartet-based polytomy test by Sayyari and Mirarab (2018) as implemented in ASTRAL. We performed the polytomy test using the gene trees inferred from the saturation and codon usage analyses [11-taxon(tree) dataset]. Because this test can be sensitive to gene tree error (Syyari and Mirarab 2018), we performed a second test using gene trees where branches with less than 75% of bootstrap support were collapsed.

### Coalescent simulations

To investigate if gene tree discordance can be explained by ILS alone, we carried out coalescent simulations similar to Cloutier et al. (2019). An ultrametric species tree with branch lengths in mutational units (µT) was estimated by constraining an ML tree search of the 11-taxon(net) concatenated alignment to the ASTRAL species tree topology with a GTR+GAMMA model while enforcing a strict molecular clock in PAUP v4.0a (build 165; Swofford 2002). The mutational branch lengths from the constrained tree and branch lengths in coalescent units (τ= T/4N_e_) from the ASTRAL species trees were used to estimate the population size parameter theta (?= µT/τ; Degnan and Rosenberg 2009) for internal branches. Terminal branches were set with a population size parameter theta of one. We used the R package Phybase v. 1.4 (Liu and Yu 2010) that uses the formula from Rannala and Yang (2003) to simulate 10,000 gene trees using the constraint tree and the estimated theta values. Then we calculated the distribution of Robinson and Foulds (1981) tree-to-tree distances between the species tree and each gene tree using the R package Phangorn v2.5.3 (Schliep 2011), and compared this with the distribution of tree-to-tree distances between the species tree and the simulated gene tree. We ran simulations using the species tree and associated gene tree distribution from the original no partition 11-taxon(net).

### Test of the anomaly zone

The anomaly zone occurs where a set of short internal branches in the species tree produces gene trees that differ from the species tree more frequently than those that are concordant [a(x); as defined in equation 4 of Degnan and Rosenberg (2006)]. To explore if gene tree discordance observed in Amaranthaceae s.l. is a product of the anomaly zone, we estimated the boundaries of the anomaly zone [a(x); as defined in equation 4 of Degnan and Rosenberg (2006)] for the internal nodes of the species tree. Here, x is the branch length in coalescent units in the species tree that has a descendant internal branch. If the length of the descendant internal branch (y) is smaller than a(x), then the internode pair is in the anomaly zone and is likely to produce anomalous gene trees (AGTs). We carried out the calculation of a(x) following Linkem et al. (2016) in the same 11-taxon(tree) ASTRAL species tree used for coalescent simulations. Additionally, to establish the frequency of gene trees that were concordant with the estimated species trees, we quantified the frequency of all 105 possible rooted gene trees with Amaranthaceae s.l. being monophyletic.

## Results

### Transcriptome sequencing, assembly, translation, and quality control

Raw reads for the 17 newly generated transcriptomes are available from the NCBI Sequence Read Archive (BioProject: PRJNA640363; Table S2). The number of raw read pairs ranged from 17 to 27 million. For the 16 samples processed using RiboZero, organelle reads accounted for 15% to 52% of read pairs (Table S2). For *Tidestromia oblongifolia* that poly-A enrichment was carried out in library prep with ∼5% of raw reads were from organelle (Table S2). The final number of orthologs was 13,024 with a mean of 9,813 orthologs per species (Table S1). Of those, 82 orthologs had a strong signal of recombination (*P* ≤ 0.05) and were removed from downstream analyses.

### Analysis of the nuclear dataset of Amaranthaceae s.l

The final set of nuclear orthologous genes included 936 genes with at least 99 out of 105 taxa and 1,000 bp in aligned length after removal of low occupancy columns (the 105-taxon dataset). The concatenated matrix consisted of 1,712,054 columns with a gene and character occupancy of 96% and 82%, respectively. The species tree from ASTRAL and the concatenated ML tree from RAxML recovered the exact same topology with most clades having maximal support [i.e., bootstrap percentage (BS) = 100, local posterior probabilities (LPP) = 1; Fig. 2; Figs S2–S3]. Both analyses recovered Chenopodiaceae as monophyletic with the relationships among major clades concordant with the cpDNA analysis from Kadereit et al. (2017; Fig. 1d). Betoideae was placed as sister of Chenopodiaceae, while Polycnemoideae was strongly supported as sister (BS = 97, LPP = 0.98) to the clade composed of Chenopodiaceae and Betoideae. Amaranthaceae s.s. had an overall topology concordant to Kadereit et al. (2017), with the exception of *Iresine*, which was recovered among the Aervoids (Fig. 2; Figs S2–S3).

**FIGURE 2.**
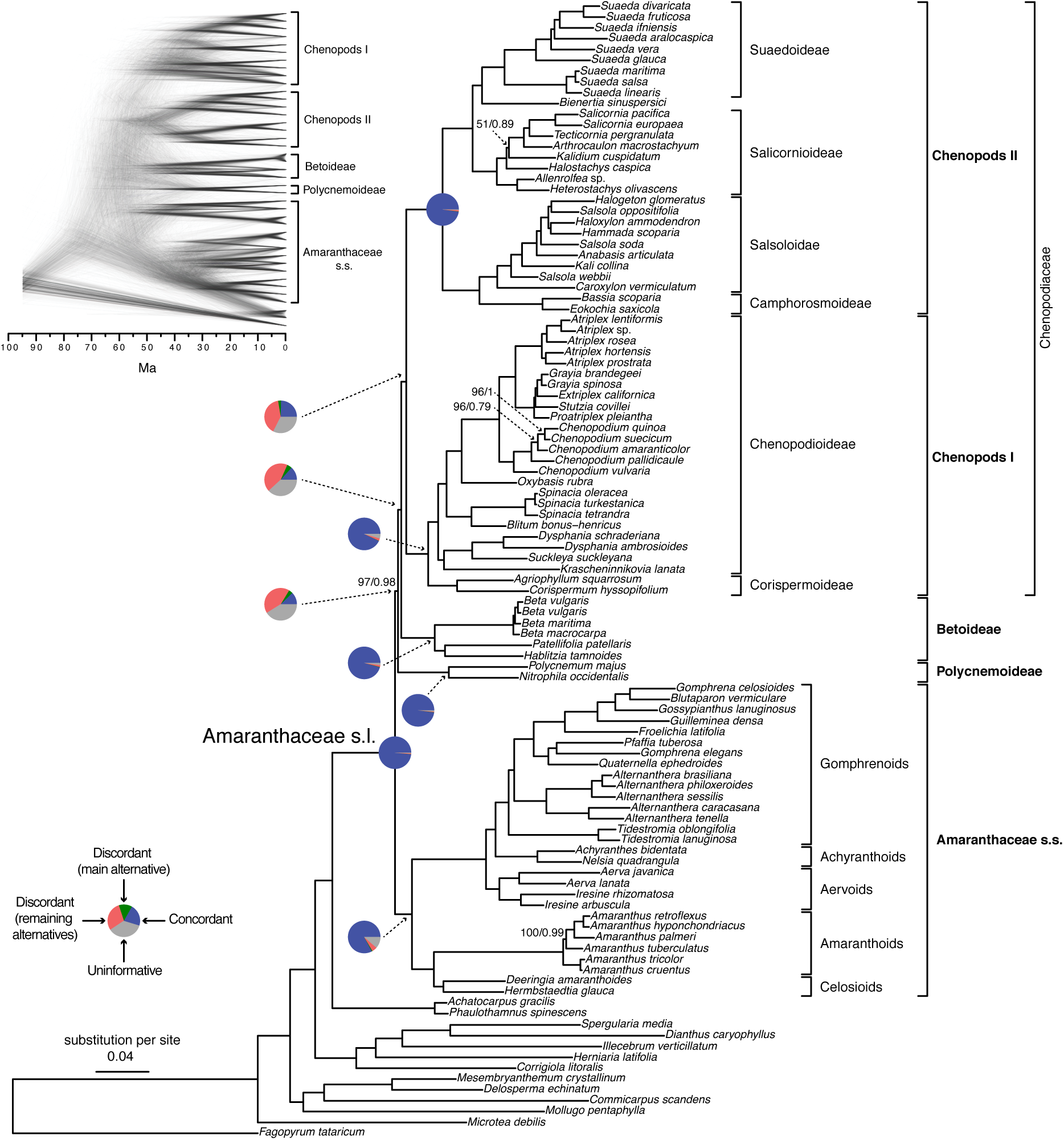
Maximum likelihood phylogeny of Amaranthaceae s.l. inferred from RAxML analysis of the concatenated 936-nuclear gene supermatrix, which had the same topology as recovered from ASTRAL. All nodes have maximal support (bootstrap = 100/ASTRAL local posterior probability = 1) unless noted. Pie charts present the proportion of gene trees that support that clade (blue), support the main alternative bifurcation (green), support the remaining alternatives (red), and the proportion (conflict or support) that have < 50% bootstrap support (gray). Only pie charts for major clades are shown (see Fig. S2 for all node pie charts). Branch lengths are in number of substitutions per site. The inset (top left) shows the Densitree cloudogram inferred from 1,242 nuclear genes for the reduced 41-taxon dataset.

The conflict analyses confirmed the monophyly of Amaranthaceae s.l. with 922 out of 930 informative gene trees being concordant (ICA= 0.94) and having full QS support (1/–/1; i.e., all sampled quartets supported that branch). Similarly, the monophyly of Amaranthaceae s.s. was highly supported by 755 of 809 informative gene trees (ICA =0.85) and the QS scores (0.92/0/1). However, the backbone of the family was characterized by high levels of gene tree discordance (Fig. 2; Figs S2–S3). The monophyly of Chenopodiaceae was supported only by 231 out of 632 informative gene trees (ICA = 0.42) and the QS score (0.25/0.19/0.99) suggested weak quartet support with a skewed frequency for an alternative placement of two well-defined clades within Chenopodiaceae, herein referred to as ‘Chenopods I’ and ‘Chenopods II’ (Fig. 2; Figs S2–S3). ‘Chenopods I’ and ‘Chenopods II’ were each supported by the majority of gene trees, 870 (ICA = 0.89) and 916 (ICA = 0.91), respectively and full QS support. Similarly, high levels of conflict among informative gene trees were detected in the placement of Betoideae (126 out of 579 informative genes being concordant, ICA = 0.28; QS score 0.31/0.57/1) and Polycnemoideae (116/511; ICA = 0.29;0.3/0.81/0.99). The Densitree cloudogram also showed significant conflict along the backbone of Amaranthaceae s.l. (Fig. 2).

Together, analysis of nuclear genes recovered five well-supported clades in Amaranthaceae s.l.: Amaranthaceae s.s., Betoideae, ‘Chenopods I’, ‘Chenopods II’, and Polycnemoideae. However, relationships among these five clades showed a high level of conflict among genes [ICA scores and gene counts (pie charts)] and among subsampled quartets (QS scores), despite having high support from both BS and LPP scores.

### Plastid phylogenetic analysis of Amaranthaceae s.l

RNA editing prediction analysis revealed editing sites only on CDS sequences of reference plastomes (Table S3), suggesting that cpDNA reads in RNA-seq libraries come from RNA rather than DNA leftover from incomplete DNase digestion during sample processing (See Discussion for details in plastid assembly from RNA-seq data).

The final alignment from 76 genes included 103 taxa and 55,517 bp in aligned length. The ML tree recovered the same five main clades within Amaranthaceae s.l. with maximal support (BS = 100; Figs S4–S6). Within each main clade, relationships were fully congruent with Kadereit et al. (2017) and mostly congruent with our nuclear analyses. However, the relationship among the five main clades differed from the nuclear tree. Here, the sister relationships between Betoideae and ‘Chenopods I’, and between Amaranthaceae s.s. and Polycnemoideae were both supported by BS =100. The sister relationship between these two larger clades was moderately supported (BS = 73), leaving ‘Chenopods II’ as sister to the rest of Amaranthaceae s.l.

Conflict analysis confirmed the monophyly of Amaranthaceae s.l. with 51 out of 69 informative gene trees supporting this clade (ICA = 0.29) and full QS support (1/–/1). On the other hand, and similar to the nuclear phylogeny, significant gene tree discordance was detected among plastid genes regarding placement of the five major clades (Figs S4–S6). The sister relationship of Betoideae and ‘Chenopods I’ was supported by only 20 gene trees (ICA = 0.06), but it had a strong support from QS (0.84/0.88/0/94). The relationship between Amaranthaceae s.s. and Polycnemoideae was supported by only 15 gene trees (ICA = 0.07), while QS showed weak support (0.41/0.21.0.78) with signals of a supported secondary evolutionary history. The clade uniting Betoideae, ‘Chenopods I’, Amaranthaceae s.s., and Polycnemoideae was supported by only four-gene trees, with counter-support from both QS (−0.29/0.42/0.75) and ICA (−0.03), suggesting that most gene trees and sampled quartets supported alternative topologies.

### Species network analysis of Amaranthaceae s.l

The reduced 11-taxon(net) dataset included 4,138 orthologous gene alignments with no missing taxon and a minimum of 1,000 bp (aligned length after removal of low occupancy columns). The 11-taxon(net) ASTRAL species tree was congruent with the 105-taxon tree, while both the nuclear and plastid ML trees from concatenated supermatrices had different topologies than their corresponding 105-taxon trees (Fig. 3). Model selection indicated that any species network was a better model than the best bifurcating nuclear or plastid trees (ASTRAL; AICc = 46972.9794; Table S5). PhyloNet identified up to five hybridization events among the clades of Amaranthaceae s.l. (Fig. 3), with the best model having five hybridization events involving all five clades (AICc = 28459.1835; Table S5). The best species network did not support the hypothesis of the hybrid origin of Betoideae or Polycnemoideae. Moreover, the best species network showed a complex reticulate pattern that involved mainly ‘Chenopods I’ and ‘Chenopods II’ (Fig. 3), but none of these reticulations events were supported by *D*-Statistic or species network results from the four-taxon analyses (see below).

**FIGURE 3.**
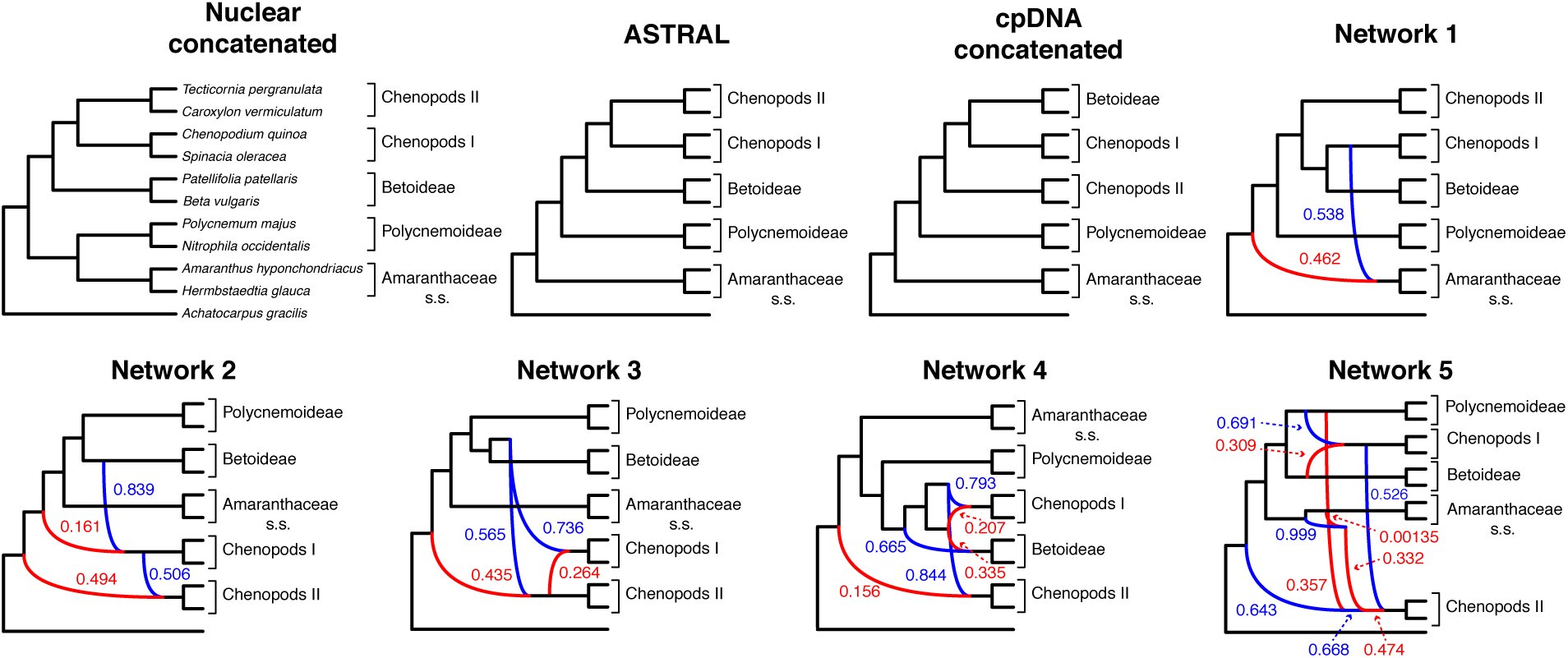
Species trees and species networks of the reduced 11-taxon(net) dataset of Amaranthaceae s.l. Nuclear concatenated phylogeny inferred from 4,138-nuclear gene supermatrix with RAxML. ASTRAL species tree inferred using 4,138 nuclear genes. cpDNA concatenated tree inferred from 76-plastid gene supermatrix with IQ-tree. Species network inferred from PhyloNet pseudolikelihood analyses with 1 to 5 maximum number of reticulations. Red and blue indicate the minor and major edges, respectively, of hybrid nodes. Number next to the branches indicates inheritance probabilities for each hybrid node.

### Four-taxon analyses

To test for hybridization events one at a time, we further reduced the 11-taxon(net) dataset to 10 four-taxon combinations that each included one outgroup and one representative each from three out of the five major ingroup clades. Between 7,756 and 8,793 genes were used for each quartet analysis (Table S6) and each quartet topology can be found in Figure 4. Only five out of the ten bifurcating quartet species trees (H0 and more frequent gene tree) were compatible with the nuclear species tree inferred from the complete 105-taxon dataset. The remaining quartets corresponded to the second most frequent gene tree topology in the 105-taxon nuclear tree, except for the quartet of Betoideae, ‘Chenopods II’ and Polycnemoideae (PBC2, which correspond to the least frequent gene tree).

**FIGURE 4.**
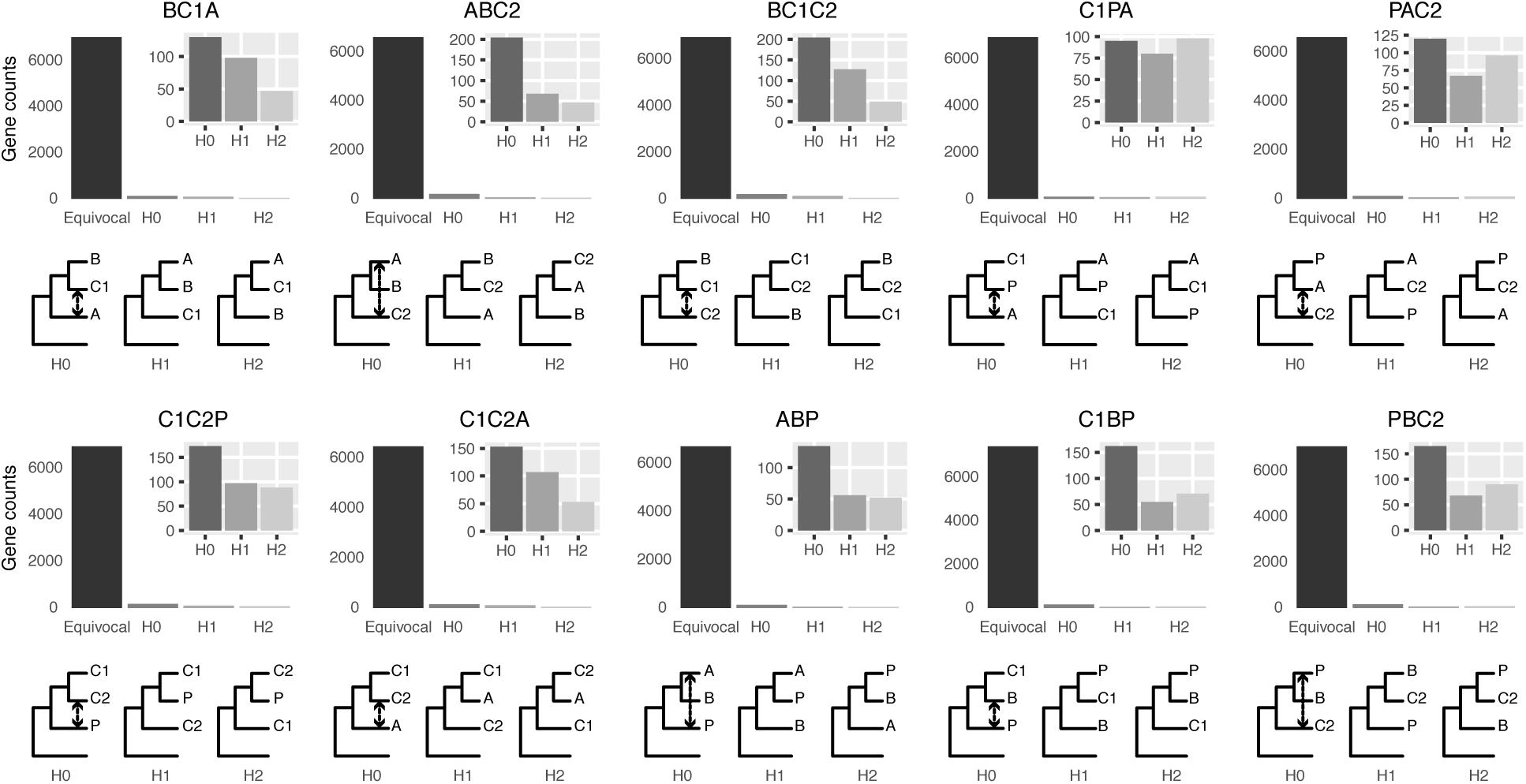
Gene counts from Approximate-Unbiased (AU) topology test of the 10 quartets from the five main clades of Amaranthaceae s.l. AU tests were carried out between the three possible topologies of each quartet. H0 represents the ASTRAL species tree of each quartet. “Equivocal” indicates gene trees that fail to reject all three alternative topologies for a quartet with p ≤ 0.05. Gene counts for each of the three alternative topologies represent gene trees supporting unequivocally one topology by rejecting the other two alternatives with p ≤ 0.05. Insets represent gene counts only for unequivocal topology support. Double arrowed lines in each H0 quartet represent the direction of introgression from the ABBA/BABA test. Each quartet is named following the species tree topology, where the first two species are sister to each other. A = Amaranthaceae s.s. (represented by *Amaranthus hypochondriacus*), B = Betoideae (*Beta vulgaris*), C1 = Chenopods I (*Chenopodium quinoa*), C2 = Chenopods II (*Caroxylum vermiculatum*), P = Polycnemoideae (*Polycnemum majus*). All quartets are rooted with *Mesembryanthemum crystallinum*.

In each of the ten quartets, the ASTRAL species tree topology (H0) was the most frequent among individual gene trees (raw counts) but only accounted for 35%–41% of gene trees, with the other two alternative topologies having balanced to slightly skewed frequencies (Fig. S7a; Table S7). Gene counts based on the raw likelihood scores from the constraint analyses showed similar patterns (Fig. S7b; Table S7). When filtered by significant likelihood support (i.e., ΔAICc ≥ 2), the number of trees supporting each of the three possible topologies dropped between 34% and 45%, but the species tree remained the most frequent topology for all quartets (Fig. S7b; Table S7). The AU topology tests failed to reject (*P* ≤ 0.05) approximately 85% of the gene trees for any of the three possible quartet topologies and rejected all but a single topology in only 3%–4.5% of cases. Among the unequivocally selected gene trees, the frequencies among the three alternative topologies were similar to ones based on raw likelihood scores (Fig S7; Table S7). Therefore, topology tests showed that most genes were uninformative for resolving the relationships among the major groups of Amaranthaceae s.l.

Across all ten quartets we found that most genes had very low TC scores (for any single node the maximum TC value is 1; Supplemental Fig. S8), showing that individual gene trees also had high levels of conflict among bootstrap replicates, which also indicated uninformative genes and was concordant with the AU topology test results. We were unable to detect any significant correlation between TC scores and alignment length, GC content or alignment gap fraction (Table S8), suggesting that filtering genes by any of these criteria was unlikely to increase the information content of the dataset.

Species network analyses followed by model selection using each of the four-taxon datasets showed that in seven out of the ten total quartets, the network with one hybridization event was a better model than any bifurcating tree topology. However, each of the best three networks from PhyloNet had very close likelihood scores and no significant ΔAICc among them (Table S6; Fig S9). For the remaining three quartets, the species trees (H0) was the best model.

The ABBA/BABA test results showed a significant signal of introgression within each of the ten quartets (Table S9; Fig 4). The possible introgression was detected between six out of the ten possible pairs of taxa. Potential introgression between Betoideae and Amaranthaceae s.s., ‘Chenopods I’ or ‘Chenopods II’, and between ‘Chenopods I’ and Polycnemoideae was not detected.

To further evaluate whether alternative quartets were randomly distributed across the genome, we mapped topologies from the quartet of Betoideae, ‘Chenopods II, and Amaranthaceae s.s. (BC1A) onto the reference genome of *Beta vulgaris*. We used the BC1A quartet as an example as all four species in this quartet have reference genomes. Synteny analysis between the diploid ingroup reference genome *Beta vulgaris* and the diploid outgroup reference genome *Mesembryanthemum crystallinum* recovered 22,179 collinear genes in 516 syntenic blocks. With the collinear ortholog pair information, we found that of the 8,258 orthologs of the BC1A quartet, 6,941 contained syntenic orthologous genes within 383 syntenic blocks. The distribution of the BC1A quartet topologies along the chromosomes of *Beta vulgaris* did not reveal any spatial clustering of any particular topology along the chromosomes (Fig. S10).

Gene Ontology enrichment analyses (not shown) using alternative topologies of the BC1A quartet did not recover any significant term associated with C_4_ photosynthesis, drought recovery, or salt stress response.

### Assessment of substitutional saturation, codon usage bias, compositional heterogeneity, sequence evolution model misspecification, and polytomy test

We assembled a second 11-taxon(tree) dataset that included 5,936 genes and a minimum of 300 bp (aligned length after removal of low occupancy columns) and no missing taxon. The saturation plots of uncorrected and predicted genetic distances showed that the first and second codon positions were unsaturated (y = 0.884x), whereas the slope of the third codon positions (y = 0.571x) showed a signal of saturation (Fig. S11). The correspondence analyses of RSCU show that some codons are more frequently used in different species, but overall the codon usage was randomly dispersed among all species and not clustered by clade (Fig. S12). This suggests that the phylogenetic signal is unlikely to be driven by differences in codon usage bias among clades. Furthermore, only 549 (∼9%) genes showed a signal of compositional heterogeneity (p < 0.05). The topology and support (LPP = 1.0) for all branches was the same for the ASTRAL species trees obtained from the different data schemes while accounting for saturation, codon usage, compositional heterogeneity, and model of sequence evolution, and was also congruent with the ASTRAL species tree and concatenated ML from the 105-taxon analyses (Fig. S13). In general, the proportion of gene trees supporting each bipartition remained the same in every analysis and showed high levels of conflict among the five major clades of Amaranthaceae s.l. (Fig S13).

The ASTRAL polytomy test resulted in the same bifurcating species tree for the 11-taxon(tree) dataset and rejected the null hypothesis that any branch is a polytomy (p < 0.01 in all cases). These results were identical when using gene trees with collapsed branches.

### Coalescent simulations and tests of the anomaly zone

The distribution of tree-to-tree distances of the empirical and simulated gene trees to the species tree from the 11-taxon(tree) dataset largely overlapped (Fig 5a), suggesting that ILS alone is able to explain most of the observed gene tree heterogeneity (Maureira-Butler et al. 2008). The anomaly zone limit calculations using species trees from the 11-taxon(tree) dataset detected two pairs of internodes among the five major groups in Amaranthaceae s.l. that fell into the anomaly zone (the red pair and the green pair, Fig. 5b; Table S10). Furthermore, gene tree counts showed that the species tree was not the most common gene tree topology, as defined for the anomaly zone (Degnan and Rosenberg 2006; Fig 5c). The species tree was the fourth most common gene tree topology (119 out of 4,425 gene trees), while the three most common gene tree topologies occurred 170, 136, and 127 times (Fig. 5c).

**FIGURE 5.**
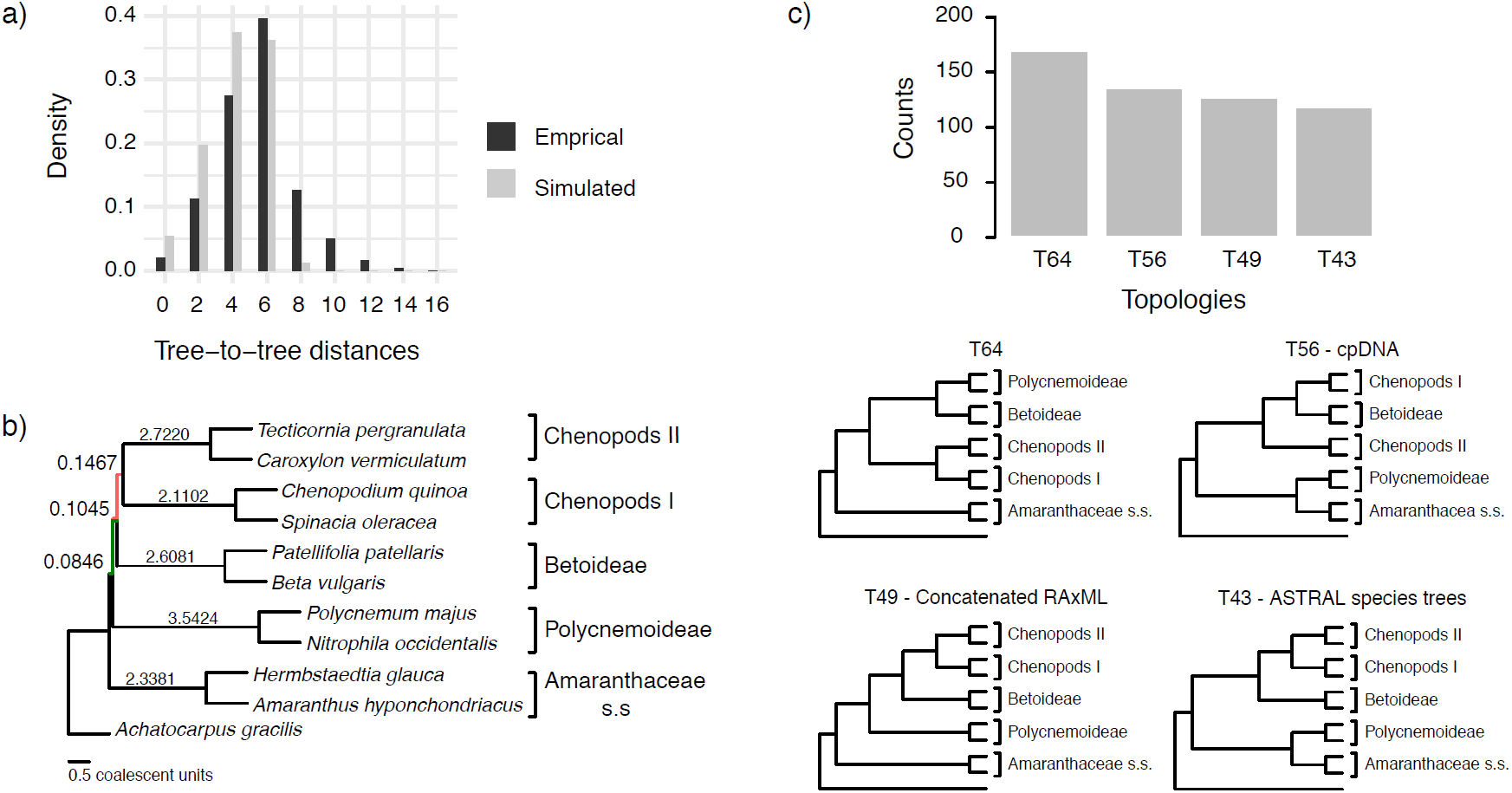
Coalescent simulations and tests of the anomaly zone from the 11-taxon(tree) dataset estimated from individual gene trees. a) Distribution of tree-to-tree distances between empirical gene trees and the ASTRAL species tree, compared to those from the coalescent simulation. b) ASTRAL species tree showing branch length in coalescent units. Green and red branches represent the internodes that fall in the anomaly zone (see Table S10 for anomaly zone limits). c) Gene tree counts (top) of the four most common topologies (bottom). Gene trees that do not support the monophyly of any of the five major clades were ignored.

## Discussion

The exploration of gene tree discordance has become a fundamental step to understand recalcitrant relationships across the Tree of Life. Recently, new tools have been developed to identify and visualize gene tree discordance (e.g., Salichos et al. 2014; Smith et al. 2015; Huang et al. 2016; Pease et al. 2018). However, downstream methods that evaluate processes generating observed patterns of gene tree discordance are still in their infancy. In this study, by combining transcriptomes and genomes, we were able to create a rich and dense dataset to start to tease apart alternative hypotheses concerning the sources of conflict along the backbone phylogeny of Amaranthaceae s.l. We found that gene tree heterogeneity observed in Amaranthaceae s.l. can be explained by a combination of processes, including ILS, ancient hybridization, and uninformative genes, that might have acted simultaneously and/or cumulatively.

### Gene tree discordance detected among plastid genes

Although both our concatenation-based plastid and nuclear phylogenies supported the same five major clades of Amaranthaceae s.l., the relationships among these clades are incongruent (Figs. 2 & S4). Cytonuclear discordance is well-known in plants and it has been traditionally attributed to reticulate evolution (Rieseberg and Soltis 1991; Sang et al. 1995; Soltis and Kuzoff 1995). Such discordance continues to be treated as evidence in support of hybridization in more recent phylogenomic studies that assume the plastome to be a single, linked locus (e.g., Folk et al. 2017; Vargas et al. 2017; Morales-Briones et al. 2018b; Lee-Yaw et al. 2019). However, recent work showed that the plastome might not necessarily act as a single locus and high levels of tree conflict have been detected (Gonçalves et al. 2019; Walker et al. 2019).

In Amaranthaceae s.l., previous studies based on plastid protein-coding genes or introns (Fig. 1; Kadereit et al. 2003; Müller and Borsch 2005; Hohmann et al. 2006; Kadereit et al. 2017) resulted in different relationships among the five main clades and none in agreement with our 76-gene plastid phylogeny. Our conflict and QS analyses of the plastid dataset (Figs S5–S6) revealed strong signals of gene tree discordance among the five major clades of Amaranthaceae s.l., likely due to heteroplasmy, although the exact sources of conflict are yet to be clarified (Gonçalves et al. 2019). Unlike the results found by Walker et al. (2019), our individual plastid gene trees had highly supported nodes (i.e., BS ≥ 70, Fig S5), suggesting that low phylogenetic information content is not the main source of conflict in our plastid dataset.

Our results support previous studies showing RNA-seq data can be a reliable source for plastome assembly (Smith 2013; Osuna-Mascaró et al. 2018; Gitzendanner et al. 2018). RNA-seq libraries can contain some genomic DNA due to incomplete digestion during RNA purification (Smith 2013). Given the AT-rich nature of plastomes, plastid DNA may survive the poly-A selection during mRNA enrichment (Schliesky et al. 2012). However, RNA editing prediction results showed that our Amaranthaceae s.l. cpDNA assemblies came from RNA rather than DNA contamination regardless of library preparation by poly-A enrichment (71 transcriptomes) or RiboZero (16 transcriptomes). Similarly, Osuna-Mascaró et al. (2018) also found highly similar plastome assemblies (i.e., general genome structure, and gene number and composition) from RNA-seq and genomic libraries, supporting the idea that plastomes are fully transcribed in photosynthetic eukaryotes (Shi et al. 2016). Furthermore, the backbone topology of our plastid tree built mainly from RNA-seq data (97 out of 105 samples) was consistent with a recent complete plastome phylogeny of Caryophyllales mainly from genomic DNA (Yao et al. 2019), showing the utility of recovering plastid gene sequences from RNA-seq data. Nonetheless, RNA editing might be problematic when combining samples from RNA-seq and genomic DNA, especially when resolving phylogenetic relationships among closely related species.

### Identifiability in methods for detecting reticulation events

All methods that we used to detect ancient hybridization inferred the presence of reticulation events. However, our results suggest that these methods all struggle with ancient, rapid radiations. Advances have been made in recent years in developing methods to infer species networks in the presence of ILS (reviewed in Elworth et al. 2019). These methods have been increasingly used in phylogenetic studies (e.g., Wen et al. 2016; Copetti et al. 2017; Morales-Briones et al. 2018a; Crowl et al. 2020). To date, however, species network inference is still computationally intensive and limited to a small number of species and a few hybridization events (Hejase and Liu 2016; but see Hejase et al. 2018 and Zhu et al. 2019). Furthermore, studies evaluating the performance of different phylogenetic network inference approaches are scarce and restricted to simple hybridization scenarios. Kamneva and Rosenberg (2017) showed that likelihood methods like Yu et al. (2014) are often robust to ILS and gene tree error when symmetric hybridization (equal genetic contribution of both parents) events are considered. While this approach usually does not overestimate hybridization events, it fails to detect skewed hybridization (unequal genetic contribution of both parents) events in the presence of significant ILS. Methods developed to scale to larger numbers of species and hybridizations like the ones using pseudo-likelihood approximations (i.e., Solís-Lemus and Ané 2016; Yu and Nakhleh 2015) are yet to be evaluated independently, but in the case of the Yu and Nakhleh (2015) method based on rooted triples, it cannot distinguish the correct network when other networks can produce the same set of triples (Yu and Nakhleh 2015). On the other hand, the method of Solís-Lemus and Ané (2016), based on unrooted quartets, is better at avoiding indistinguishable networks, but it is limited to only level-1 network scenarios.

Applying the above methods to our data set recovered multiple reticulation events. Analysis of our 11-taxon(net) dataset using a pseudo-likelihood approach detected up to five hybridization events involving all five major clades of Amaranthaceae s.l. (Fig. 3). Model selection, after calculating the full likelihood of the obtained networks, also chose the 5-reticulation species as the best model. Likewise, we found that any species network had a better ML score than a bifurcating tree (Table S5). However, further analyses demonstrated that full likelihood network searches with up to one hybridization event are indistinguishable from each other (Table S6), resembling a random gene tree distribution. This pattern can probably be explained by the high levels of gene tree discordance and lack of phylogenetic signal in the inferred quartet gene trees (Fig. 4), suggesting that the 11-taxon(net) network searches can potentially overestimate reticulation events due to high levels of gene tree error or ILS.

Using the *D*-Statistic (Green et al. 2010; Durand et al. 2011) we also detected signals of introgression in seven possible locations among the five main groups of Amaranthaceae s.l. (Table S9). The inferred introgression events agreed with at least one of the reticulation scenarios from the phylogenetic network analysis. However, the *D*-Statistic did not detect any introgression that involves Betoideae, which was detected in the phylogenetic network analysis with either four or five reticulations events. The *D*-Statistic has been shown to be robust to a wide range of divergence times, but it is sensitive to relative population size (Zheng and Janke 2018), which agrees with the notion that large effective population sizes and short branches increase the chances of ILS (Pamilo and Nei 1988) and in turn can dilute the signal for the *D*-Statistic (Zheng and Janke 2018). Recently, Elworth et al. (2018) found that multiple or ‘hidden’ reticulations can cause the signal of the *D*-statistic to be lost or distorted. Furthermore, when multiple reticulations are present, the traditional approach of dividing datasets into quartets can be problematic as it largely underestimates *D* values (Elworth et al. 2018). Given short internal branches in the backbone of Amaranthaceae s.l. and the phylogenetic network results showing multiple hybridizations, it is plausible that our *D*-statistic may be affected by these issues.

Our analysis highlights problems with identifiability in relying on *D-*statistic or phylogenetic network analysis alone to detect reticulation events, especially in cases of ancient and rapid diversification. Both analyses resulted in highly complex and inconsistent reticulate scenarios that cannot be distinguished from ILS or gene tree error. Hence, despite the use of genome-scale data and exhaustive hypothesis testing, support is lacking for the hybrid origin of Polycnemoideae or Betoideae, or any particular hybridization event among major groups in Amaranthaceae s.l. In addition to potential hybridization events, rapid speciation, short branches, and large ancestral population size all impacting our ability to resolve relationships among major clades in Amaranthaceae s.l. Simulating combinations of these scenarios is beyond the scope of this particular manuscript.

### ILS and the Anomaly Zone

ILS is ubiquitous in multi-locus phylogenetic datasets. In its most severe cases ILS produces the ‘anomaly zone’, defined as a set of short internal branches in the species tree that produce anomalous gene trees (AGTs) that are more likely than the gene tree that matches the species tree (Degnan and Rosenberg 2006). Rosenberg (2013) expanded the definition of the anomaly zone to require that a species tree contain two consecutive internal branches in an ancestor–descendant relationship in order to produce AGTs. To date, only a few empirical examples of the anomaly zone have been reported (Linkem et al. 2016; Cloutier et al. 2019). Our results show that the species tree of Amaranthaceae s.l. has three consecutive short internal branches that lay within the limits of the anomaly zone (i.e., y < a[x]; Fig. 5; Table S10) and that the species tree is not the most frequent gene tree (Fig. 4). While both lines of evidence support the presence of AGTs, it is important to point out that our quartet analysis showed that most quartet gene trees were equivocal (94–96%; Fig. 4), and therefore, were uninformative. Huang and Knowles (2009) pointed out that the gene tree discordance produced from the anomaly zone can be produced by uninformative gene trees and that for species trees with short branches the most probable gene tree topology is a polytomy rather than an AGT. Our ASTRAL polytomy test, however, rejected a polytomy along the backbone of Amaranthaceae s.l. in any of the gene tree sets used. While we did not test for polytomies in individual gene trees, our ASTRAL polytomy test using gene trees with branches of <75% bootstrap support collapsed also rejected the presence of a polytomy. Therefore, the distribution of gene tree frequency in combination with short internal branches in the species tree supports the presence of an anomaly zone in Amaranthaceae s.l.

### Taxonomic implications

Despite the strong signal of gene tree discordance, both nuclear and plastid datasets strongly supported five major clades within Amaranthaceae s.l.: Amaranthaceae s.s, ‘Chenopods I’, ‘Chenopods II’, Betoideae, and Polycnemoideae (Figs. 2 & S4). These five clades are congruent with morphology and previous taxonomic treatments of the group. However, the relationships among these five lineages remain elusive with our data. Taken together, our tests of sources of incongruence for these early-diverging nodes indicate that no single source such as a particular ancient hybridization event can confidently account for the strong signal of gene tree discordance, suggesting that the discordance results primarily from ancient and rapid lineage diversification. Thus, the backbone of Amaranthaceae s.l. remains, and likely will remain, unresolved even with genome-scale data. The stem age of Amaranthaceae s.l. dates back to the early Tertiary (Paleocene; Kadereit et al. 2012; Di Vincenzo et al. 2018; Yao et al. 2019), but due to nuclear and plastid gene tree along the backbone, the geographic origin of Amaranthaceae s.l. remains ambiguous.

Therefore, for the sake of taxonomic stability, we suggest retaining Amaranthaceae s.l. sensu APG IV (The Angiosperm Phylogeny Group 2016), which includes the previously recognized Chenopodiaceae. Amaranthaceae s.l. is characterized by a long list of anatomical, morphological and phytochemical characters such as minute sessile flowers with five tepals, a single whorl of epitepalous stamens, and one basal ovule (Kadereit et al. 2003). Here, we recognize five subfamilies within Amaranthaceae s.l. represented by the five well-supported major clades recovered in this study (Fig. 2): Amaranthoideae (Amaranthaceae s.s.), Betoideae, Chenopodioideae (‘Chenopods I’), Polycnemoideae, and Salicornioideae (‘Chenopods II’).

### Conclusions

Our analyses highlight the need to test for multiple sources of conflict in phylogenomic analyses, especially when trying to resolve phylogenetic relationships with extensive phylogenetic conflict. Furthermore, one needs to be aware of the strengths and limitations of different phylogenetic methods and be cautious about relying on any single analysis, for example in the usage of phylogenetics species networks over coalescent-based species trees (Blair and Ané 2020). We make the following recommendation on five essential steps towards exploring heterogeneous phylogenetic signals in phylogenomic datasets in general. 1) Study design: consider whether the taxon sampling and marker choice enable testing alternative sources of conflicting phylogenetic signal. For example, will there be sufficient phylogenetic signal and sufficient taxon coverage in individual gene trees for methods such as phylogenetic network analyses? 2) Data processing: care should be taken in data cleaning, partitioning (e.g., nuclear vs. plastid), and using orthology inference methods that explicitly address paralogy issues (e.g., tree-based orthology inference and synteny information). 3) Species tree inference: select species tree methods that accommodate the dataset size and data type (e.g., ASTRAL for gene tree-based inferences or SVDquartet [Chifman and Kubatko 2014] for SNP-based inferences), followed by visualization of phylogenetic conflict using tools such as the pie charts (e.g., PhyParts) and quartet-based tools (e.g., Quartet Sampling; Quadripartition Internode Certainty [Zhou et al. 2020]; Concordance Factors [Minh et al. 2020]). 4) Assessing hybridization: if phylogenetic conflict cannot be explained by processes like ILS, phylogenetic species network analyses (e.g., PhyloNet) reduced taxon sampling can be applied to test hybridization hypotheses given results in step 3; 5) Hypothesis testing: additional tests can be performed given the results of recommendation 3 and 4 and depending on the scenario. These could include testing for model misspecification, anomaly zone, uninformative gene tree, and if hybridization is hypothesized, testing putative reticulation events one at a time, as illustrated in this study.

Despite using genome-scale data and exhaustive hypothesis testing, the backbone phylogeny of Amaranthaceae s.l. remains unresolved, and we were unable to distinguish ancient hybridization events from ILS or uninformative gene trees. Similar situations might not be atypical across the Tree of Life. As we leverage more genomic data and explore gene tree discordance in more detail, these steps will be informative in other clades, especially in those that are products of ancient and rapid lineage diversification (e.g., Widhelm et al. 2019; Koenen et al. 2020). Ultimately, such endeavors will be instrumental in gaining a full understanding of the complexity of the Tree of Life.

## Supplementary Material

Data available from the Dryad Digital Repository: http://dx.doi.org/10.5061/.[NNNN]

## Acknowledgments

The authors thank H. Freitag, J.M. Bena and the Millennium Seed Bank for providing seeds; U. Martiné for assisting with RNA extraction; N. Wang and Y.-Y. Huang for sample sequencing. A. Crum, R. Ree, B. Carstens, and three anonymous reviewers for providing helpful comments; the Minnesota Supercomputing Institute (MSI) at the University of Minnesota for providing access to computational resources. This work was supported by the University of Minnesota, the University of Michigan, the US National Science Foundation (DEB 1354048), and the Department of Energy, Office of Science, Genomic Science Program (Contract Number DE-SC0008834).

