## Supplemental Material for "Disentangling Sources of Gene Tree Discordance in Phylogenomic Datasets: Testing Ancient Hybridizations in Amaranthaceae s.l"

### SUPPLEMENTARY METHODS

#### *Library preparation*

Library preparation was carried out using either poly-A enrichment of ribosomal RNA depletion. For *Tidestromia oblongifolia*, tissue collection, RNA isolation, library preparation was carried out using the KAPA Stranded mRNA-Seq Kits (KAPA Biosystems, Wilmington, Massachusetts, USA). The library was multiplexed with 10 other samples from a different project on an Illumina HiSeq2500 platform with V4 chemistry at the University of Michigan Sequencing Core. For the remaining 16 samples total RNA was isolated from c. 70-125 mg leaf tissue collected in liquid nitrogen using the RNeasy Plant Mini Kit (Qiagen) following the manufacturer's protocol (June 2012). A DNase digestion step was included with the RNase-Free DNase Set (Qiagen). Quality and quantity of RNA were checked using the 2100 Bioanalyzer (Agilent Technologies). Library preparation was carried out using the TruSeq® Stranded Total RNA Library Prep Plant with RiboZero probes (96 Samples. Illumina, #20020611; Schuierer et al. 2017). Indexed libraries were normalized, pooled and size selected to 320bp +/- 5% using the Pippin Prep HT instrument to generate libraries with mean inserts of 200 bp, and sequenced on the Illumina HiSeq2500 platform with V4 chemistry at the University of Minnesota Genomics Center. Reads from all 17 libraries were paired-end 125 bp.

*Transcriptome data processing and assembly*

Scripts and instructions for read processing, assembly, translation, and homology and orthology search can be found at https://bitbucket.org/yanglab/phylogenomic_dataset_construction/ as part of an updated 'phylogenomic dataset construction' pipeline (Yang and Smith 2014).

We processed raw reads for all 88 transcriptome datasets (except *Bienertia sinuspersici*) used in this study (Table S1). Sequencing errors in raw reads were corrected with Rcorrector (Song and Florea 2015) and reads flagged as uncorrectable were removed. Sequencing adapters and low-quality bases were removed with Trimmomatic v0.36 (ILLUMINACLIP: TruSeq_ADAPTER: 2:30:10 SLIDINGWINDOW: 4:5 LEADING: 5 TRAILING: 5 MINLEN: 25; Bolger et al. 2014). Additionally, chloroplast and mitochondrial reads were filtered with Bowtie2 v 2.3.2 (Langmead and Salzberg 2012) using publicly available Caryophyllales organelle genomes from the Organelle Genome Resources database (RefSeq; [Pruitt et al. 2007]; last accessed on October 17, 2018) as references. Read quality was assessed with FastQC v 0.11.7 (https://www.bioinformatics.babraham.ac.uk/projects/fastqc/). Overrepresented sequences detected with FastQC were discarded. *De novo* assembly was carried out with Trinity v 2.5.1 (Grabherr et al. 2011) with default settings, but without in silico normalization. Assembly quality was assessed with Transrate v 1.0.3 (Smith-Unna et al. 2016). Low quality and poorly supported transcripts were removed using individual cut-off values for three contig score components of Transrate: 1) proportion of nucleotides in a contig that agrees in identity with the aligned read, $s(Cnuc) \leq 0.25$; 2) proportion of nucleotides in a contig that have one or more mapped reads, $s(Ccov) \leq 0.25$; and 3) proportion of reads that map to the contig in correct orientation, $s(Cord) \leq 0.5$. Furthermore, chimeric transcripts (*trans*-self and *trans*-multi-gene) were removed following the approach described in Yang and Smith (2013) using *Beta vulgaris* as the reference proteome, and percentage similarity and length cutoffs of 30 and 100, respectively. In order to remove isoforms and assembly artifacts, filtered reads were remapped to filtered transcripts with Salmon v 0.9.1 (Patro et al. 2017) and putative genes were clustered with Corset v 1.07 (Davidson and Oshlack 2014) using default settings, except that we used a minimum of five reads as threshold to remove transcripts with low coverage (-m 5). Only the longest transcript of each putative gene inferred by Corset was retained (Chen et al. 2019). Filtered transcripts were translated with TransDecoder v 5.0.2 (Haas et al. 2013) with default settings and the proteome of *Beta vulgaris* and *Arabidopsis thaliana* to identify open reading frames. Finally, coding sequences (CDS) from translated amino acids were further reduced with CD-HIT v 4.7 (-c 0.99; [Fu et al. 2012]) to remove near-identical sequences.

#### *Homology and orthology inference of nuclear genes*

Initial homology inference was carried out following Yang and Smith (2014) with some modifications. First, an all-by-all BLASTN search was performed on CDS using an *E* value cutoff of 10 and max_target_seqs set to 100. Raw BLAST output was filtered with a hit fraction of 0.4. Then putative homologs groups were clustered using MCL v 14-137 (van Dongen 2000) with a minimum minus log-transformed *E* value cutoff of 5 and an inflation value of 1.4. Finally, only clusters with a minimum of 25 taxa were retained. Individual clusters were aligned using MAFFT v 7.307 (Katoh and Standley 2013) with settings '–genafpair –maxiterate 1000'. Aligned columns with more than 90% missing data were removed using Phyx (Brown et al. 2017). Homolog trees were built using RAxML v 8.2.11 (Stamatakis 2014) with a GTRCAT model and clade support assessed with 200 rapid bootstrap (BS) replicates. Spurious or outlier long tips were detected and removed with TreeShrink v 1.0.0. by maximally reducing the tree diameter (Mai and Mirarab 2018). Monophyletic and paraphyletic tips that belonged to the same taxon were removed keeping the tip with the highest number of characters in the trimmed alignment. After visual inspection of ca. 50 homolog trees, we determined that internal branches longer than 0.25 were likely representing deep paralogs. These branches were cut apart, keeping resulting subclades with a minimum of 25 taxa. Homolog tree inference, tip and outlier removal, and deep paralog cutting was carried out for a second time using the same settings to obtain final homologs. Orthology inference was carried out following the 'monophyletic outgroup' approach from Yang and Smith (2014), keeping only ortholog groups with at least 25 ingroup taxa. The 'monophyletic outgroup' approach filters for clusters that have outgroup taxa being monophyletic and single-copy, and therefore filters for single- and low-copy genes. It then roots the gene tree by the outgroups, traverses the rooted tree from root to tip, and removes the side with less taxa when gene duplication is detected.

*Synteny analyses*

To visualize any genomic patterns of the phylogenetic history of *Beta vulgaris* regarding its relationship with Amaranthaceae s.s. and Chenopodiaceae, we first identified syntenic regions between the genomes of *Beta vulgaris* and the outgroup *Mesembryanthemum crystallinum* using the SynNet pipeline (https://github.com/zhaotao1987/SynNet-Pipeline; Zhao and Schranz 2019). We used DIAMOND v.0.9.24.125 (Buchfink et al. 2015) to perform all-by-all inter- and intra-pairwise protein searches with default parameters, and MCScanX (Wang et al. 2012) for pairwise synteny block detection with default parameters, except match score (-k) that was set to five. Then, we plot the nine chromosomes of *Beta vulgaris* by assigning each of the 8,258 orthologs of the quartet composed of *Mesembryanthemum crystallinum* (outgroup), *Amaranthus* *hypochondriacus, Beta vulgaris,* and *Chenopodium quinoa* (BC1A) to synteny blocks and to one of the three possible quartet topologies based on best likelihood score.

*Assessment of substitutional saturation, codon usage bias, compositional heterogeneity, and model of sequence evolution misspecification*

We refiltered the final 105-taxon ortholog alignments to again include genes that have the same 11 taxa (referred herein as 11-taxon(tree) dataset used for the species network analyses. We realigned individual genes using MACSE v.2.03 (Ranwez et al. 2018) to account for codon structure and frameshifts. Codons with frameshifts were replaced with gaps, and ambiguous alignment sites were removed using GBLOCKS v0.9b (Castresana 2000) while accounting for codon alignment (-t=c -b1=6 -b2=6 -b3=2 -b4=2 -b5=h). After realignment and removal of ambiguous sites, we kept genes with a minimum of 300 aligned base pairs. To detect potential saturation, we plotted the uncorrected genetic distances against the inferred distances as described in Philippe and Forterre (1999). The level of saturation was determined by the slope of the linear regression between the two distances where a shallow slope (i.e < 1) indicates saturation. We estimated the level of saturation by concatenating all genes and dividing the first and second codon positions from the third codon positions. We calculated uncorrected, and inferred distances with the TN93 substitution model using APE v5.3 (Paradis and Schliep 2019) in R. To determine the effect of saturation in the phylogenetic inferences we estimated individual gene trees using three partition schemes. We inferred ML trees with an unpartitioned alignment, a partition by first and second codon positions, and the third codon positions, and by removing all third codon positions. All tree searches were carried out in RAxML with a GTRGAMMA model and 200 bootstrap replicates. A species tree for each of the three data schemes was estimated with ASTRAL-III v5.6.3 (Zhang et al. 2018) and gene tree discordance was examined with PhyParts (Smith et al. 2015).

Codon usage bias was evaluated using a correspondence analysis of the Relative Synonymous Codon Usage (RSCU), which is defined as the number of times a particular codon is observed relative to the number of times that the codon would be observed in the absence of any codon usage bias (Sharp and Li 1986). RSCU for each codon in the 11-taxon concatenated alignment was estimated with CodonW v.1.4.4 (Peden 1999). Correspondence analysis was carried out using FactoMineR v1.4.1(Lê et al. 2008) in R (R Core Team 2019). . To determine the effect of codon usage bias in the phylogenetic inferences we estimated individual gene trees using codon-degenerated alignments. Alignments were recoded to eliminate signals associated with synonymous substitutions by degenerating the first and third codon positions using ambiguity coding using DEGEN v1.4 (Regier et al. 2010; Zwick et al. 2012). Gene tree inference and discordance analyses were carried out on the same three data schemes as previously described.

To examine the presence of among-lineage compositional heterogeneity, individual genes were evaluated using the compositional homogeneity test that uses a null distribution from simulations as proposed by Foster (2004). We performed the compositional homogeneity test by optimizing individual gene trees with a GTRGAMMA model and 1,000 simulations in P4 (Foster 2004). To assess if compositional heterogeneity had an effect in species tree inference and gene tree discordance, gene trees that showed the signal of compositional heterogeneity were removed from saturation and codon usage analyses and the species tree and discordance analyses were rerun.

To explore the effect of sequence evolution model misspecification, we reanalyzed the datasets from the saturation and codon usage analyses using inferred gene trees that accounted for model selection. We performed extended model selection followed by ML gene tree inference and 1,000 ultrafast bootstrap replicates for branch support in IQ-Tree v.1.6.1 (Nguyen et al. 2015). Species tree inference, conflict analysis and removal of genes with compositional heterogeneity were carried out as previously described.

Finally, we also used amino acid alignments from MACSE to account for substitutional saturation. Amino acid positions with frameshifts were replaced with gaps, and ambiguous alignment sites were removed with Phyx requiring a minimal occupancy of 30%. We inferred individual gene trees with IQ-tree to account for a model of sequence evolution and carried out species tree inference, conflict analysis, and removal of genes with compositional heterogeneity as described for the nucleotide alignments.

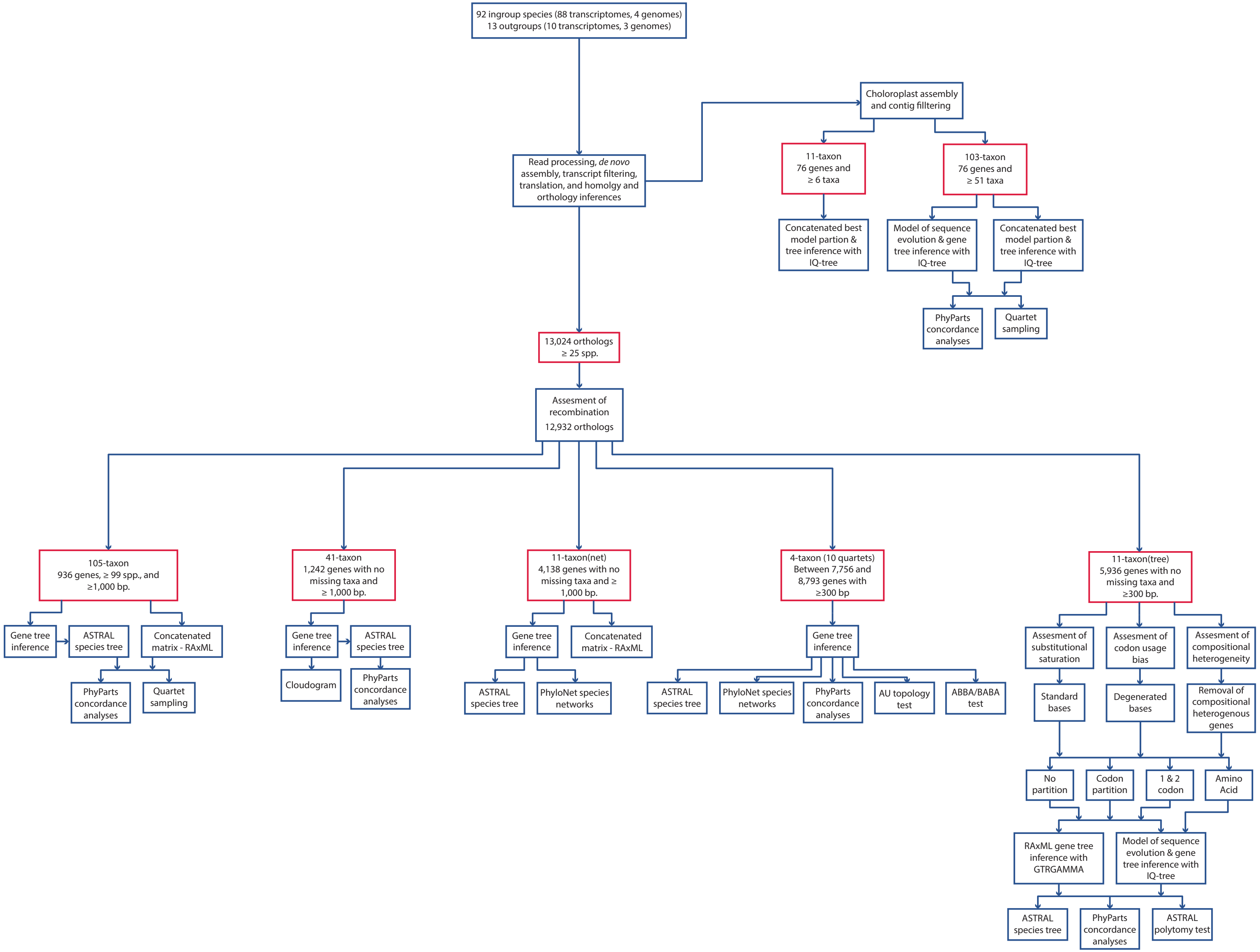


**FIGURE S1**. Overview of all datasets (red boxes) and analyses (blue boxes and arrows) in this study.

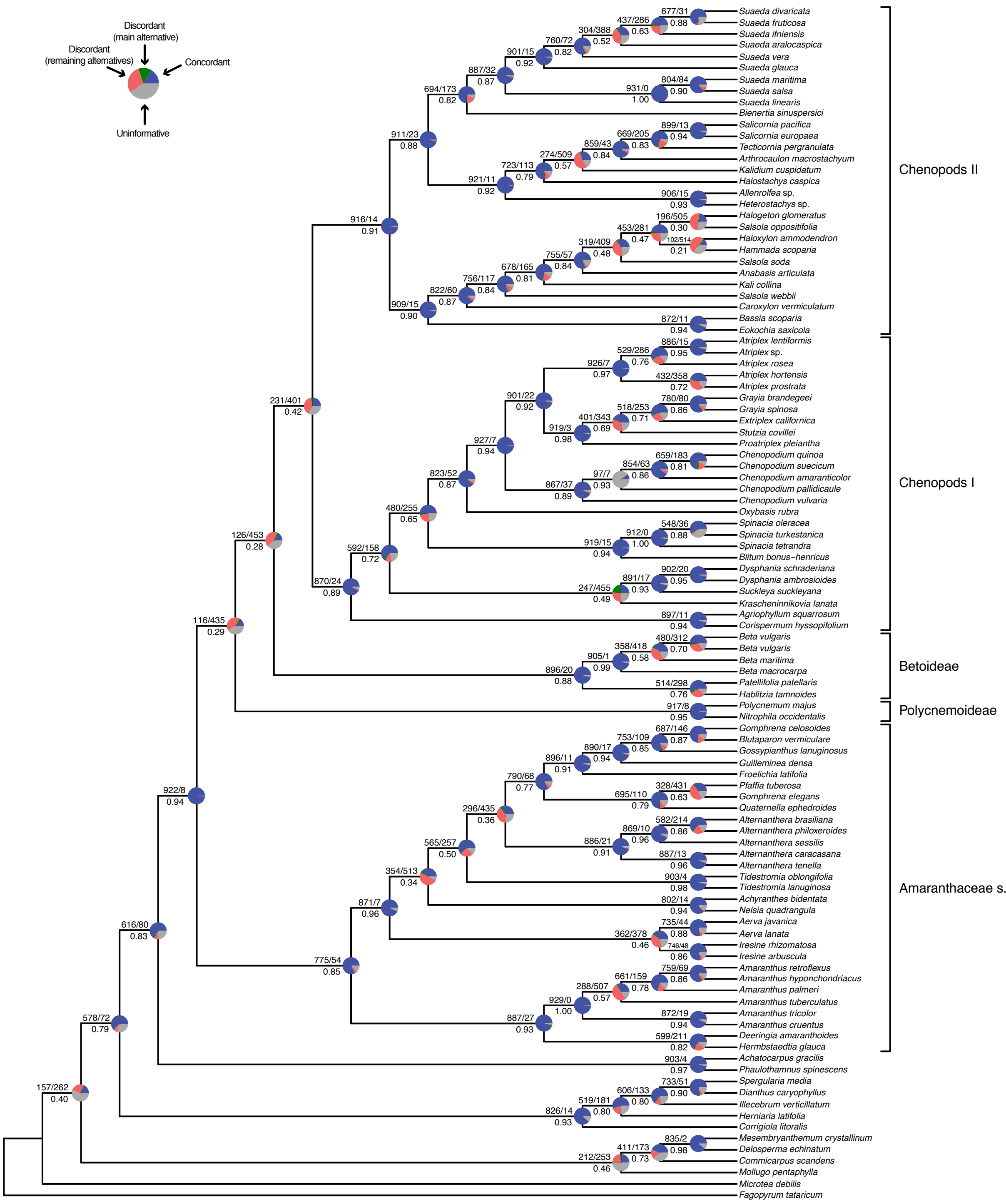


**FIGURE S2.** Maximum likelihood cladogram of Amaranthaceae s.l. inferred from RAxML analysis of the concatenated 936-nuclear gene supermatrix. Numbers above branches indicate the number of gene trees concordant/conflicting with that node in the species tree. Numbers below the branches are the Internode Certainty All (ICA) score. Pie charts on nodes present the proportion of gene trees that support that clade (blue), the proportion that support the main alternative bifurcation (green), the proportion that support the remaining alternatives (red), and the proportion (conflict or support) that have < 50% bootstrap support (gray).

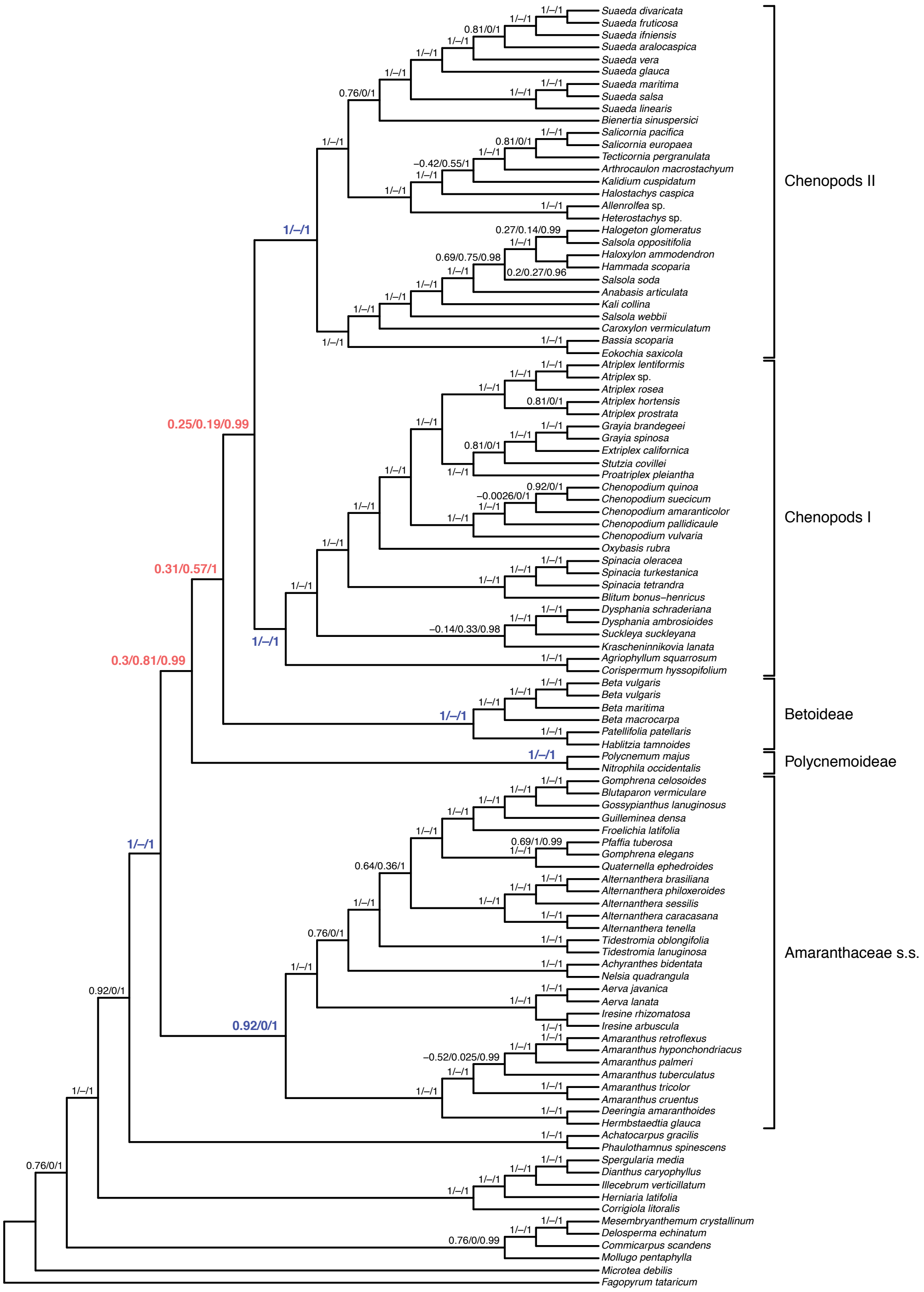


**FIGURE S3.** Maximum likelihood cladogram of Amaranthaceae s.l. inferred from RAxML analysis of the concatenated 936-nuclear gene supermatrix. Numbers above branches indicate the Quartet sampling internal node score. Quartet concordance/Quartet differential/ Quartet informativeness. Scores in blue indicate strong support for the species tree topology, while red scores indicate strong support for alternative topologies.

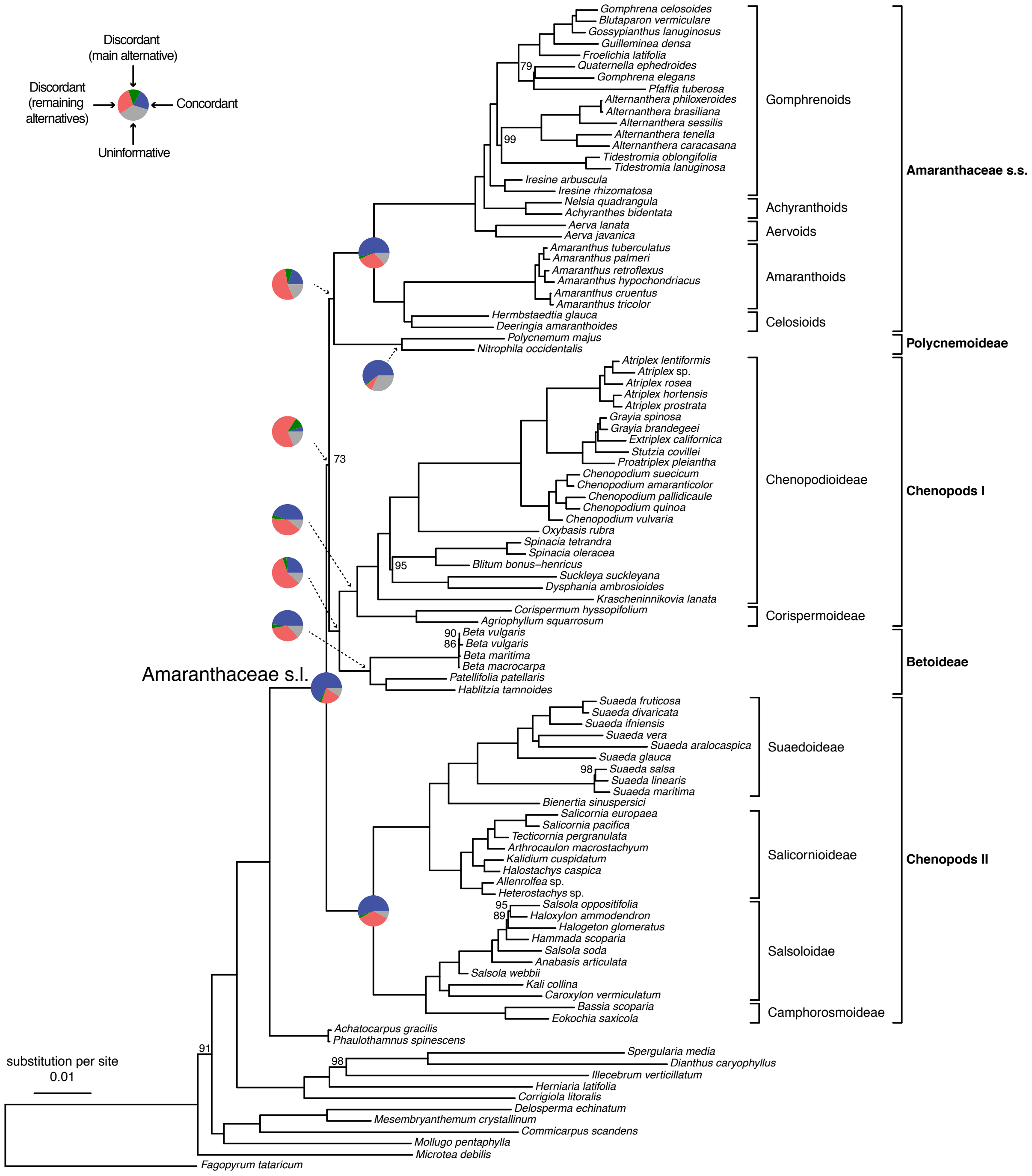


**FIGURE S4.** Maximum likelihood phylogeny of Amaranthaceae s.l. inferred from IQ-tree analysis of concatenated 76-plastid gene supermatrix. All nodes have full support (Bootstrap = 100) unless noted next to nodes. Pie charts present the proportion of gene trees that support that clade (blue), the proportion that support the main alternative bifurcation (green), the proportion that support the remaining alternatives (red), and the proportion (conflict or support) that have < 50% bootstrap support (gray). Only pie charts for major clades are shown (see Fig. S5 for all node pie charts). Branch lengths are in number of substitutions per site.

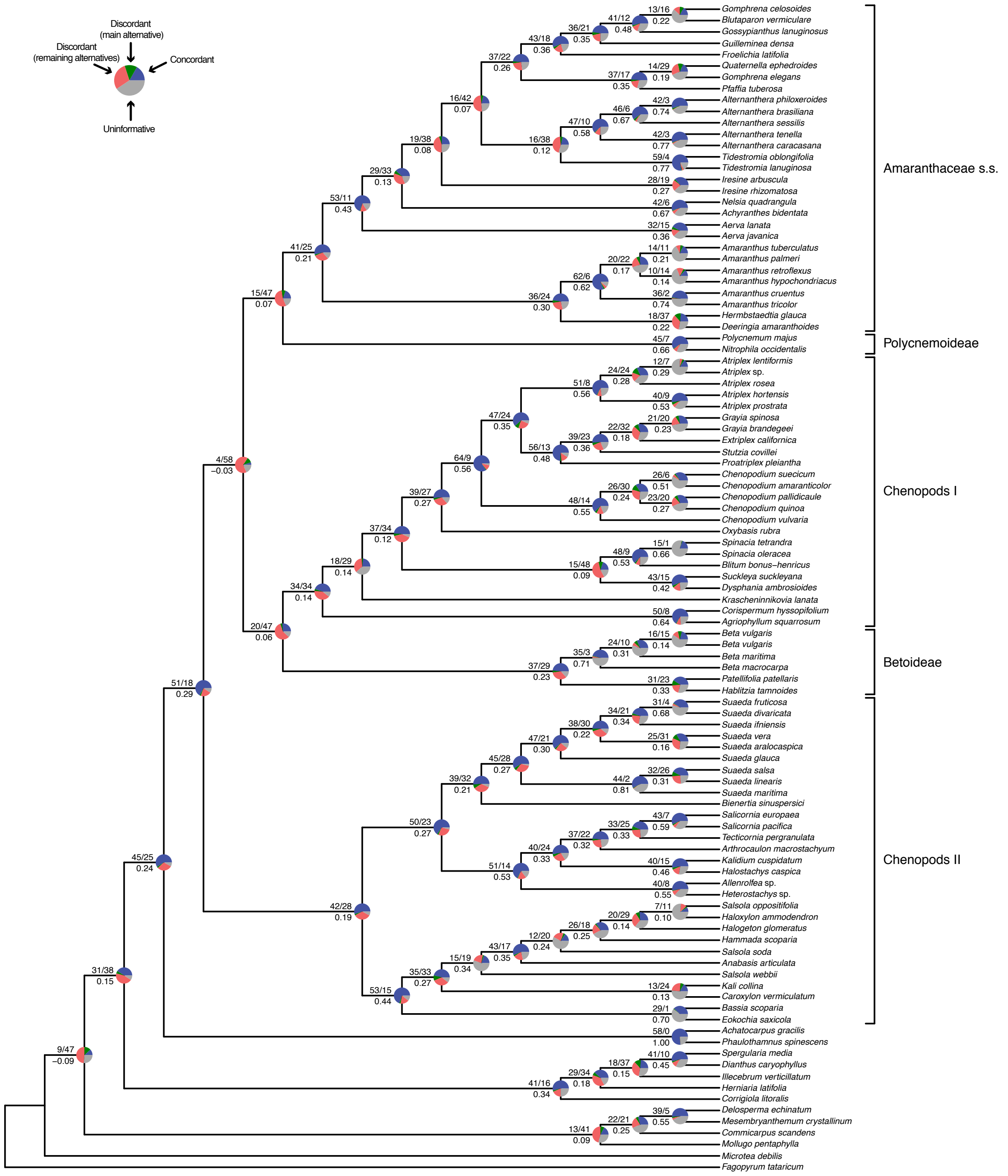


**FIGURE S5.** Maximum likelihood cladogram of Amaranthaceae s.l. inferred from IQ-tree analysis of concatenated 76-plastid gene supermatrix. Numbers above branches indicate the number of gene trees concordant/conflicting with that node in the species tree. Numbers below the branches are the Internode Certainty All (ICA) score. Pie charts on nodes present the proportion of gene trees that support that clade (blue), the proportion that support the main alternative bifurcation (green), the proportion that support the remaining alternatives (red), and the proportion (conflict or support) that have < 50% bootstrap support (gray).

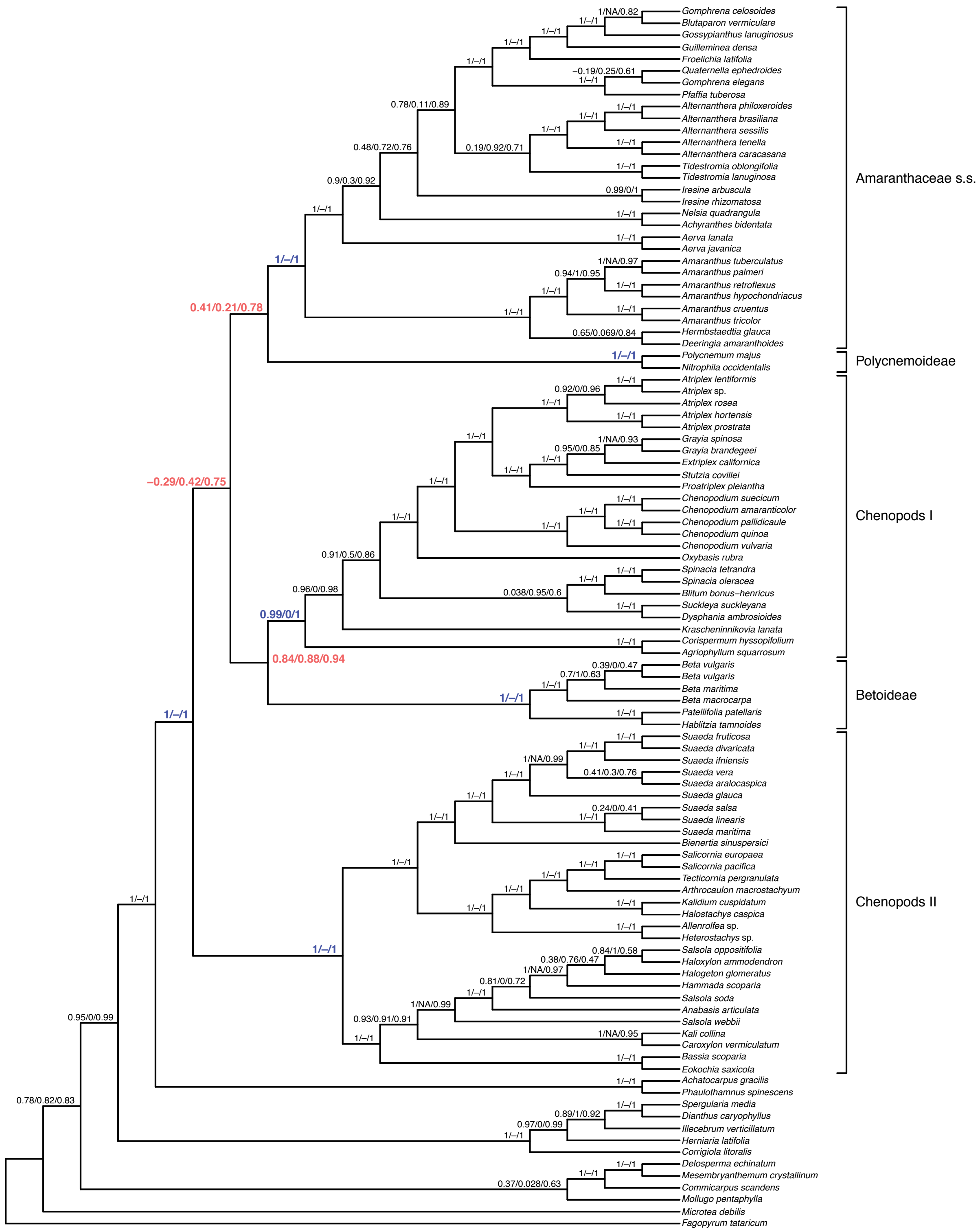

**FIGURE S6.** Maximum likelihood cladogram of Amaranthaceae s.l. inferred from IQ-tree analysis of concatenated 76-plastid gene supermatrix. Numbers above branches indicate the Quartet sampling internal node score. Quartet concordance/Quartet differential/ Quartet informativeness. Scores in blue indicate strong support for the species tree topology, while red scores indicate strong support for alternative topologies.

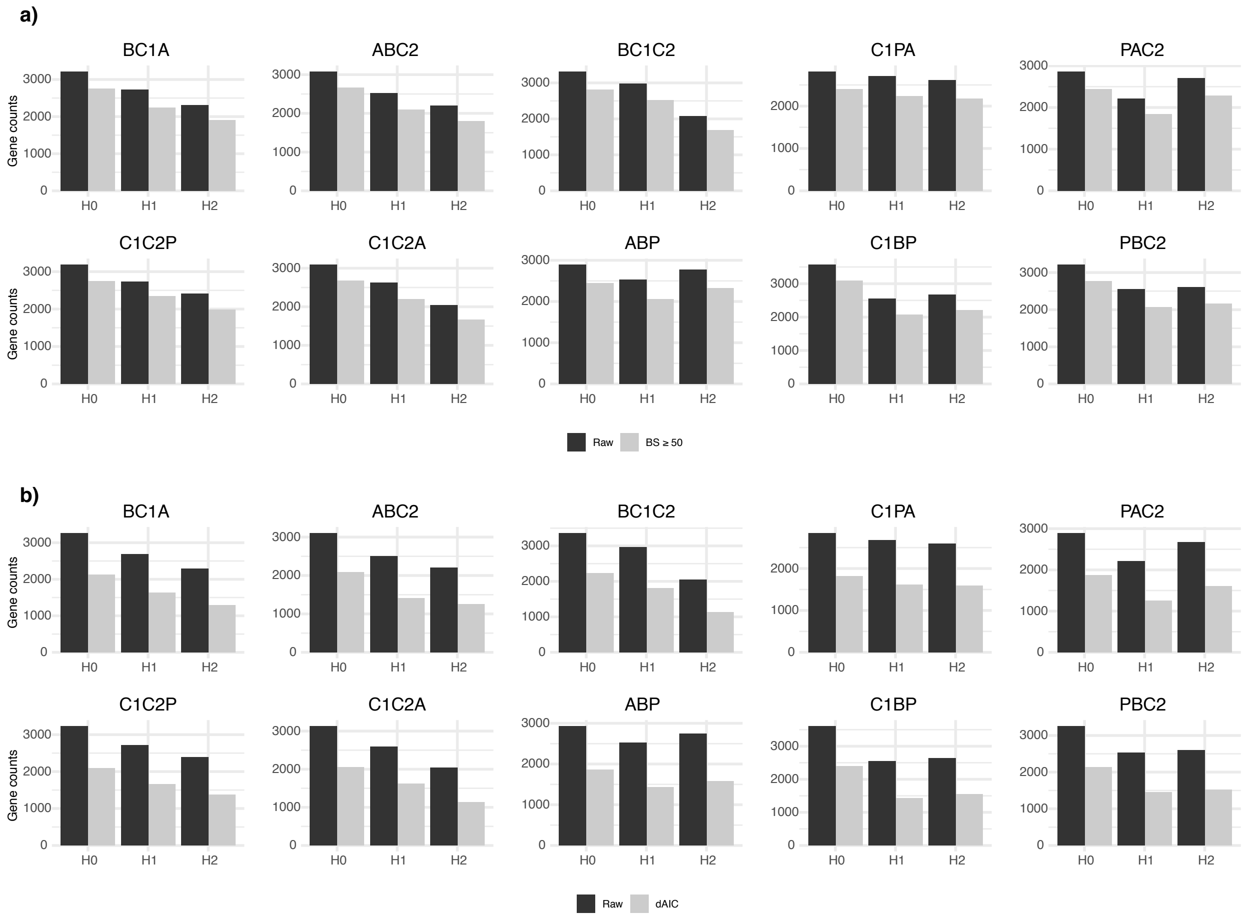


**FIGURE S7.** Counts of gene tree inferences for the 10 quartets from the five main clades of Amaranthaceae s.l. (a) Bars represent raw gene tree counts and counts of gene trees with bootstrap support ≥ 50. Gene counts of constrained maximum likelihood searches for the 10 quartets from the five main clades of Amaranthaceae s.l. (b) Bars represent counts based on raw maximum likelihood scores and counts based on trees with significant support (trees with a delta corrected Akaike Information Criteria (ΔAICc) ≥ 2 than the next best model). H0 represents the ASTRAL species tree of each quartet inferred. Each quartet is named following the species tree topology, where the first two species are sister to each other (all topologies can be found in Figure S6). A = Amaranthaceae. s.s. (*Amaranthus hypochondriacus*), B = Betoideae (*Beta vulgaris*), C1 = Chenopods I (*Chenopodium quinoa*), C2 = Chenopods II (*Caroxylum vermiculatum*), P = Polycnemoideae (*Polycnemum majus*).

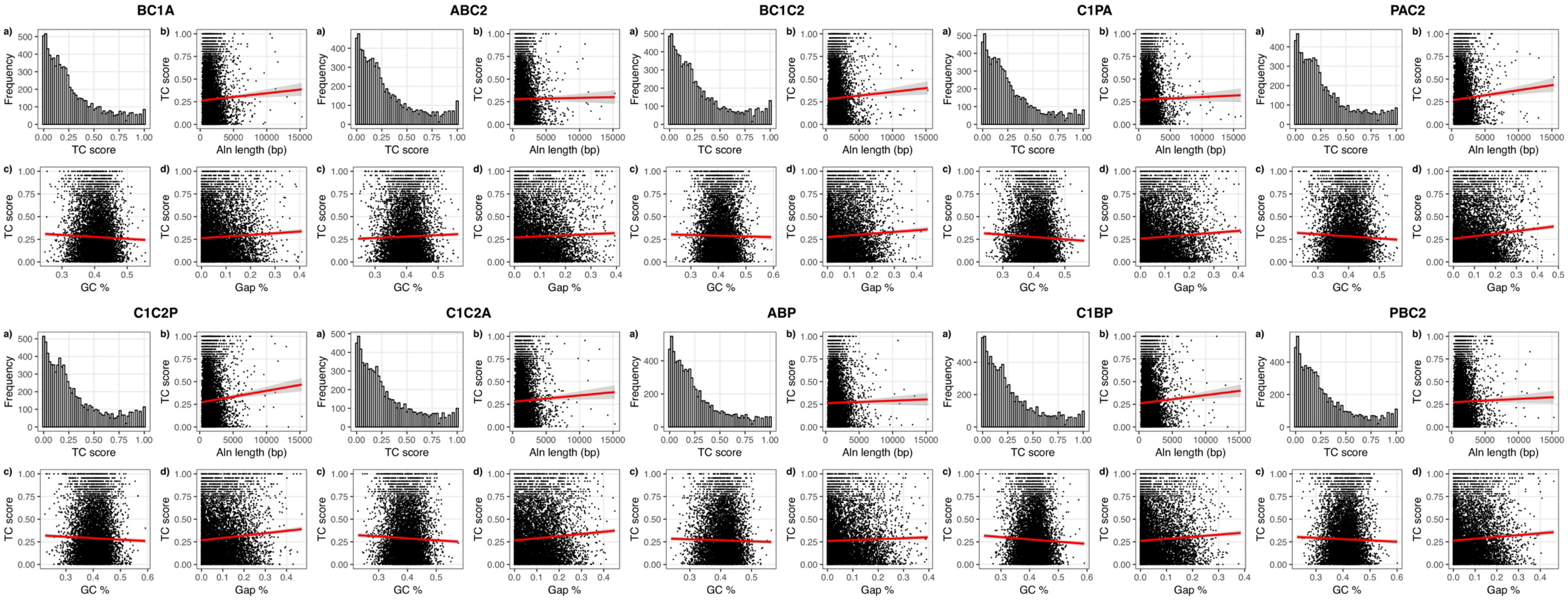


**FIGURE S8.** Alignment and tree scores for each of the 10 quartets from the five main clades of Amaranthaceae s.l. Quartet Tree Certainty (TC) score distribution (a) and its correlation with alignment length (b), alignment GC content (c), and alignment gap percentage (d). Quartets named following the ASTRAL species tree topology (see Figure 6 for quartet topologies). A = Amaranthaceae. s.s. (*Amaranthus hypochondriacus*), B = Betoideae (*Beta vulgaris*), C1 = Chenopods I (*Chenopodium quinoa*), C2 = Chenopods II (*Caroxylum vermiculatum*), P = Polycnemoideae (*Polycnemum majus*).

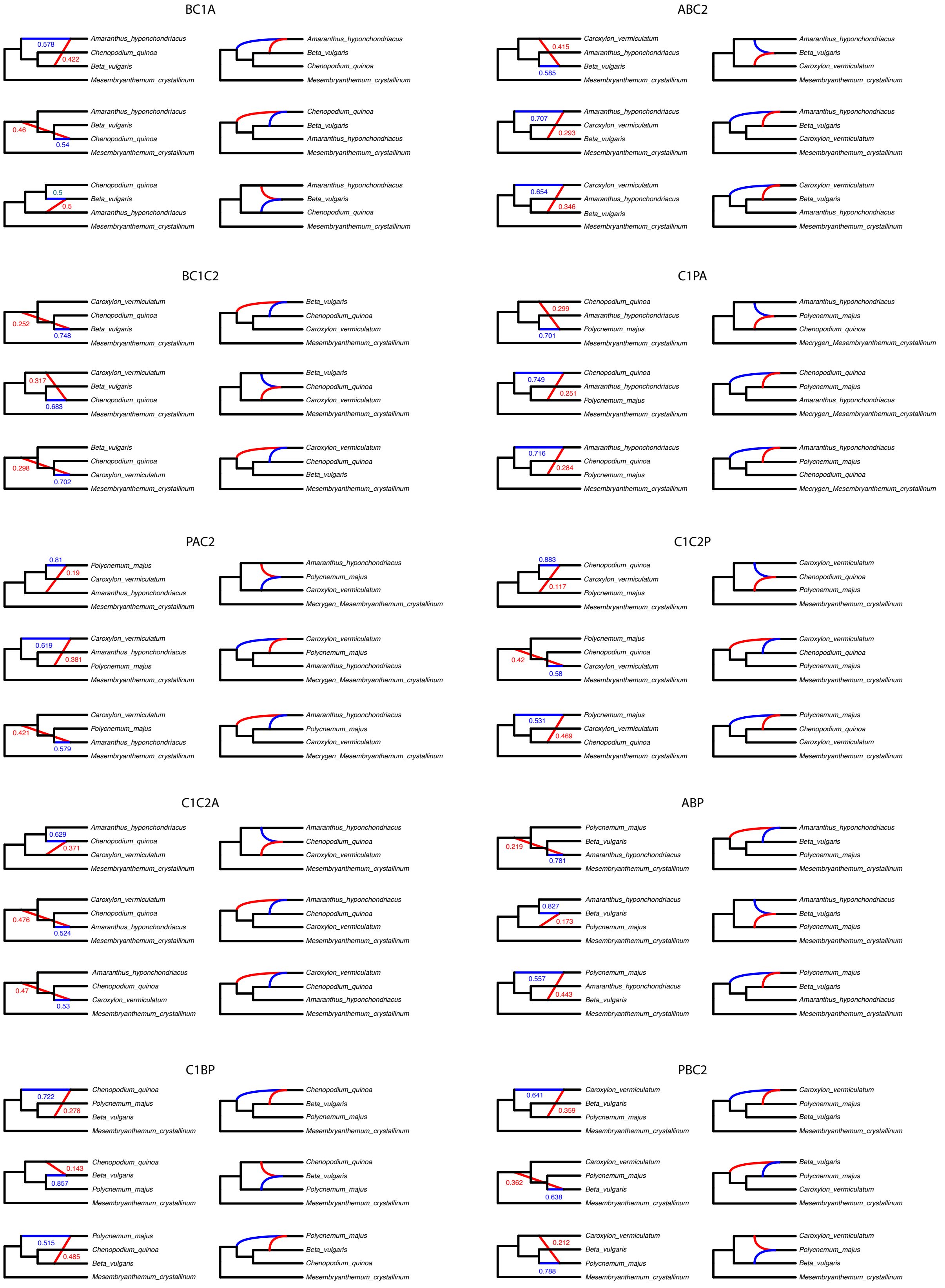


**FIGURE S9.** Best three species networks for the 10 quartets from the five main clades of Amaranthaceae s.l. Red and blue indicates the minor and major edges, respectively, of hybrid nodes. Number next to the branches indicates inheritance probabilities for each hybrid node. Network visualization with PhyloPlots (left) and network visualization with Dendroscope (right). Each quartet is named following the species tree topology, where the first two are sisters (all topologies can be found in Figure S6). A = Amaranthaceae. s.s. (*Amaranthus hypochondriacus*), B = Betoideae (*Beta vulgaris*), C1 = Chenopods I (*Chenopodium quinoa*), C2 = Chenopods II (*Caroxylum vermiculatum*), P = Polycnemoideae (*Polycnemum majus*).

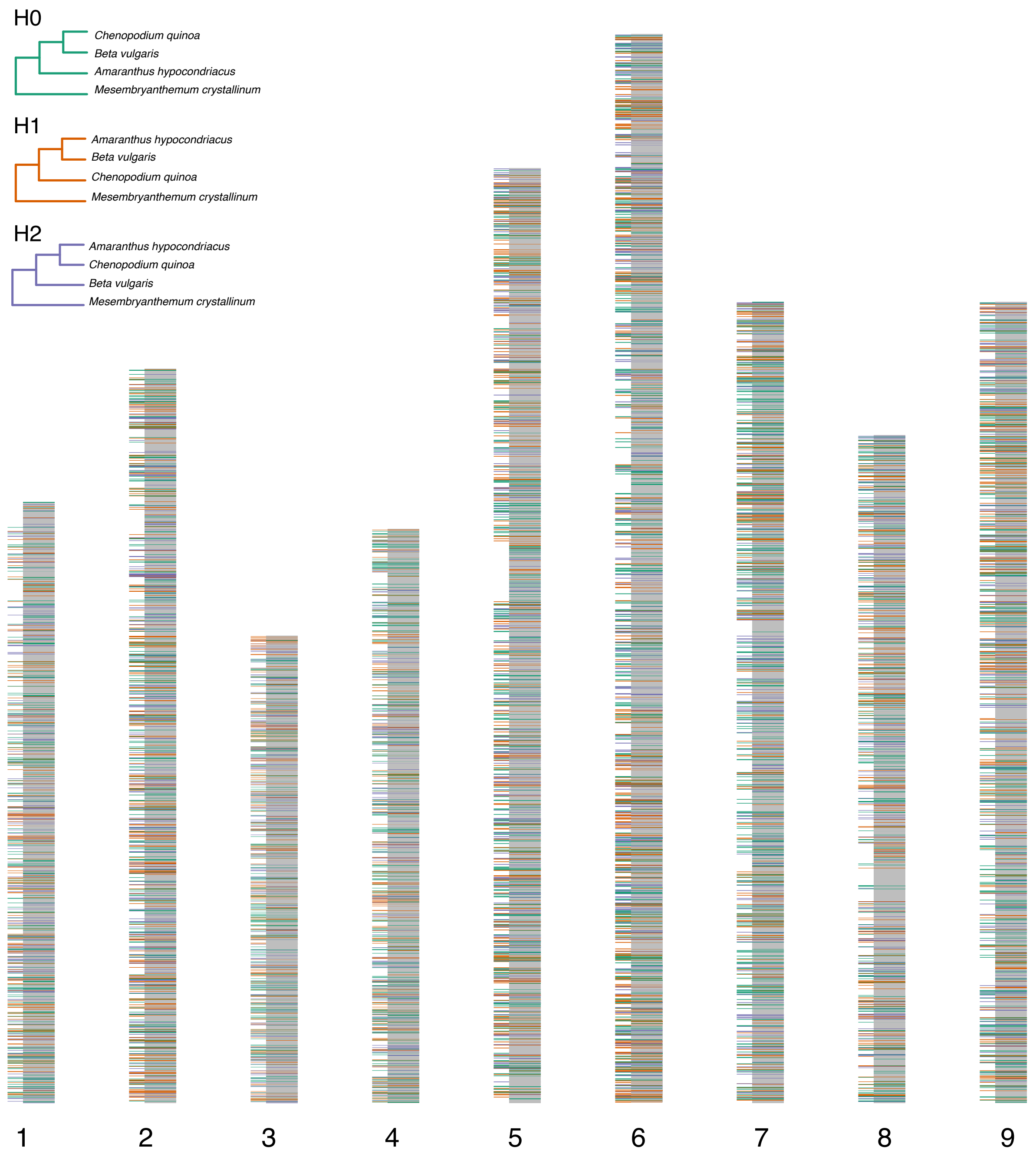


**FIGURE S10.** Chromosomes of *Beta vulgaris* with gene tree topologies from the quartet BC1A mapped. Each gene is colored according to gene tree topology (H0, H1, or H2). H0 represents the ASTRAL species tree of the quartet. Longer colored lines along the chromosomes represent syntenic genes (6,941) between *Beta vulgaris* and *Mesembryanthemum crystallinum.* Grey represents genes not present in the ortholog set of the quartet BC1A. A = Amaranthaceae. s.s. (*Amaranthus hypochondriacus*), B = Betoideae (*Beta vulgaris*), C1 = Chenopods I (*Chenopodium quinoa*), C2 = Chenopods II (*Caroxylum vermiculatum*), P = Polycnemoideae (*Polycnemum majus*).

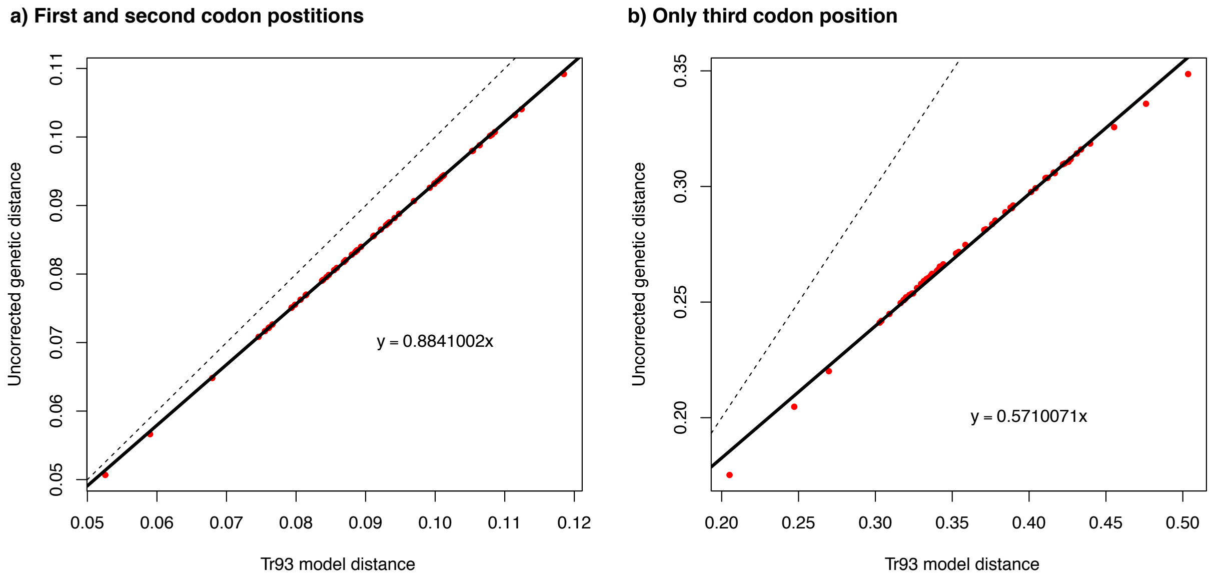


**FIGURE S11.** Saturation plots from the 11-taxon(tree) concatenated alignment. a) Saturation analysis for the first and second codon positions, and b) saturation analyses for the third codon position. Dotted lines represent expected unsaturated curves, and solid lines represent the observed saturation curves.

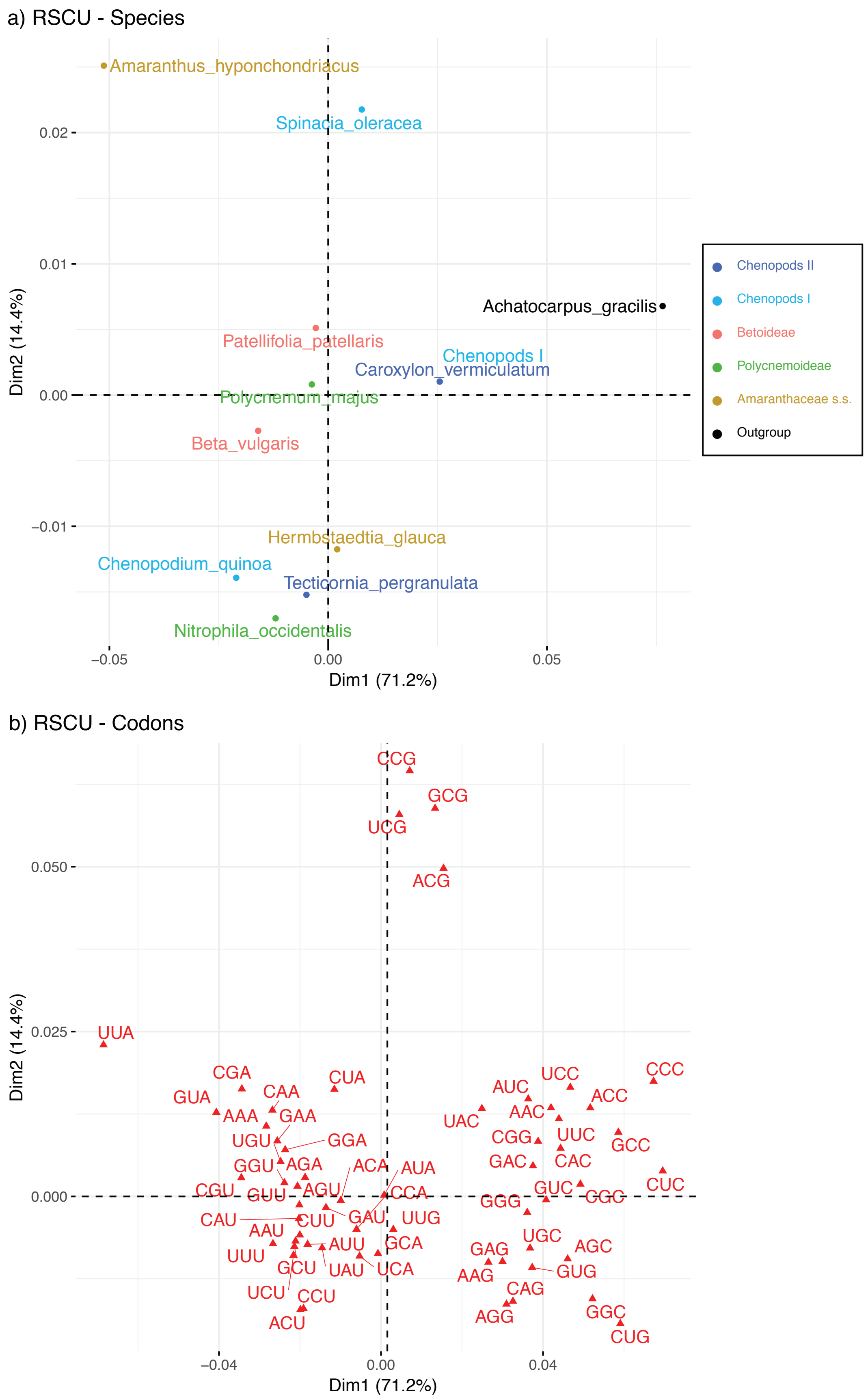


**FIGURE S12.** Correspondence analyses results of the Synonymous Codon Usage (RSCU) of the 11-taxon(tree) concatenated alignment. Results by a) species and (b) codons.

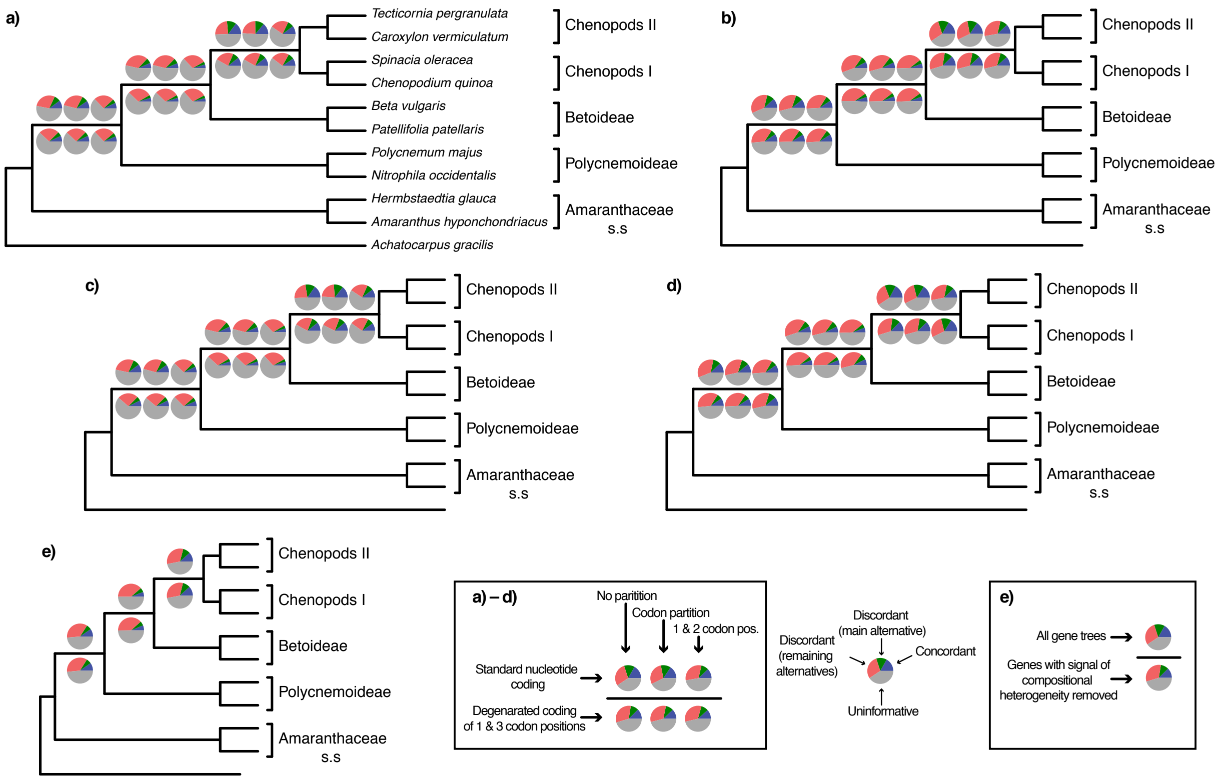


**FIGURE S13.** ASTRAL species trees from the 11-taxon(net) dataset estimated from gene trees inferred using multiple data schemes. a) Gene trees inferred with RAxML with a GTR-GAMMA model. b) Gene trees inferred with IQ-tree allowing for automatic model selection of sequence evolution. c) Gene trees inferred with RAxML with a GTR-GAMMA model and removal of genes that had signal of compositional heterogeneity. d) Gene trees inferred with IQ-tree allowing for automatic model selection of sequence evolution and removal of genes that had signal of compositional heterogeneity. a–d) Gene trees were inferred with no partition, codon partition (first and second codon, and third codon) and, only first and second codon positions (third codon position removed and no partition). Gene trees were inferred using codon alignments with standard nucleotide coding, and alignments with degenerated coding of the first and third codon positions. e) All gene trees and gene trees after removal of genes that had signal of compositional heterogeneity, inferred with IQ-tree using amino acid sequences allowing for automatic model selection of sequence evolution. Pie charts on nodes present the proportion of gene trees that support that clade (blue), the proportion that support the main alternative bifurcation (green), the proportion that support the remaining alternatives (red), and the proportion (conflict or support) that have < 50% bootstrap support (gray).

**Table S1.** Taxon sampling and source of data

| Family | Subfamily | Species code | Genus | Species | Authority name | Taxon name in tree files and intermediate files | Library reads single/paired-end | Library stranded? | No. of final CDS used for BLASTN | Reference coverage (*Beta vulgaris*) | No. MO orthologs | Perc. MO orthologs (total 13024) | Usage in previous phylotranscriptomic papers | Source database | Source accession | Souce reference | Souce title |
|---|---|---|---|---|---|---|---|---|---|---|---|---|---|---|---|---|---|
| Amaranthaceae s.l. | Betoideae | Betamac | *Beta* | *macrocarpa* | Guss. | Betamac_Beta_macrocarpa | Paired | Non-stranded | 17741 | 0.54 | 10279 | 78.9% | --- | SRA | SRR1038481 | Fan et al. 2015 | Transcriptome Analysis of Beta macrocarpa and Identification of Differentially Expressed Transcripts in Response to Beet Necrotic Yellow Vein Virus Infection |
| Amaranthaceae s.l. | Betoideae | FVXD | *Beta* | *maritima* | L. | FVXD_Beta_maritima | Paired | Non-stranded | 15017 | 0.23 | 7774 | 59.7% | used in Yang et. al 2015 | SRA | ERR2040223 | 1KP/Matasci et al., 2014 | Data access for the 1,000 Plants (1KP) project |
| Amaranthaceae s.l. | Betoideae | Betatrig | *Beta* | *vulgaris* | L. | Betatrig_Beta_trigina[a] | Paired | Stranded | 17048 | 0.53 | 10662 | 81.9% | --- | Newly sequenced | Newly sequenced | Newly sequenced | Newly sequenced |
| Amaranthaceae s.l. | Betoideae | Betavulgen | *Beta* | *vulgaris* | L. | Bevulgen_Beta_vulgaris | --- | --- | 26922 | --- | 11753 | 90.2% | --- | http://bvseq.molgen.mpg.de/index.shtml | v.1.2 | Dohm et al. 2013 | The genome of the recently domesticated crop plant sugar beet (Beta vulgaris) |
| Amaranthaceae s.l. | Betoideae | Habtam | *Hablitzia* | *tamnoides* | M. Bieb. | Habtam_Hablitzia_tamnoides | Paired | Stranded | 15735 | 0.44 | 9809 | 75.3% | --- | Newly sequenced | Newly sequenced | Newly sequenced | Newly sequenced |
| Amaranthaceae s.l. | Betoideae | Patpat | *Patellifolia* | *patellaris* | (Moq.) A.J. Scott, Ford-Lloyd & J.T. Williams | Patpat_Patellifolia_patellaris | Paired | Stranded | 22777 | 0.53 | 11040 | 84.8% | --- | Newly sequenced | Newly sequenced | Newly sequenced | Newly sequenced |
| Amaranthaceae s.l. | Polycnemoideae | MJM3242 | *Nitrophila* | *occidentalis* | (Moq.) S. Watson | MJM3242_Nitrophila_occidentalis | Paired | Stranded | 39370 | 0.55 | 11649 | 89.4% | used in Walker et al. 2018 | SRA | SRR6435357 | Walker et al. 2018 | From cacti to carnivores: Improved phylotranscriptomic sampling and hierarchical homology inference provide further insight into the evolution of Caryophyllales |
| Amaranthaceae s.l. | Polycnemoideae | PolmaSFB | *Polycnemum* | *majus* | A. Braun ex Bogenh. | PolmaSFB_Polycnemum_majus | Paired | Stranded | 42557 | 0.57 | 11668 | 89.6% | used in Walker et al. 2018 | SRA | SRR6435356 | Walker et al. 2018 | From cacti to carnivores: Improved phylotranscriptomic sampling and hierarchical homology inference provide further insight into the evolution of Caryophyllales |
| Amaranthaceae s.s. | Achyranthoids | Achbid | *Achyranthes* | *bidentata* | Blume | Achbid_Achyranthes_bidentata | Paired | Non-stranded | 25583 | 0.5 | 8598 | 66.0% | --- | SRA | SRR5591716 | Yang et al. 2018 | The root transcriptome of Achyranthes bidentata and the identification of the genes involved in the replanting benefit |
| Amaranthaceae s.s. | Achyranthoids | NequaSFB | *Nelsia* | *quadrangula* | Schinz | NequaSFB_Nelsia_quadrangula | Paired | Stranded | 24387 | 0.54 | 9313 | 71.5% | used in Walker et al. 2018 | SRA | SRR6435358 | Walker et al. 2018 | From cacti to carnivores: Improved phylotranscriptomic sampling and hierarchical homology inference provide further insight into the evolution of Caryophyllales |
| Amaranthaceae s.s. | Aervoids | HDSY | *Aerva* | *javanica* | (Burm. f.) Juss. | HDSY_Aerva_javanica | Paired | Non-stranded | 27385 | 0.43 | 6929 | 53.2% | used in Yang et. al 2015 | SRA | ERR2040202 /ERR2040203 | 1KP/Matasci et al., 2014 | Data access for the 1,000 Plants (1KP) project |
| Amaranthaceae s.s. | Aervoids | PDQH | *Aerva* | *lanata* | (L.) Juss. ex Schult. | PDQH_Aerva_lanata | Paired | Non-stranded | 19501 | 0.41 | 6879 | 52.8% | used in Yang et. al 2015 | SRA | ERR2040200 /ERR2040201 | 1KP/Matasci et al., 2014 | Data access for the 1,000 Plants (1KP) project |
| Amaranthaceae s.s. | Aervoids | MJM2678 | *Iresine* | *arbuscula* | Uline & W.L. Bray | MJM2678_Iresine_arbuscula | Paired | Stranded | 26935 | 0.53 | 7997 | 61.4% | used in Walker et al. 2018 | SRA | SRR6435327 | Walker et al. 2018 | From cacti to carnivores: Improved phylotranscriptomic sampling and hierarchical homology inference provide further insight into the evolution of Caryophyllales |
| Amaranthaceae s.s. | Aervoids | MJM2943 | *Iresine* | *rhizomatosa* | Standl. | MJM2943_Iresine_rhizomatosa | Paired | Stranded | 21890 | 0.53 | 7571 | 58.1% | used in Walker et al. 2018 | SRA | SRR6435359 | Walker et al. 2018 | From cacti to carnivores: Improved phylotranscriptomic sampling and hierarchical homology inference provide further insight into the evolution of Caryophyllales |
| Amaranthaceae s.s. | Amaranthoids | XSSD | *Amaranthus* | *cruentus* | L. | XSSD_Amaranthus_cruentus | Paired | Non-stranded | 22359 | 0.48 | 10215 | 78.4% | used in Yang et. al 2015 | SRA | ERR2040205 | 1KP/Matasci et al., 2014 | Data access for the 1,000 Plants (1KP) project |
| Amaranthaceae s.s. | Amaranthoids | Amhy | *Amaranthus* | *hypochondriacus* | L. | Amhypgen_Amaranthus_hyponchondriacus | --- | --- | 23843 | --- | 10612 | 81.5% | --- | Phytozome | v1.0 | Lightfoot et al. 2017 | Single-molecule sequencing and Hi-C-based proximity-guided assembly of amaranth (Amaranthus hypochondriacus) chromosomes provide insights into genome evolution. |
| Amaranthaceae s.s. | Amaranthoids | Ampal | *Amaranthus* | *palmeri* | S. Watson | Ampal_Amaranthus_palmeri | Paired | Non-stranded | 23336 | 0.5 | 9960 | 76.5% | --- | SRA | SRR5759382 | Salas-Perez et al. 2018 | RNA-Seq transcriptome analysis of Amaranthus palmeri with differential tolerance to glufosinate herbicide. |
| Amaranthaceae s.s. | Amaranthoids | WMLW | *Amaranthus* | *retroflexus* | L. | WMLW_Amaranthus_retroflexus | Paired | Non-stranded | 19417 | 0.29 | 7859 | 60.3% | used in Yang et. al 2015 | SRA | ERR2040206 | 1KP/Matasci et al., 2014 | Data access for the 1,000 Plants (1KP) project |
| Amaranthaceae s.s. | Amaranthoids | Amtric | *Amaranthus* | *tricolor* | L. | Amtric_Amaranthus_tricolor | Paired | Non-stranded | 24201 | 0.5 | 9933 | 76.3% | --- | SRA | SRR5930345 | Liu et al. 2018 | Identification of microRNAs in the green and red sectors of Amaranthus tricolor L. leaves based on Illumina sequencing data |
| Amaranthaceae s.s. | Amaranthoids | Amtub | *Amaranthus* | *tuberculatus* | (Moq.) J.D. Sauer | Amtub_Amaranthus_tuberculatus | Paired | Non-stranded | 22363 | 0.48 | 9965 | 76.5% | --- | SRA | SRR6705609 | --- | Submitted to SRA in 2018. No reference found |
| Amaranthaceae s.s. | Celosioids | Deeamar | *Deeringia* | *amaranthoides* | (Lam.) Merr. | Deeamar_Deeringia_amaranthoides | Paired | Stranded | 24155 | 0.52 | 10833 | 83.2% | --- | Newly sequenced | Newly sequenced | Newly sequenced | Newly sequenced |
| Amaranthaceae s.s. | Celosioids | Hergla | *Hermbstaedtia* | *glauca* | Reichb. ex Steud. | Hergla_Hermbstaedtia_glauca | Paired | Stranded | 24119 | 0.54 | 11212 | 86.1% | --- | Newly sequenced | Newly sequenced | Newly sequenced | Newly sequenced |
| Amaranthaceae s.s. | Gomphrenoids | ZBPY | *Alternanthera* | *brasiliana* | (L.) Kuntze | ZBPY_Alternanthera_brasiliana | Paired | Non-stranded | 39746 | 0.47 | 9141 | 70.2% | used in Yang et. al 2015 | SRA | ERR2040215/ERR2040216 | 1KP/Matasci et al., 2014 | Data access for the 1,000 Plants (1KP) project |
| Amaranthaceae s.s. | Gomphrenoids | OHKC | *Alternanthera* | *caracasana* | Kunth | OHKC_Alternanthera_caracasana | Paired | Non-stranded | 41349 | 0.41 | 8905 | 68.4% | used in Yang et. al 2015 | SRA | ERR2040217/ERR2040218 | 1KP/Matasci et al., 2014 | Data access for the 1,000 Plants (1KP) project |
| Amaranthaceae s.s. | Gomphrenoids | Alph | *Alternanthera* | *philoxeroides* | (Mart.) Griseb. | Alph_Alternanthera_philoxeroides | Paired | Non-stranded | 51167 | 0.52 | 9726 | 74.7% | --- | SRA | SRR1661509 | Liu et al. 2019 | RNA sequencing characterizes transcriptome differences in cold response between northern and southern Alternanthera philoxeroides and highlight adaptations associated with northward expansion |
| Amaranthaceae s.s. | Gomphrenoids | BWRK | *Alternanthera* | *sessilis* | (L.) R. Br. ex DC. | BWRK_Alternanthera_sessilis | Paired | Non-stranded | 23148 | 0.42 | 8664 | 66.5% | used in Yang et. al 2015 | SRA | ERR2040219 /ERR2040220 | 1KP/Matasci et al., 2014 | Data access for the 1,000 Plants (1KP) project |
| Amaranthaceae s.s. | Gomphrenoids | EYRD | *Alternanthera* | *tenella* | Colla | EYRD_Alternanthera_tenella | Paired | Non-stranded | 25884 | 0.4 | 8909 | 68.4% | used in Yang et. al 2015 | SRA | ERR2040221/ERR2040222 | 1KP/Matasci et al., 2014 | Data access for the 1,000 Plants (1KP) project |
| Amaranthaceae s.s. | Gomphrenoids | CUTE | *Blutaparon* | *vermiculare* | (L.) Mears | CUTE_Blutaparon_vermiculare | Paired | Non-stranded | 18831 | 0.39 | 8661 | 66.5% | used in Yang et. al 2015 | SRA | ERR2040207 | 1KP/Matasci et al., 2014 | Data access for the 1,000 Plants (1KP) project |
| Amaranthaceae s.s. | Gomphrenoids | MJM1665 | *Froelichia* | *latifolia* | R.A. McCauley | MJM1665_Froelichia_latifolia | Paired | Stranded | 30906 | 0.46 | 9395 | 72.1% | used in Yang et. al 2018 | SRA | SRR1979685 | Brockington et al 2015 | Lineage-specific gene radiations underlie the evolution of novel betalain pigmentation in Caryophyllales |
| Amaranthaceae s.s. | Gomphrenoids | Gomcel | *Gomphrena* | *celosioides* | Mart. | Gomcel_Gomphrena_celosoides[a] | Paired | Stranded | 21652 | 0.5 | 10127 | 77.8% | --- | Newly sequenced | Newly sequenced | Newly sequenced | Newly sequenced |
| Amaranthaceae s.s. | Gomphrenoids | Gomele | *Gomphrena* | *elegans* | Mart. | Gomele_Gomphrena_elegans | Paired | Stranded | 26535 | 0.52 | 10147 | 77.9% | --- | Newly sequenced | Newly sequenced | Newly sequenced | Newly sequenced |
| Amaranthaceae s.s. | Gomphrenoids | MJM1807 | *Gossypianthus* | *lanuginosus* | (Poir.) Moq. | MJM1807_Gossypianthus_lanuginosus | Paired | Stranded | 19778 | 0.54 | 10319 | 79.2% | used in Yang et. al 2018 | SRA | SRR6787476 | Yang et al. 2018 | Improved transcriptome sampling pinpoints 26 ancient and more recent polyploidy events in Caryophyllales, including two allopolyploidy events |
| Amaranthaceae s.s. | Gomphrenoids | MJM2445 | *Guilleminea* | *densa* | (Humb. & Bonpl. ex Schult.) Moq. | MJM2445_Guilleminea_densa | Paired | Stranded | 26558 | 0.53 | 10237 | 78.6% | used in Yang et. al 2018 | SRA | SRR6787501 | Yang et al. 2018 | Improved transcriptome sampling pinpoints 26 ancient and more recent polyploidy events in Caryophyllales, including two allopolyploidy events |
| Amaranthaceae s.s. | Gomphrenoids | Pfatub | *Pfaffia* | *tuberosa* | (Spreng.) Hicken | Pfatub_Pfaffia_tuberosa | Paired | Stranded | 26649 | 0.5 | 9798 | 75.2% | --- | Newly sequenced | Newly sequenced | Newly sequenced | Newly sequenced |
| Amaranthaceae s.s. | Gomphrenoids | Quaephe | *Quaternella* | *ephedroides* | Pedersen | Quaephe_Quaternella_ephedroides | Paired | Stranded | 23950 | 0.47 | 9557 | 73.4% | --- | Newly sequenced | Newly sequenced | Newly sequenced | Newly sequenced |
| Amaranthaceae s.s. | Gomphrenoids | MJM2259 | *Tidestromia* | *lanuginosa* | (Nutt.) Standl. | MJM2259_Tidestromia_lanuginosa | Paired | Stranded | 24837 | 0.57 | 9779 | 75.1% | used in Yang et. al 2018 | SRA | SRR6787504 | Yang et al. 2018 | Improved transcriptome sampling pinpoints 26 ancient and more recent polyploidy events in Caryophyllales, including two allopolyploidy events |
| Amaranthaceae s.s. | Gomphrenoids | Tisp1013 | *Tidestromia* | *oblongifolia* | (S. Watson) Standl. | Tisp1013_Tidestromia_suffruticosa[a] | Paired | Stranded | 22679 | 0.51 | 9294 | 71.4% | --- | Newly sequenced | Newly sequenced | Newly sequenced | Newly sequenced |
| Chenopodiaceae | -- | Bisi | *Bienertia* | *sinuspersici* | Akhani | Bisi_Bienertia_sinuspersici | --- | --- | 42004 | 0.43 | 9401 | 72.2% | used in Yang et. al 2018 | Not assembled. CDS from Yang et al. 2017 | SRR1770364 | Sharpe, 2014 | Gene expression profiling in single cell c4 and related photosynthetic species in Suaedoideae |
| Chenopodiaceae | Camphorosmoideae | WGET | *Bassia* | *scoparia* | (L.) A.J. Scott | WGET_Bassia_scoparia | Paired | Non-stranded | 16019 | 0.3 | 8665 | 66.5% | used in Yang et. al 2015 | SRA | ERR364385 | 1KP/Matasci et al., 2014 | Data access for the 1,000 Plants (1KP) project |
| Chenopodiaceae | Camphorosmoideae | EosaxSFB | *Eokochia* | *saxicola* | (Guss.) Freitag & G. Kadereit | EosaxSFB_Eokochia_saxicola | Paired | Stranded | 29936 | 0.55 | 11193 | 85.9% | used in Walker et al. 2018 | SRA | SRR6435348 | Walker et al. 2018 | From cacti to carnivores: Improved phylotranscriptomic sampling and hierarchical homology inference provide further insight into the evolution of Caryophyllales |
| Chenopodiaceae | Chenopodioideae | ONLQ | *Atriplex* | *hortensis* | L. | ONLQ_Atriplex_hortensis | Paired | Non-stranded | 19485 | 0.46 | 10536 | 80.9% | used in Yang et. al 2015 | SRA | ERR2040208 /ERR2040209 | 1KP/Matasci et al., 2014 | Data access for the 1,000 Plants (1KP) project |
| Chenopodiaceae | Chenopodioideae | Atrlen | *Atriplex* | *lentiformis* | (Torr.) S. Watson | Atrlen_Atriplex_lentiformis | Paired | Non-stranded | 17454 | 0.5 | 10623 | 81.6% | --- | SRA | SRR1041720 | Li et al. 2015 | Understanding the biochemical basis of temperature-induced lipid pathway adjustments in plants. |
| Chenopodiaceae | Chenopodioideae | AAXJ | *Atriplex* | *prostrata* | Boucher ex DC. | AAXJ_Atriplex_prostrata | Paired | Non-stranded | 19161 | 0.43 | 10285 | 79.0% | used in Yang et. al 2015 | SRA | ERR2040210/ERR2040211 | 1KP/Matasci et al., 2014 | Data access for the 1,000 Plants (1KP) project |
| Chenopodiaceae | Chenopodioideae | CBJR | *Atriplex* | *rosea* | L. | CBJR_Atriplex_rosea | Paired | Non-stranded | 19991 | 0.45 | 10580 | 81.2% | used in Yang et. al 2015 | SRA | ERR2040212/ERR2040213 | 1KP/Matasci et al., 2014 | Data access for the 1,000 Plants (1KP) project |
| Chenopodiaceae | Chenopodioideae | MJM3295 | *Atriplex* | sp. | | MJM3295_Atriplex_sp | Paired | Stranded | 20391 | 0.5 | 10901 | 83.7% | used in Walker et al. 2018 | SRA | SRR6435344 | Walker et al. 2018 | From cacti to carnivores: Improved phylotranscriptomic sampling and hierarchical homology inference provide further insight into the evolution of Caryophyllales |
| Chenopodiaceae | Chenopodioideae | Chenbohen | *Blitum* | *bonus-henricus* | (L.) C.A. Mey. | Chenbohen_Chenopodium_bonus-henricus | Paired | Stranded | 27367 | 0.54 | 11337 | 87.0% | --- | Newly sequenced | Newly sequenced | Newly sequenced | Newly sequenced |
| Chenopodiaceae | Chenopodioideae | Cheama | *Chenopodium* | *amaranticolor* | (H.J. Coste & A. Reyn.) H.J. Coste & A. Reyn. | Cheama_Chenopodium_amaranticolor | Paired | Non-stranded | 33616 | 0.44 | 9915 | 76.1% | used in Yang et. al 2018 | SRA | SRR503600 | Zhang et al. 2012 | De novo foliar transcriptome of Chenopodium amaranticolor and analysis of its gene expression during virus-induced hypersensitive response |
| Chenopodiaceae | Chenopodioideae | Chpal | *Chenopodium* | *pallidicaule* | Aellen | Chpal_Chenopodium_pallidicaule | Paired | Non-stranded | 17132 | 0.51 | 3297 | 25.3% | --- | SRA | SRR4425240 | Jarvis et al. 2017 | The genome of Chenopodium quinoa |
| Chenopodiaceae | Chenopodioideae | Chquigen | *Chenopodium* | *quinoa* | Willd. | Chquigen_Chenopodium_quinoa | --- | --- | 44776 | --- | 11769 | 90.4% | --- | Phytozome | v1.0 | Jarvis et al. 2017 | The genome of Chenopodium quinoa |
| Chenopodiaceae | Chenopodioideae | Chsue | *Chenopodium* | *suecicum* | J. Murr | Chsue_Chenopodium_suecicum | Paired | Non-stranded | 17928 | 0.5 | 9503 | 73.0% | --- | SRA | SRR4425602 | Jarvis et al. 2017 | The genome of Chenopodium quinoa |
| Chenopodiaceae | Chenopodioideae | Chenvul | *Chenopodium* | *vulvaria* | L. | Chenvul_Chenopodium_vulvaria | Paired | Stranded | 16598 | 0.53 | 11153 | 85.6% | --- | Newly sequenced | Newly sequenced | Newly sequenced | Newly sequenced |
| Chenopodiaceae | Chenopodioideae | Dysamb | *Dysphania* | *ambrosioides* | (L.) Mosyakin & Clemants | Dysamb_Dysphania_ambrosia[a] | Paired | Stranded | 25436 | 0.54 | 11250 | 86.4% | --- | Newly sequenced | Newly sequenced | Newly sequenced | Newly sequenced |
| Chenopodiaceae | Chenopodioideae | Dyapha | *Dysphania* | *schraderiana* | (Schult.) Mosyakin & Clemants | Dyapha_Dysphania_schraderiana | Paired | Non-stranded | 14749 | 0.42 | 10070 | 77.3% | --- | SRA | SRR5989515 | --- | Submitted to SRA in 2017. No refrence found |
| Chenopodiaceae | Chenopodioideae | MJM3214 | *Extriplex* | *californica* | (Moq.) E.H. Zacharias | MJM3214_Extriplex_californica | Paired | Stranded | 16229 | 0.51 | 10966 | 84.2% | used in Yang et. al 2018 | SRA | SRR6787489 | Yang et al. 2018 | Improved transcriptome sampling pinpoints 26 ancient and more recent polyploidy events in Caryophyllales, including two allopolyploidy events |
| Chenopodiaceae | Chenopodioideae | Grabra | *Grayia* | *brandegeei* | A. Gray | Grabra_Grayia_brandegeei | Paired | Stranded | 15864 | 0.49 | 10554 | 81.0% | used in Walker et al. 2018 | SRA | SRR6435284 | Walker et al. 2018 | From cacti to carnivores: Improved phylotranscriptomic sampling and hierarchical homology inference provide further insight into the evolution of Caryophyllales |
| Chenopodiaceae | Chenopodioideae | MJM3268 | *Grayia* | *spinosa* | (Hook.) Moq. | MJM3268_Grayia_spinosa | Paired | Stranded | 16030 | 0.5 | 10664 | 81.9% | used in Yang et. al 2018 | SRA | SRR6787510 | Yang et al. 2018 | Improved transcriptome sampling pinpoints 26 ancient and more recent polyploidy events in Caryophyllales, including two allopolyploidy events |
| Chenopodiaceae | Chenopodioideae | MJM2311 | *Krascheninnikovia* | *lanata* | (Pursh) A. Meeuse & A. Smit | MJM2311_Krascheninnikovia_lanata | Paired | Stranded | 18996 | 0.53 | 11264 | 86.5% | used in Yang et. al 2018 | SRA | SRR6787516 | Yang et al. 2018 | Improved transcriptome sampling pinpoints 26 ancient and more recent polyploidy events in Caryophyllales, including two allopolyploidy events |
| Chenopodiaceae | Chenopodioideae | Oxru | *Oxybasis* | *rubra* | (L.) S. Fuentes-B., Uotila & Borsch | Oxru_Oxybasis_rubra | Paired | Non-stranded | 25776 | 0.53 | 11340 | 87.1% | used in Walker et al. 2018 | SRA | SRR2913184 | --- | Submitted to SRA in 2015. No refrence found |
| Chenopodiaceae | Chenopodioideae | MJM3031 | *Proatriplex* | *pleiantha* | (W.A. Weber) Stutz & G.L. Chu | MJM3031_Proatriplex_pleiantha | Paired | Stranded | 14808 | 0.49 | 10299 | 79.1% | used in Walker et al. 2018 | SRA | SRR6435312 | Walker et al. 2018 | From cacti to carnivores: Improved phylotranscriptomic sampling and hierarchical homology inference provide further insight into the evolution of Caryophyllales |
| Chenopodiaceae | Chenopodioideae | Spolegen | *Spinacia* | *oleracea* | L. | Spolegen_Spinacia_oleracea | --- | --- | 25495 | --- | 11586 | 89.0% | --- | Genome | v1.0 | Xu et al. 2017 | Draft genome of spinach and transcriptome diversity of 120 Spinaciaaccessions |
| Chenopodiaceae | Chenopodioideae | Spitet | *Spinacia* | *tetrandra* | Steven ex M.Bieb. | Spitet_Spinacia_tetrandra | Single | Non-stranded | 18802 | 0.29 | 8819 | 67.7% | used in Walker et al. 2018 | SRA | SRR1766331 | Xu et al. 2017 | Draft genome of spinach and transcriptome diversity of 120 Spinaciaaccessions |
| Chenopodiaceae | Chenopodioideae | Spitur | *Spinacia* | *turkestanica* | Iljin | Spitur_Spinacia_turkestanica | Single | Non-stranded | 9066 | 0.11 | 4256 | 32.7% | Species used in Walker et al. 2018, but a different SRA accession | SRA | SRR1766333 | Xu et al. 2017 | Draft genome of spinach and transcriptome diversity of 120 Spinaciaaccessions |
| Chenopodiaceae | Chenopodioideae | MJM3228 | *Stutzia* | *covillei* | (Standl.) E.H. Zacharias | MJM3228_Stutzia_covillei | Paired | Stranded | 14806 | 0.39 | 9807 | 75.3% | used in Walker et al. 2018 | SRA | SRR6435340 | Walker et al. 2018 | From cacti to carnivores: Improved phylotranscriptomic sampling and hierarchical homology inference provide further insight into the evolution of Caryophyllales |
| Chenopodiaceae | Corispermoideae | Agsq | *Agriophyllum* | *squarrosum* | (L.) Moq. | Agsq_Agriophyllum_squarrosum | Paired | Non-stranded | 19671 | 0.54 | 11211 | 86.1% | used in Yang et. al 2018 | SRA | SRR1559276 | Zhao et al. 2014 | Transcriptomic analysis of a psammophyte food crop, sand rice (Agriophyllum squarrosum) and identification of candidate genes essential for sand dune adaptation. |
| Chenopodiaceae | Corispermoideae | MSB0000170SFB | *Corispermum* | *hyssopifolium* | L. | MSB0000170SFB_Corispermum_hyssopifolium | Paired | Stranded | 16634 | 0.56 | 11409 | 87.6% | used in Walker et al. 2018 | SRA | SRR6435347 | Walker et al. 2018 | From cacti to carnivores: Improved phylotranscriptomic sampling and hierarchical homology inference provide further insight into the evolution of Caryophyllales |
| Chenopodiaceae | Salicornioideae | SalspSFB | *Allenrolfea* | sp. | | SalspSFB_Allenrolfea_sp | Paired | Stranded | 16123 | 0.51 | 10716 | 82.3% | used in Walker et al. 2018 | SRA | SRR6435351 | Walker et al. 2018 | From cacti to carnivores: Improved phylotranscriptomic sampling and hierarchical homology inference provide further insight into the evolution of Caryophyllales |
| Chenopodiaceae | Salicornioideae | ArmacSFB | *Arthrocaulon* | *macrostachyum* | (Moric.) Piirainen & G.Kadereit | ArmacSFB_Arthrocnemum_macrostachyum[a] | Paired | Stranded | 18895 | 0.5 | 10634 | 81.6% | used in Walker et al. 2018 | SRA | SRR6435310 | Walker et al. 2018 | From cacti to carnivores: Improved phylotranscriptomic sampling and hierarchical homology inference provide further insight into the evolution of Caryophyllales |
| Chenopodiaceae | Salicornioideae | Halcas | *Halostachys* | *caspica* | C.A. Mey. ex Schrenk | Halcas_Halostachys_caspica | Paired | Non-stranded | 33573 | 0.49 | 10563 | 81.1% | --- | SRA | SRR7003638 | --- | Submitted to SRA in 2018. No reference found |
| Chenopodiaceae | Salicornioideae | Heteros | *Heterostachys* | *olivascens* | (Speg.) Molfino | Heteros_Heterostachys_sp[a] | Paired | Stranded | 18385 | 0.47 | 10397 | 79.8% | --- | Newly sequenced | Newly sequenced | Newly sequenced | Newly sequenced |
| Chenopodiaceae | Salicornioideae | KalcuSFB | *Kalidium* | *cuspidatum* | (L.) Ung.-Sternb. | KalcuSFB_Kalidium_cuspidatum | Paired | Stranded | 18710 | 0.49 | 10190 | 78.2% | used in Walker et al. 2018 | SRA | SRR6435350 | Walker et al. 2018 | From cacti to carnivores: Improved phylotranscriptomic sampling and hierarchical homology inference provide further insight into the evolution of Caryophyllales |
| Chenopodiaceae | Salicornioideae | Saeu2 | *Salicornia* | *europaea* | L. | Saeu2_Salicornia_europaea | Paired | Non-stranded | 31052 | 0.37 | 9809 | 75.3% | Species used in Walker et al. 2018, but a different SRA library | SRA | SRR823398 | Ma et al. 2013 | Global transcriptome profiling of Salicornia europaea L. shoots under NaCl treatment |
| Chenopodiaceae | Salicornioideae | Salipac | *Salicornia* | *pacifica* | Standl. | Salipac_Salicornia_pacifica | Paired | Stranded | 30611 | 0.53 | 10891 | 83.6% | used in Walker et al. 2018 | SRA | SRR6435339 | Walker et al. 2018 | From cacti to carnivores: Improved phylotranscriptomic sampling and hierarchical homology inference provide further insight into the evolution of Caryophyllales |
| Chenopodiaceae | Salicornioideae | TecpeSFB | *Tecticornia* | *pergranulata* | (L.) Dumort. | TecpeSFB_Tecticornia_pergranulata | Paired | Stranded | 26784 | 0.53 | 10898 | 83.7% | used in Walker et al. 2018 | SRA | SRR6435313 | Walker et al. 2018 | From cacti to carnivores: Improved phylotranscriptomic sampling and hierarchical homology inference provide further insight into the evolution of Caryophyllales |
| Chenopodiaceae | Salsoloidae | AnaarSFB | *Anabasis* | *articulata* | Moq. | AnaarSFB_Anabasis_articulata | Paired | Stranded | 21316 | 0.52 | 10112 | 77.6% | used in Walker et al. 2018 | SRA | SRR6435311 | Walker et al. 2018 | From cacti to carnivores: Improved phylotranscriptomic sampling and hierarchical homology inference provide further insight into the evolution of Caryophyllales |
| Chenopodiaceae | Salsoloidae | CarveSFB | *Caroxylon* | *vermiculatum* | (L.) Akhani & Roalson | CarveSFB_Caroxylon_vermiculatum | Paired | Stranded | 58480 | 0.55 | 11214 | 86.1% | used in Walker et al. 2018 | SRA | SRR6435345 | Walker et al. 2018 | From cacti to carnivores: Improved phylotranscriptomic sampling and hierarchical homology inference provide further insight into the evolution of Caryophyllales |
| Chenopodiaceae | Salsoloidae | Hagl | *Halogeton* | *glomeratus* | (M. Bieb.) C.A. Mey. | Hagl_Halogeton_glomeratus | Paired | Non-stranded | 24367 | 0.34 | 9248 | 71.0% | used in Walker et al. 2018 | SRA | SRR1503502 | Yao et al. 2018 | Transcriptome sequencing and comparative analysis of differentially-expressed isoforms in the roots of Halogeton glomeratus under salt stress |
| Chenopodiaceae | Salsoloidae | Haam | *Haloxylon* | *ammodendron* | Bunge | Haam_Haloxylon_ammodendron | Paired | Non-stranded | 29314 | 0.55 | 10422 | 80.0% | used in Yang et. al 2018 | SRA | SRR1697346 | Long et al. 2014 | *De novo* assembly of the desert tree Haloxylon ammodendron (C. A. Mey.) based on RNA-Seq data provides insight into drought response, gene discovery and marker identification. |
| Chenopodiaceae | Salsoloidae | Hamsco | *Hammada* | *scoparia* | Iljin | Hamsco_Hammada_scoparia | Single | Non-stranded | 23086 | 0.49 | 9787 | 75.1% | --- | SRA | ERR2060287 | Lauterbach et al. 2017 | *De novo Transcriptome Assembly and Comparison of C3, C3-C4, and C4Species of Tribe Salsoleae (Chenopodiaceae)* |
| Chenopodiaceae | Salsoloidae | MSB0496298SFB | *Kali* | *collina* | Akhani & Roalson | MSB0496298SFB_Kali_collina | Paired | Stranded | 18660 | 0.55 | 10955 | 84.1% | used in Walker et al. 2018 | SRA | SRR6435349 | Walker et al. 2018 | From cacti to carnivores: Improved phylotranscriptomic sampling and hierarchical homology inference provide further insight into the evolution of Caryophyllales |
| Chenopodiaceae | Salsoloidae | Salopp | *Salsola* | *oppositifolia* | Desf. | Salopp_Salsola_oppositifolia | Single | Non-stranded | 48370 | 0.49 | 10448 | 80.2% | --- | SRA | ERR2060302 | Lauterbach et al. 2017 | *De novo Transcriptome Assembly and Comparison of C3, C3-C4, and C4Species of Tribe Salsoleae (Chenopodiaceae)* |
| Chenopodiaceae | Salsoloidae | Salsod | *Salsola* | *soda* | L. | Salsod_Salsola_soda | Single | Non-stranded | 18064 | 0.49 | 9714 | 74.6% | --- | SRA | SRR3544552 | Lauterbach et al. 2017 | *De novo Transcriptome Assembly and Comparison of C3, C3-C4, and C4Species of Tribe Salsoleae (Chenopodiaceae)* |
| Chenopodiaceae | Salsoloidae | Salweb | *Salsola* | *webbii* | Moq. | Salweb_Salsola_webbii | Single | Non-stranded | 30337 | 0.49 | 10529 | 80.8% | --- | SRA | ERR2060308 | Lauterbach et al. 2017 | *De novo Transcriptome Assembly and Comparison of C3, C3-C4, and C4Species of Tribe Salsoleae (Chenopodiaceae)* |
| Chenopodiaceae | Suaedoideae | Suaara | *Suaeda* | *aralocaspica* | Freitag & Schutze | Suaara_Suaeda_aralocaspica | Paired | Non-stranded | 18279 | 0.38 | 9879 | 75.9% | --- | SRA | SRR3706496 | Wang et al. 2017 | Transcriptome assembly in Suaeda aralocaspica to reveal the distinct temporal gene/miRNA alterations between the dimorphic seeds during germination |
| Chenopodiaceae | Suaedoideae | Suadivar | *Suaeda* | *divaricata* | Moq. | Suadivar_Suaeda_divaricata | Paired | Stranded | 28206 | 0.5 | 10837 | 83.2% | --- | Newly sequenced | Newly sequenced | Newly sequenced | Newly sequenced |
| Chenopodiaceae | Suaedoideae | Sufr | *Suaeda* | *fruticosa* | Forssk. ex J.F. Gmel. | Sufr_Suaeda_fruticosa | Paired | Non-stranded | 12139 | 0.18 | 6473 | 49.7% | used in Yang et. al 2018 | SRA | SRR1947661 | Diray-Arce et al. 2015 | ranscriptome assembly, profiling and differential gene expression analysis of the halophyte Suaeda fruticosa provides insights into salt tolerance |
| Chenopodiaceae | Suaedoideae | Suagla | *Suaeda* | *glauca* | (Bunge) Bunge | Suagla_Suaeda_glauca | Paired | Non-stranded | 16780 | 0.49 | 10454 | 80.3% | --- | SRA | SRR2416214 | Jin et al. 2016 | Salt-Responsive Transcriptome Profiling of Suaeda glaucavia RNA Sequencing |
| Chenopodiaceae | Suaedoideae | Suaifni | *Suaeda* | *ifniensis* | Caball. ex Maire | Suaifni_Suaeda_ifniensis | Paired | Stranded | 17461 | 0.48 | 10477 | 80.4% | --- | Newly sequenced | Newly sequenced | Newly sequenced | Newly sequenced |
| Chenopodiaceae | Suaedoideae | MJM1679 | *Suaeda* | *linearis* | (Elliott) Moq. | MJM1679_Suaeda_linearis | Paired | Stranded | 33753 | 0.48 | 10648 | 81.8% | used in Yang et. al 2018 | SRA | SRR6787513 | Yang et al. 2017 | Improved transcriptome sampling pinpoints 26 ancient and more recent polyploidy events in Caryophyllales, including two allopolyploidy events |
| Chenopodiaceae | Suaedoideae | Suma | *Suaeda* | *maritima* | (L.) Dumort. | Suma_Suaeda_maritima | Paired | Non-stranded | 46819 | 0.56 | 10828 | 83.1% | used in Walker et al. 2018 | SRA | SRR3218589 | Gharat et al. 2016 | Transcriptome Analysis of the Response to NaCl in Suaeda maritima Provides an Insight into Salt Tolerance Mechanisms in Halophytes |
| Chenopodiaceae | Suaedoideae | Suasal | *Suaeda* | *salsa* | Pall. | Suasal_Suaeda_salsa | Paired | Non-stranded | 18734 | 0.46 | 9991 | 76.7% | --- | SRA | SRR5877201 | Xu et al. 2017 | Transcriptomic profiling of genes in matured dimorphic seeds of euhalophyte Suaeda salsa |
| Chenopodiaceae | Suaedoideae | Suavera | *Suaeda* | *vera* | Forssk. ex J.F. Gmel. | Suavera_Suaeda_vera | Paired | Stranded | 16827 | 0.48 | 10341 | 79.4% | --- | Newly sequenced | Newly sequenced | Newly sequenced | Newly sequenced |
| Chenopodiaceae | Suaedoideae | MJM3086 | *Suckleya* | *suckleyana* | (Torr.) Rydb. | MJM3086_Suckleya_suckleyana | Paired | Stranded | 15189 | 0.51 | 10908 | 83.8% | used in Walker et al. 2018 | SRA | SRR6435288 | Walker et al. 2018 | From cacti to carnivores: Improved phylotranscriptomic sampling and hierarchical homology inference provide further insight into the evolution of Caryophyllales |
| Achatocarpaceae - OUTGROUP | | MJM2704 | *Achatocarpus* | *gracilis* | H. Walter | MJM2704_Achatocarpus_gracilis | Paired | Stranded | 24544 | 0.48 | 10397 | 79.8% | used in Walker et al. 2018 | SRA | SRR6435362 | Walker et al. 2018 | From cacti to carnivores: Improved phylotranscriptomic sampling and hierarchical homology inference provide further insight into the evolution of Caryophyllales |
| Achatocarpaceae - OUTGROUP | | MJM1677 | *Phaulothamnus* | *spinescens* | A. Gray | MJM1677_Phaulothamnus_spinescens | Paired | Stranded | 28006 | 0.45 | 9714 | 74.6% | used in Yang et. al 2018 | SRA | SRR1979686 | Brockington et al 2015 | Lineage-specific gene radiations underlie the evolution of novel betalain pigmentation in Caryophyllales |
| Aizoaceae - OUTGROUP | | BJKT | *Delosperma* | *echinatum* | (Lam.) Schwantes | BJKT_Delosperma_echinatum | Paired | Non-stranded | 18194 | 0.26 | 7498 | 57.6% | used in Yang et. al 2018 | SRA | ERR2040192 | 1KP/Matasci et al., 2014 | Data access for the 1,000 Plants (1KP) project |
| Aizoaceae - OUTGROUP | | Mecrygen | *Mesembryanthemum* | *crystallinum* | L. | Mecrygen_Mesembryanthemum_crystallinum | --- | --- | 25435 | --- | 10488 | 80.5% | --- | Yim and Cushman, Unpublished data | --- | --- | Yim and Cushman, Unpublished data |
| Caryophyllaceae - OUTGROUP | | CorSFB | *Corrigiola* | *litoralis* | L. | CorSFB_Corrigiola_litoralis | Paired | Stranded | 19203 | 0.52 | 10577 | 81.2% | used in Yang et. al 2018 | SRA | SRR6787499 | Yang et al. 2018 | Improved transcriptome sampling pinpoints 26 ancient and more recent polyploidy events in Caryophyllales, including two allopolyploidy events. |
| Caryophyllaceae - OUTGROUP | | Dicargen | *Dianthus* | *caryophyllus* | L. | Dicargen_Dianthus_caryophyllus | --- | --- | 43264 | --- | 7938 | 60.9% | --- | http://carnation.kazusa.or.jp | DCA_r1.0 | Yagi et al. 2014 | Sequence analysis of the genome of carnation (Dianthus caryophyllus L.). |
| Caryophyllaceae - OUTGROUP | | HerSFB | *Herniaria* | *latifolia* | Lapeyr. | HerSFB_Herniaria_latifolia | Paired | Stranded | 38364 | 0.54 | 9901 | 76.0% | used in Yang et. al 2018 | SRA | SRR6787494 | Yang et al. 2018 | Improved transcriptome sampling pinpoints 26 ancient and more recent polyploidy events in Caryophyllales, including two allopolyploidy events. |
| Caryophyllaceae - OUTGROUP | | IlleSFB | *Illecebrum* | *verticillatum* | L. | IlleSFB_Illecebrum_verticillatum | Paired | Stranded | 22434 | 0.5 | 9666 | 74.2% | used in Yang et. al 2018 | SRA | SRR6787493 | Yang et al. 2018 | Improved transcriptome sampling pinpoints 26 ancient and more recent polyploidy events in Caryophyllales, including two allopolyploidy events. |
| Caryophyllaceae - OUTGROUP | | TJES | *Spergularia* | *media* | (L.) C. Presl | TJES_Spergularia_media | Paired | Non-stranded | 21020 | 0.28 | 7643 | 58.7% | used in Yang et. al 2015 | SRA | ERR2040231 | 1KP/Matasci et al., 2014 | Data access for the 1,000 Plants (1KP) project |
| Microteaceae - OUTGROUP | | YNFJ | *Microtea* | *debilis* | Sw. | YNFJ_Microtea_debilis | Paired | Non-stranded | 29481 | 0.34 | 8629 | 66.3% | used in Yang et. al 2015 | SRA | ERR2040255 | 1KP/Matasci et al., 2014 | Data access for the 1,000 Plants (1KP) project |
| Molluginaceae - OUTGROUP | | HURS | *Mollugo* | *pentaphylla* | L. | HURS_Mollugo_pentaphylla | Paired | Non-stranded | 16900 | 0.36 | 7925 | 60.8% | used in Yang et. al 2015 | SRA | ERR2040238 | 1KP/Matasci et al., 2014 | Data access for the 1,000 Plants (1KP) project |
| Nyctaginaceae - OUTGROUP | | MJM2726B | *Commicarpus* | *scandens* | (L.) Standl. | MJM2726B_Commicarpus_scandens | Paired | Stranded | 24282 | 0.48 | 8747 | 67.2% | used in Walker et al. 2018 | SRA | SRR6435360 | Walker et al. 2018 | From cacti to carnivores: Improved phylotranscriptomic sampling and hierarchical homology inference provide further insight into the evolution of Caryophyllales |
| Polygonaceae - OUTGROUP | | Fatatgen | *Fagopyrum* | *tataricum* | (L.) Gaertn. | Fatatgen_Fagopyrum_tataricum | --- | --- | 31834 | --- | 6120 | 47.0% | --- | http://www.mbkbase.org/Pinku1 | v2.0 | Zhang et al. 2017 | The Tartary Buckwheat Genome Provides Insights into Rutin Biosynthesis and Abiotic Stress Tolerance |

[a]Taxon names updated in ms figures after analyses.

**Table S2.** Voucher information of newly sequenced transcriptomes.

| Family | Subfamily | Species code | Genus | Species | Authority name | Voucher information[a] | Locality | Growth condition | Tissue | RNA extraction | RNA-seq library preparation | Raw read pairs | Filtered nuclear read pairs | Filtered organelle read pairs | SRA accession |
|---|---|---|---|---|---|---|---|---|---|---|---|---|---|---|---|
| Amaranthaceae s.l. | Betoideae | Betatrig | *Beta* | *vulgaris* | L. | Unvouchered | Cultivated at Botanical Garden University Mainz, orig. coll. Bulgaria. Living Collection Botanic Garden Mainz 63. | Cultivated at Botanical Garden University Mainz | Leaf | QIAGEN RNeasy Plant Mini Kit with in-column DNase digestion by Delphine Tefarikis at Kadereit Lab Jun 2018 | TruSeq Stranded Total RNA Library Prep Plant with RiboZero by U Minnesota Genomics Center Aug 2018 | 22,125,749 | 12,050,145 | 9,280,410 | SRR12121649 |
| Amaranthaceae s.l. | Betoideae | Habtam | *Hablitzia* | *tamnoides* | M. Bieb. | Kadereit, G. s.n. (MJG 027642) | Cultivated at Botanical Garden University Mainz, seeds obtained from a nursery | Cultivated at Botanical Garden University Mainz | Leaf | QIAGEN RNeasy Plant Mini Kit with in-column DNase digestion by Delphine Tefarikis at Kadereit Lab Jun 2018 | TruSeq Stranded Total RNA Library Prep Plant with RiboZero by U Minnesota Genomics Center Aug 2018 | 16,820,668 | 6,783,386 | 7,402,286 | SRR12121648 |
| Amaranthaceae s.l. | Betoideae | Patpat | *Patellifolia* | *patellaris* | (Moq.) A.J. Scott, Ford-Lloyd & J.T. Williams | H. Freitag 40031 (MJG 013582) | SW Morocco. Living Collection Botanic Garden Mainz 58. | Cultivated at Botanical Garden University Mainz | Leaf | QIAGEN RNeasy Plant Mini Kit with in-column DNase digestion by Delphine Tefarikis at Kadereit Lab Jun 2018 | TruSeq Stranded Total RNA Library Prep Plant with RiboZero by U Minnesota Genomics Center Aug 2018 | 20,112,651 | 12,691,402 | 6,396,902 | SRR12121640 |
| Amaranthaceae s.s | Celosioids | Deeamar | *Deeringia* | *amaranthoides* | (Lam.) Merr. | Millennium Seed Bank 301198 (MJG 023037) | Orig. coll. China, Yunnan. Living Collection Botanic Garden Mainz 344. | Cultivated at Botanical Garden University Mainz | Leaf | QIAGEN RNeasy Plant Mini Kit with in-column DNase digestion by Delphine Tefarikis at Kadereit Lab Jun 2018 | TruSeq Stranded Total RNA Library Prep Plant with RiboZero by U Minnesota Genomics Center Aug 2018 | 21,713,722 | 17,238,976 | 3,048,492 | SRR12121639 |
| Amaranthaceae s.s. | Celosioids | Hergla | *Hermbstaedtia* | *glauca* | Reichb. ex Steud. | Millennium Seed Bank 140498 (MJG 023036) | Orig. coll. South Africa, Cape Provinces. Living Collection Botanic Garden Mainz 310. | Cultivated at Botanical Garden University Mainz | Leaf | QIAGEN RNeasy Plant Mini Kit with in-column DNase digestion by Delphine Tefarikis at Kadereit Lab Jun 2018 | TruSeq Stranded Total RNA Library Prep Plant with RiboZero by U Minnesota Genomics Center Aug 2018 | 21,285,421 | 16,726,894 | 3,619,788 | SRR12121638 |
| Amaranthaceae s.s. | Gomphrenoids | Gomcel | *Gomphrena* | *celosioides* | Mart. | Aagesen, L. 41 (SI) | Argentina, Misiones, Leandro N. Alem. | Cultivated at Botanical Garden University Mainz | Leaf | QIAGEN RNeasy Plant Mini Kit with in-column DNase digestion by Delphine Tefarikis at Kadereit Lab Jun 2018 | TruSeq Stranded Total RNA Library Prep Plant with RiboZero by U Minnesota Genomics Center Aug 2018 | 21,432,362 | 14,924,209 | 5,402,969 | SRR12121636 |
| Amaranthaceae s.s. | Gomphrenoids | Gomele | *Gomphrena* | *elegans* | Mart. | R. Norte (MJG 027640) | Argentina, Buenos Aires, Isidro; Cultivada en el vivero de Ribera norte. | Cultivated at Botanical Garden University Mainz | Leaf | QIAGEN RNeasy Plant Mini Kit with in-column DNase digestion by Delphine Tefarikis at Kadereit Lab Jun 2018 | TruSeq Stranded Total RNA Library Prep Plant with RiboZero by U Minnesota Genomics Center Aug 2018 | 21,301,322 | 14,457,205 | 5,711,422 | SRR12121635 |
| Amaranthaceae s.s. | Gomphrenoids | Pfatub | *Pfaffia* | *tuberosa* | (Spreng.) Hicken | Aagesen, L. 51 (SI) | Argentina, Corrientes, Santo Tomé. | Cultivated at Botanical Garden University Mainz | Leaf | QIAGEN RNeasy Plant Mini Kit with in-column DNase digestion by Delphine Tefarikis at Kadereit Lab Jun 2018 | TruSeq Stranded Total RNA Library Prep Plant with RiboZero by U Minnesota Genomics Center Aug 2018 | 21,358,353 | 11,557,356 | 8,516,444 | SRR12121634 |
| Amaranthaceae s.s. | Gomphrenoids | Quaephe | *Quaternella* | *ephedroides* | Pedersen | Millennium Seed Bank 107437 (MJG 023038) | Orig. coll. Brazil, Bahia Palmeiras. Living Collection Botanic Garden Mainz 270. | Cultivated at Botanical Garden University Mainz | Leaf | QIAGEN RNeasy Plant Mini Kit with in-column DNase digestion by Delphine Tefarikis at Kadereit Lab Jun 2018 | TruSeq Stranded Total RNA Library Prep Plant with RiboZero by U Minnesota Genomics Center Aug 2018 | 21,181,847 | 11,913,784 | 8,293,073 | SRR12121637 |
| Amaranthaceae s.s. | Gomphrenoids | Tisp1013 | *Tidestromia* | *oblongifolia* | (S. Watson) Standl. | Bell 7656 | Orig. coll. Orocopia Mountains; southwestern section of the range; collecting in an unnamed canyon to the southeast of Orocopia Canyon; about 3.6 air miles northeast of Dos Palmas. Cultivated at California Botanic Garden. | Culitivated at California Botanic Garden | Leaf | QIAGEN RNeasy Plant Mini Kit and QIAGEN RNAse-Free DNAse Set by Alfonso Timoneda at Brockington lab. Cambridge Sep 2016 | KAPA Stranded RNA-Seq Library Preparation Kit (KK8420) with poly-A enrichment by Ning Wang at Smith Lab Oct 13, 2016 | 26,708,058 | 20,480,127 | 1,022,721 | SRR12121633 |
| Chenopodiaceae | Chenopodioideae | Chenbohen | *Blitum* | *bonus-henricus* | (L.) C.A Mey. | Kadereit, G. s.n. (MJG 027643) | Cultivated at Botanical Garden University Mainz. | Cultivated at Botanical Garden University Mainz | Leaf | QIAGEN RNeasy Plant Mini Kit with in-column DNase digestion by Delphine Tefarikis at Kadereit Lab Jun 2018 | TruSeq Stranded Total RNA Library Prep Plant with RiboZero by U Minnesota Genomics Center Aug 2018 | 20,383,537 | 13,403,745 | 5,683,389 | SRR12121647 |
| Chenopodiaceae | Chenopodioideae | Chenvul | *Chenopodium* | *vulvaria* | L. | Kadereit, G. s.n. (MJG 027658) | Cultivated at Botanical Garden University of Mainz. Seeds obtained from Botanical Garden Innsbruck (Austria). Living Collection Botanic Garden Mainz 241. | Cultivated at Botanical Garden University Mainz | Leaf, flower bud | QIAGEN RNeasy Plant Mini Kit and QIAGEN RNAse-Free DNAse Set by Alfonso Timoneda at Brockington lab. | TruSeq Stranded Total RNA Library Prep Plant with RiboZero by U Minnesota Genomics Center Aug 2018 | 21,559,488 | 11,800,777 | 7,538,649 | SRR12121646 |
| Chenopodiaceae | Chenopodioideae | Dysamb | *Dysphania* | *ambrosioides* | (L.) Mosyakin & Clemants | Millennium Seed Bank 57738 (MJG 022990) | Cultivated at the Botanical Garden Mainz, orig. coll. unknown. | Cultivated at Botanical Garden University Mainz | Leaf | QIAGEN RNeasy Plant Mini Kit with in-column DNase digestion by Delphine Tefarikis at Kadereit Lab Jun 2018 | TruSeq Stranded Total RNA Library Prep Plant with RiboZero by U Minnesota Genomics Center Aug 2018 | 21,362,486 | 14,081,344 | 5,073,626 | SRR12121644 |
| Chenopodiaceae | Salicornioideae | Heteros | *Heterostachys* | *olivascens* | (Speg.) Molfino | Kadereit, G. s.n. (MJG 027635) | Cultivated at the Botanical Garden Mainz, orig. coll. San Antonio del Oeste, Rio Negro, Argentina, source seed exchange. Living Collection Botanic Garden Mainz 34. | Cultivated at Botanical Garden University Mainz | Leaf | QIAGEN RNeasy Plant Mini Kit with in-column DNase digestion by Delphine Tefarikis at Kadereit Lab Jun 2018 | TruSeq Stranded Total RNA Library Prep Plant with RiboZero by U Minnesota Genomics Center Aug 2018 | 19,449,518 | 11,602,378 | 7,033,278 | SRR12121643 |
| Chenopodiaceae | Suaedoideae | Suadivar | *Suaeda* | *divaricata* | Moq. | Unvouchered | Cultivated at the Botanical Garden Mainz, orig. coll. Comodoro Rivadavia, Chubut, Argentina; source seed exchange. Living Collection Botanic Garden Mainz 35. | Cultivated at Botanical Garden University Mainz | Leaf and apical meristem | QIAGEN RNeasy Plant Mini Kit and QIAGEN RNAse-Free DNAse Set by Alfonso Timoneda at Brockington lab. | TruSeq Stranded Total RNA Library Prep Plant with RiboZero by U Minnesota Genomics Center Aug 2018 | 20,571,773 | 15,310,442 | 4,298,598 | SRR12121642 |
| Chenopodiaceae | Suaedoideae | Suaifni | *Suaeda* | *ifniensis* | Caball. ex Maire | H. Freitag 40022 (MJG 013577) | Cultivated at the Botanical Garden Mainz, orig. coll. SW Morocco, Merlift ca. 25 km behind NE Sidi Ifni. Living Collection Botanic Garden Mainz 205 (MJG 027655). | Cultivated at Botanical Garden University Mainz | Leaf and apical meristem | QIAGEN RNeasy Plant Mini Kit and QIAGEN RNAse-Free DNAse Set by Alfonso Timoneda at Brockington lab. | TruSeq Stranded Total RNA Library Prep Plant with RiboZero by U Minnesota Genomics Center Aug 2018 | 23,255,093 | 13,880,141 | 8,274,304 | SRR12121645 |
| Chenopodiaceae | Suaedoideae | Suavera | *Suaeda* | *vera* | Forssk. ex J.F. Gmel. | Kadereit, G. s.n. (MJG 027641) | Cultivated at Botanical Garden Mainz, seeds obtained from Botanical Garden Lisboa (Portugal). Living Collection Botanic Garden Mainz 107. | Cultivated at Botanical Garden University Mainz | Leaf and apical meristem | QIAGEN RNeasy Plant Mini Kit and QIAGEN RNAse-Free DNAse Set by Alfonso Timoneda at Brockington lab. | TruSeq Stranded Total RNA Library Prep Plant with RiboZero by U Minnesota Genomics Center Aug 2018 | 22,667,424 | 12,662,690 | 8,652,755 | SRR12121641 |

[a]Note: MJG = Johannes Gutenberg-Universität Herbarium; SI = Instituto de Botánica Darwinion Herbarium

**Table S3.** Chloroplast references used for plastome assembly and tree inference.

| **Family** | **Genus** | **Species** | **Authority name** | **Notes** | **Source database** | **Source code** | **Souce reference** | **Souce title** |
|---|---|---|---|---|---|---|---|---|
| Aizoaceae | *Mesembryanthemum* | *crystallinum* | L. | Used as reference and and tree inference - OUTGROUP | GenBank | NC_029049 | --- | Yim, Ha, and Cushman, 2016. Unpublished. |
| Amaranthaceae | *Amaranthus* | *hypochondriacus* | L. | Used as reference and and tree inference | GenBank | NC_030770 | Chaney et al. 2016 | The complete chloroplast genome sequences for four *Amaranthus* species (Amaranthaceae) |
| Betoideae | *Beta* | *vulgaris* | L. | Used as reference and and tree inference | GenBank | KR230391 | Stadermann et al. 2015 | SMRT sequencing only de novo assembly of the sugar beet (*Beta vulgaris* ) chloroplast genome |
| Caryophyllaceae | *Dianthus* | *caryophyllus* | L. | Used as reference and and tree inference - OUTGROUP | GenBank | MG989277 | Chen et al. 2018 | Structural characteristic and phylogenetic analysis of the complete chloroplast genome of *Dianthus caryophyllus* . |
| Chenopodiaceae | *Bienertia* | *sinuspersici* | Akhani | Used as reference and and tree inference | GenBank | KU726550 | Kim et al. 2016 | The complete chloroplast genome sequence of *Bienertia sinuspersici* |
| Chenopodiaceae | *Chenopodium* | *quinoa* | Willd. | Used as reference and and tree inference | GenBank | NC_034949 | Hong et al. 2017 | Complete Chloroplast Genome Sequences and Comparative Analysis of *Chenopodium quinoa* and *C. album* . |
| Chenopodiaceae | *Haloxylon* | *persicum* | Bunge ex Boiss. & Buhse | Used only as assembly reference | GenBank | NC_027669 | Dong et al. 2016 | Comparative analysis of the complete chloroplast genome sequences in psammophytic *Haloxylon* species (Amaranthaceae) |
| Chenopodiaceae | *Salicornia* | *europaea* | L. | Used only as assembly reference | GenBank | NC_027225 | --- | Ho et al., 2015. Unpublished. |
| Chenopodiaceae | *Spinacia* | *oleracea* | L. | Used as reference and and tree inference | GenBank | NC_002202 | Schmitz-Linneweber et al. 2001 | The plastid chromosome of spinach (*Spinacia oleracea* ): complete nucleotide sequence and gene organization |
| Chenopodiaceae | *Suaeda* | *malacosperma* | Hara | Used only as assembly reference | GenBank | MG813535 | Park et al. 2018 | The complete plastid genome of *Suaeda malacosperma* (Amaranthaceae/Chenopodiaceae), a vulnerable halophyte in coastal regions of Korea and Japan |
| Montiaceae | *Cistanthe* | *longiscapa* | (Barnéoud) Carolin ex Hershk. | Used only as assembly reference | GenBank | NC_035140 | Stroll et al. 2017 | Development of microsatellite markers and assembly of the plastid genome in *Cistanthe longiscapa* (Montiaceae) based on low-coverage whole genome sequencing |
| Polygonaceae | *Fagopyrum* | *tataricum* | (L.) Gaertn. | Used as reference and and tree inference - OUTGROUP | GenBank | NC_027161 | Cho et al. 2015 | Complete Chloroplast Genome Sequence of Tartary Buckwheat (*Fagopyrum tataricum* ) and Comparative Analysis with Common Buckwheat (*F. esculentum* ). |

**Table S4.** Assembled plastid CDS and alignment stats.

| Gene name | No. taxa | Alignment length | No. nucleotides | % missing data |
|---|---|---|---|---|
| *accD* | 74 | 1452 | 99245 | 7.6% |
| *atpA* | 100 | 1529 | 152424 | 0.3% |
| *atpB* | 93 | 1500 | 136630 | 2.1% |
| *atpE* | 81 | 408 | 32895 | 0.5% |
| *atpF* | 100 | 555 | 54953 | 1.0% |
| *atpH* | 100 | 246 | 24600 | 0.0% |
| *atpI* | 100 | 744 | 73977 | 0.6% |
| *ccsA* | 64 | 972 | 61080 | 1.8% |
| *cemA* | 93 | 690 | 63866 | 0.5% |
| *clpP* | 70 | 588 | 34555 | 16.0% |
| *infA* | 95 | 234 | 22196 | 0.2% |
| *matK* | 73 | 1588 | 102683 | 11.4% |
| *ndhA* | 92 | 1099 | 95991 | 5.1% |
| *ndhB* | 81 | 777 | 58130 | 7.6% |
| *ndhC* | 98 | 363 | 35574 | 0.0% |
| *ndhD* | 73 | 1505 | 103259 | 6.0% |
| *ndhE* | 62 | 306 | 18777 | 1.0% |
| *ndhF* | 51 | 2291 | 99017 | 15.3% |
| *ndhG* | 63 | 531 | 33326 | 0.4% |
| *ndhH* | 91 | 1182 | 106082 | 1.4% |
| *ndhI* | 76 | 513 | 38416 | 1.5% |
| *ndhJ* | 95 | 477 | 45315 | 0.0% |
| *ndhK* | 100 | 684 | 67804 | 0.9% |
| *petA* | 96 | 963 | 91872 | 0.6% |
| *petB* | 64 | 642 | 41072 | 0.0% |
| *petD* | 95 | 479 | 45372 | 0.3% |
| *petG* | 76 | 114 | 8664 | 0.0% |
| *petL* | 72 | 96 | 6896 | 0.2% |
| *petN* | 53 | 90 | 4770 | 0.0% |
| *psaA* | 101 | 2254 | 225017 | 1.2% |
| *psaB* | 100 | 2205 | 218615 | 0.9% |
| *psaC* | 59 | 246 | 14514 | 0.0% |
| *psaI* | 93 | 111 | 10323 | 0.0% |
| *psaJ* | 73 | 135 | 9844 | 0.1% |
| *psbA* | 80 | 1066 | 85280 | 0.0% |
| *psbB* | 100 | 1527 | 151853 | 0.6% |
| *psbC* | 103 | 1435 | 146143 | 1.1% |
| *psbD* | 101 | 1009 | 101607 | 0.3% |
| *psbE* | 94 | 252 | 23688 | 0.0% |
| *psbF* | 94 | 120 | 11280 | 0.0% |
| *psbH* | 77 | 222 | 17045 | 0.3% |
| *psbI* | 67 | 111 | 7417 | 0.3% |
| *psbJ* | 94 | 123 | 11562 | 0.0% |

| Gene name | No. taxa | Alignment length | No. nucleotides | % missing data |
|---|---|---|---|---|
| *psbK* | 67 | 180 | 12058 | 0.0% |
| *psbL* | 94 | 117 | 10971 | 0.2% |
| *psbM* | 56 | 105 | 5880 | 0.0% |
| *psbN* | 76 | 132 | 10032 | 0.0% |
| *psbT* | 98 | 108 | 10497 | 0.8% |
| *psbZ* | 81 | 189 | 15309 | 0.0% |
| *rbcL* | 96 | 1428 | 134885 | 1.6% |
| *rpl2* | 85 | 825 | 70105 | 0.0% |
| *rpl14* | 95 | 366 | 34769 | 0.0% |
| *rpl16* | 92 | 358 | 32868 | 0.2% |
| *rpl20* | 85 | 387 | 32821 | 0.2% |
| *rpl22* | 88 | 599 | 50296 | 4.6% |
| *rpl23* | 82 | 268 | 21845 | 0.6% |
| *rpl32*[a] | 23 | 174 | 3956 | 1.1% |
| *rpl33* | 67 | 201 | 13402 | 0.5% |
| *rpl36* | 99 | 114 | 11257 | 0.3% |
| *rpoA* | 99 | 994 | 97641 | 0.8% |
| *rpoB* | 71 | 3261 | 176067 | 24.0% |
| *rpoC1* | 70 | 2049 | 129381 | 9.8% |
| *rpoC2* | 74 | 4267 | 235989 | 25.3% |
| *rps2* | 95 | 711 | 66704 | 1.2% |
| *rps3* | 92 | 657 | 59822 | 1.0% |
| *rps4* | 88 | 606 | 53206 | 0.2% |
| *rps7* | 90 | 468 | 42069 | 0.1% |
| *rps8* | 95 | 405 | 38475 | 0.0% |
| *rps11* | 98 | 417 | 40855 | 0.0% |
| *rps12* | 77 | 372 | 21630 | 24.5% |
| *rps14* | 96 | 303 | 28948 | 0.5% |
| *rps15* | 87 | 273 | 23680 | 0.3% |
| *rps16* | 64 | 268 | 16793 | 2.1% |
| *rps18* | 74 | 306 | 22618 | 0.1% |
| *rps19* | 86 | 279 | 23640 | 1.5% |
| *ycf2*[a] | 46 | 6514 | 258235 | 13.8% |
| *ycf3* | 93 | 515 | 43410 | 9.4% |
| *ycf4* | 96 | 555 | 52771 | 1.0% |

[a]Gene excluded from phylogenetic analyses due to low taxon occupancy

**Table S5.** Model selection between maximum number of reticulations in species networks searches.

| **Topology** | **Maximum number of reticulations allowed** | **Number of inferred reticulations** | ln(*L*) | **Parameters** | **Number of loci** | **AICc** | **Δ**AICc | **BIC** | **Δ**BIC |
|---|---|---|---|---|---|---|---|---|---|
| Nuclear concatenated | NA | NA | -24486.331 | 19 | 4138 | 49048.847 | 20589.6635 | 49130.8939 | 20546.6229 |
| ASTRAL | NA | NA | -23448.397 | 19 | 4138 | 46972.9794 | 18513.7959 | 47055.0262 | 18470.7552 |
| cpDNA concatenated | NA | NA | -24568.333 | 19 | 4138 | 49212.8503 | 20753.6668 | 49294.8971 | 20710.6261 |
| Network 1 | 1 | 1 | -21177.791 | 21 | 4138 | 42439.8068 | 13980.6233 | 42530.4696 | 13946.1986 |
| Network 2 | 2 | 2 | -17275.625 | 23 | 4138 | 34643.5188 | 6184.33532 | 34742.7937 | 6158.52273 |
| Network 3 | 3 | 2 | -16741.991 | 23 | 4138 | 33576.2506 | 5117.06715 | 33675.5256 | 5091.25455 |
| Network 4 | 4 | 3 | -15415.8 | 25 | 4138 | 30931.9164 | 2472.73289 | 31039.7994 | 2455.52844 |
| **Network 5** | **5** | **5** | **-14171.38** | **29** | **4138** | **28459.1835** | **0** | **28584.271** | **0** |

**Table S6.** Model selection between quartet tree topologies and species networks. Trees correspond to each of the three possible quartet topologies where H0 is the ASTRAL quartet species tree. Networks correspond to the best three networks for searches with one hybridization event allowed.

| **Quartet[a]** | **Topology[b]** | **ln(*L*)** | **Parameters** | **Number of loci** | **AICc** | **ΔAICc** | **BIC** | **ΔBIC** |
|---|---|---|---|---|---|---|---|---|
| BC1A | H0 | -9014.809786 | 5 | 8258 | 18049.62684 | 24.73436754 | 18074.71426 | 14.70279692 |
| | H1 | -9072.456373 | 5 | 8258 | 18164.92002 | 140.0275408 | 18190.00743 | 129.9959702 |
| | H2 | -9073.888783 | 5 | 8258 | 18167.78484 | 142.8923611 | 18192.87225 | 132.8607905 |
| | **Net 1** | **-8998.43945** | **7** | **8258** | **18024.89248** | **0** | **18060.01146** | **0** |
| | Net 2 | -8998.439526 | 7 | 8258 | 18024.89263 | 0.000151947 | 18060.01162 | 0.000151947 |
| | Net 3 | -8998.441478 | 7 | 8258 | 18024.89653 | 0.004056302 | 18060.01552 | 0.004056302 |
| ABC2 | H0 | -8516.854413 | 5 | 7811 | 17053.71651 | 12.87079823 | 17078.52527 | 2.950887757 |
| | H1 | -8581.563051 | 5 | 7811 | 17183.13379 | 142.2880731 | 17207.94254 | 132.3681626 |
| | H2 | -8582.670875 | 5 | 7811 | 17185.34944 | 144.5037223 | 17210.15819 | 134.5838118 |
| | **Net 1** | **-8506.415681** | **7** | **7811** | **17040.84572** | **0** | **17075.57438** | **0** |
| | Net 2 | -8506.415769 | 7 | 7811 | 17040.84589 | 0.000176519 | 17075.57456 | 0.000176519 |
| | Net 3 | -8506.42071 | 7 | 7811 | 17040.85577 | 0.010057548 | 17075.58444 | 0.010057548 |
| BC1C2 | H0 | -9140.191425 | 5 | 8385 | 18300.39001 | 156.347016 | 18325.55385 | 146.2848258 |
| | H1 | -9201.981045 | 5 | 8385 | 18423.96925 | 279.9262567 | 18449.13309 | 269.8640665 |
| | H2 | -9214.405292 | 5 | 8385 | 18448.81775 | 304.7747517 | 18473.98158 | 294.7125615 |
| | **Net 1** | **-9058.014812** | **7** | **8385** | **18144.04299** | **0** | **18179.26902** | **0** |
| | Net 2 | -9058.019338 | 7 | 8385 | 18144.05205 | 0.009052497 | 18179.27807 | 0.009052497 |
| | Net 3 | -9058.024046 | 7 | 8385 | 18144.06146 | 0.018468011 | 18179.28749 | 0.018468011 |
| C1PA | **H0** | **-8932.927759** | **5** | **8134** | **17885.8629** | **0** | **17910.87456** | **0** |
| | H1 | -8936.145955 | 5 | 8134 | 17892.29929 | 6.436391285 | 17917.31095 | 6.436391285 |
| | H2 | -8936.481125 | 5 | 8134 | 17892.96963 | 7.106730999 | 17917.98129 | 7.106730999 |
| | Net 1 | -8932.077808 | 7 | 8134 | 17892.1694 | 6.306498884 | 17927.18227 | 16.30771403 |
| | Net 2 | -8932.078011 | 7 | 8134 | 17892.16981 | 6.306905172 | 17927.18268 | 16.30812032 |
| | Net 3 | -8932.078714 | 7 | 8134 | 17892.17121 | 6.308310587 | 17927.18408 | 16.30952573 |
| PAC2 | H0 | -8530.661274 | 5 | 7784 | 17081.33026 | 40.10000797 | 17106.12168 | 30.18704595 |
| | H1 | -8552.9448 | 5 | 7784 | 17125.89731 | 84.66706025 | 17150.68873 | 74.75409823 |

| | | | | | | | | |
|---|---|---|---|---|---|---|---|---|
| | H2 | -8548.291438 | 5 | 7784 | 17116.59059 | 75.36033576 | 17141.382 | 65.44737374 |
| | **Net 1** | **-8506.607925** | **7** | **7784** | **17041.23025** | **0** | **17075.93463** | **0** |
| | Net 2 | -8506.609795 | 7 | 7784 | 17041.23399 | 0.00373969 | 17075.93837 | 0.00373969 |
| | Net 3 | -8506.618966 | 7 | 7784 | 17041.25233 | 0.02208072 | 17075.95671 | 0.02208072 |
| C1C2P | H0 | -9119.250871 | 5 | 8341 | 18258.50894 | 12.50997925 | 18283.64643 | 2.458344441 |
| | H1 | -9163.685997 | 5 | 8341 | 18347.37919 | 101.38023 | 18372.51669 | 91.32859519 |
| | H2 | -9164.83263 | 5 | 8341 | 18349.67246 | 103.6734974 | 18374.80995 | 93.62186263 |
| | **Net 1** | **-9108.992761** | **7** | **8341** | **18245.99896** | **0** | **18281.18809** | **0** |
| | Net 2 | -9108.994383 | 7 | 8341 | 18246.00221 | 0.003244509 | 18281.19133 | 0.003244509 |
| | Net 3 | -9108.994843 | 7 | 8341 | 18246.00313 | 0.0041636 | 18281.19225 | 0.0041636 |
| C1C2A | H0 | -8447.623029 | 5 | 7756 | 16915.2538 | 63.6063012 | 16940.02717 | 53.70057058 |
| | H1 | -8520.509174 | 5 | 7756 | 17061.02609 | 209.378593 | 17085.79946 | 199.4728624 |
| | H2 | -8522.764578 | 5 | 7756 | 17065.5369 | 213.889401 | 17090.31027 | 203.9836704 |
| | **Net 1** | **-8411.816521** | **7** | **7756** | **16851.6475** | **0** | **16886.3266** | **0** |
| | Net 2 | -8411.819912 | 7 | 7756 | 16851.65428 | 0.006781956 | 16886.33338 | 0.006781956 |
| | Net 3 | -8411.820308 | 7 | 7756 | 16851.65507 | 0.007573446 | 16886.33417 | 0.007573446 |
| ABP | H0 | -9008.115816 | 5 | 8206 | 18036.23895 | 3.307596079 | 18061.29474 | -6.711300872 |
| | H1 | -9015.941176 | 5 | 8206 | 18051.88967 | 18.95831519 | 18076.94546 | 8.939418238 |
| | H2 | -9014.738462 | 5 | 8206 | 18049.48424 | 16.55288764 | 18074.54003 | 6.533990688 |
| | **Net 1** | **-9002.458846** | **7** | **8206** | **18032.93135** | **0** | **18068.00604** | **0** |
| | Net 2 | -9002.460142 | 7 | 8206 | 18032.93395 | 0.002592568 | 18068.00863 | 0.002592568 |
| | Net 3 | -9002.464397 | 7 | 8206 | 18032.94246 | 0.011102577 | 18068.01714 | 0.011102577 |
| C1BP | **H0** | **-9557.910518** | **5** | **8793** | **19135.82787** | **0** | **19161.22959** | **0** |
| | H1 | -9661.475396 | 5 | 8793 | 19342.95762 | 207.1297559 | 19368.35935 | 207.1297559 |
| | H2 | -9661.009687 | 5 | 8793 | 19342.0262 | 206.1983365 | 19367.42793 | 206.1983365 |
| | Net 1 | -9556.24034 | 7 | 8793 | 19140.49343 | 4.665563813 | 19176.05266 | 14.82306554 |
| | Net 2 | -9556.243036 | 7 | 8793 | 19140.49882 | 4.670955519 | 19176.05805 | 14.82845724 |
| | Net 3 | -9556.246261 | 7 | 8793 | 19140.50527 | 4.677405326 | 19176.0645 | 14.83490705 |
| PBC2 | **H0** | **-9158.309463** | **5** | **8379** | **18336.62609** | **0** | **18361.78635** | **0** |
| | H1 | -9206.127177 | 5 | 8379 | 18432.26152 | 95.63542753 | 18457.42177 | 95.63542753 |
| | H2 | -9205.933131 | 5 | 8379 | 18431.87343 | 95.24733612 | 18457.03368 | 95.24733612 |

| | | | | | | | |
|---|---|---|---|---|---|---|---|
| Net 1 | -9158.016519 | 7 | 8379 | 18344.04642 | 7.42032489 | 18379.26742 | 17.48107897 |
| Net 2 | -9158.017286 | 7 | 8379 | 18344.04795 | 7.421858749 | 18379.26896 | 17.48261282 |
| Net 3 | -9158.017377 | 7 | 8379 | 18344.04813 | 7.422042036 | 18379.26914 | 17.48279611 |

[a]Each quartet is named following the species tree topology, where the first two are sister. A = Amaranthaceae. s.s. (*Amaranthus hypochondriacus* ), B = Betoideae (*Beta vulgaris* ), C1 = Chenopods I (*Chenopodium quinoa* ), C2 = Chenopods II (*Caroxylum vermiculatum* ), P = Polycnemoideae (*Polycnemum majus* ).

[b]All quartet tree topologies can be found in Figure 5 and quartet network topologies in Figure S9.

**Table S7.** Gene count based on raw likelihood, **Δ**AIC, and AU topology test. Trees correspond to each of the tree posible quartet topologies[a] where

| | | Raw bipartitions | | | Bipartition with bootstrap >= 50 | | | Raw likelihood | | | **Δ**AIC | | | AU topology test p-value | | | |
|---|---|---|---|---|---|---|---|---|---|---|---|---|---|---|---|---|---|
| **Quartet[b]** | **Number of loci** | **H0** | **H1** | **H2** | **H1** | **H2** | **H0** | **H0** | **H1** | **H2** | **H0** | **H1** | **H2** | **H0** | **H1** | **H2** | **Equivocal** |
| BC1A | 8258 | 3218 | 2723 | 2317 | 2753 | 2243 | 1912 | 3268 | 2692 | 2298 | 2130 | 1636 | 1295 | 130 | 98 | 47 | 7017 |
| ABC2 | 7811 | 3083 | 2521 | 2207 | 2666 | 2100 | 1804 | 3108 | 2502 | 2201 | 2085 | 1415 | 1252 | 204 | 68 | 47 | 6605 |
| BC1C2 | 8385 | 3319 | 2988 | 2078 | 2819 | 2516 | 1682 | 3365 | 2969 | 2051 | 2228 | 1817 | 1133 | 204 | 127 | 49 | 6945 |
| C1PA | 8134 | 2819 | 2705 | 2610 | 2407 | 2241 | 2173 | 2850 | 2684 | 2600 | 1822 | 1619 | 1592 | 95 | 80 | 98 | 6925 |
| PAC2 | 7784 | 2866 | 2216 | 2702 | 2444 | 1837 | 2286 | 2896 | 2210 | 2678 | 1876 | 1255 | 1605 | 120 | 67 | 96 | 6603 |
| C1C2P | 8341 | 3190 | 2738 | 2413 | 2741 | 2346 | 1982 | 3232 | 2720 | 2389 | 2091 | 1655 | 1378 | 173 | 97 | 88 | 6935 |
| C1C2A | 7756 | 3095 | 2619 | 2042 | 2679 | 2193 | 1670 | 3132 | 2588 | 2036 | 2054 | 1624 | 1136 | 153 | 107 | 53 | 6466 |
| ABP | 8206 | 2897 | 2532 | 2777 | 2444 | 2061 | 2320 | 2932 | 2526 | 2748 | 1859 | 1434 | 1583 | 134 | 56 | 52 | 6697 |
| C1BP | 8793 | 3573 | 2544 | 2676 | 3085 | 2070 | 2212 | 3609 | 2542 | 2642 | 2397 | 1433 | 1555 | 162 | 55 | 71 | 7429 |
| PBC2 | 8379 | 3216 | 2554 | 2609 | 2764 | 2073 | 2160 | 3257 | 2527 | 2595 | 2136 | 1454 | 1523 | 165 | 68 | 90 | 7062 |

[a]All topologies can be found in Figure 5.

[b]Each quartet is named following the species tree topology, where the first two are sister. A = Amaranthaceae. s.s. (*Amaranthus hypochondriacus* ), B = Betoideae (*Beta vulgaris* ), C1 = Chenopods I (*Chenopodium quinoa* ), C2 = Chenopods II (*Caroxylum vermiculatum* ), P = Polycnemoideae (*Polycnemum majus* ).

**Table S8.** Linear model results for correlation between quartet Tree consistency (TC) scores and alignment

| Quartet[a] | Sample size | Alignment length | | Gap content | | GC content | |
|---|---|---|---|---|---|---|---|
| | | R2 | p-value | R2 | p-value | R2 | p-value |
| BC1A | 8258 | 0.001147 | 0.002079 | 0.002228 | 1.78E-05 | 0.001008 | 0.003914 |
| ABC2 | 7811 | 3.76E-05 | 0.5882 | 0.001275 | 0.001599 | 0.0006525 | 0.02397 |
| BC1C2 | 8385 | 0.001113 | 0.002251 | 0.002348 | 9.02E-06 | 1.46E-04 | 0.2695 |
| C1PA | 8134 | 1.95E-04 | 0.2083 | 0.003885 | 1.85E-08 | 0.001725 | 0.0001788 |
| PAC2 | 7784 | 2.06E-03 | 6.26E-05 | 0.008132 | 1.57E-15 | 0.001579 | 0.0004539 |
| C1C2P | 8341 | 0.002681 | 2.23E-06 | 0.005697 | 5.12E-12 | 0.0006883 | 0.01657 |
| C1C2A | 7756 | 0.0007874 | 1.35E-02 | 0.005023 | 4.14E-10 | 0.001103 | 0.003436 |
| ABP | 8206 | 0.0001394 | 0.2849 | 0.0008305 | 0.009037 | 0.0003169 | 0.1068 |
| C1BP | 8793 | 0.001477 | 0.0003123 | 0.003134 | 1.50E-07 | 0.001401 | 0.0004478 |
| PBC2 | 8379 | 0.000225 | 0.1698 | 0.003343 | 1.18E-07 | 0.0005339 | 0.03442 |

[a]Each quartet is named following the species tree topology, where the first two are sister. A = Amaranthaceae. s.s. (*Amaranthus hypochondriacus* ), B = Betoideae (*Beta vulgaris* ), C1 = Chenopods I (*Chenopodium quinoa* ), C2 = Chenopods II (*Caroxylum vermiculatum* ), P = Polycnemoideae (*Polycnemum majus* ).

**Table S9.** ABBA/BABA test results of Amaranthaceae s.l. five main groups quartets.

| Quartet (H0)[a] | Number of loci | Sites in alignment | ABBA | BABA | Raw D-statistic | Z-score | P-value | Introgression direction |
|---|---|---|---|---|---|---|---|---|
| BC1A[b] | 8258 | 12778649 | 287226 | 254617 | 0.06018164 | 41.1085 | ≤ 0.001 | A⇔C1 |
| ABC2 | 7811 | 12105324 | 252772 | 376755 | -0.1969463 | 124.4161 | ≤ 0.001 | A⇔C2 |
| BC1C2 | 8385 | 13192317 | 306570 | 258349 | 0.08535914 | 54.59751 | ≤ 0.001 | C1⇔C2 |
| C1PA[b] | 8134 | 12635201 | 342350 | 286813 | 0.08827124 | 64.62297 | ≤ 0.001 | A⇔P |
| PAC2 | 7784 | 12049734 | 344726 | 405627 | -0.08116313 | 42.88069 | ≤ 0.001 | C2⇔P |
| C1C2P[b] | 8341 | 13127397 | 445384 | 276652 | 0.2336892 | 136.0151 | ≤ 0.001 | C2⇔P |
| C1C2A[b] | 7756 | 12114778 | 396219 | 292561 | 0.1504951 | 101.3243 | ≤ 0.001 | A⇔C2 |
| ABP | 8206 | 12622625 | 276319 | 312060 | -0.06074486 | 36.64264 | ≤ 0.001 | A⇔P |
| C1BP[b] | 8793 | 13712853 | 273286 | 261620 | 0.02180944 | 18.08364 | ≤ 0.001 | B⇔P |
| PBC2 | 8379 | 13074019 | 217549 | 415616 | -0.3128205 | 196.8972 | ≤ 0.001 | C2⇔P |

[a]Each quartet is named following the species tree topology, where the first two are sister. A = Amaranthaceae. s.s. (*Amaranthus hypochondriacus* ), B = Betoideae (*Beta vulgaris* ), C1 = Chenopods I (*Chenopodium quinoa* ), C2 = Chenopods II (*Caroxylum vermiculatum* ), P = Polycnemoideae (*Polycnemum majus* ). H0 topologies can be found in Figure 6

[b]Quartet compatible with the complete 105-taxon species trees

**Table S10.** Anomaly zone limit calculations in 11-taxon species trees. Bold rows show pair of internodes in the anomaly zone when y < a(x).

| Species tree[a] | Clade (x)[b] | Clade (y)[b] | x | y | a(x) |
|---|---|---|---|---|---|
| No partition | (C1, C2) | (C1) | 0.1467 | 2.722 | 0.1799 |
| | (C1, C2) | (C2) | 0.1467 | 2.1102 | 0.1799 |
| | **((C1, C2), B)** | **(C1, C2)** | **0.1045** | **0.1467** | **0.3084** |
| | ((C1, C2), B) | (B) | 0.1045 | 2.6081 | 0.3084 |
| | **(((C1, C2), B), P)** | **((C1, C2), B)** | **0.0846** | **0.1045** | **0.4003** |
| | (((C1, C2), B), P) | (P) | 0.0846 | 3.5424 | 0.4003 |

[a]Species tree topology can be found in Figure 6.

[b] B = Betoideae (*Beta vulgaris* ), C1 = Chenopods I (*Chenopodium quinoa* ), C2 = Chenopods II (*Caroxylum vermiculatum* ), P = Polycnemoideae (*Polycnemum majus* ).
